# Whole-genome sequencing reveals the molecular basis of sex determination in the dioecious wild yam *Dioscorea tokoro*

**DOI:** 10.1101/2025.04.28.650915

**Authors:** Aoi Kudoh, Satoshi Natsume, Yu Sugihara, Hiroaki Kato, Akira Abe, Kaori Oikawa, Motoki Shimizu, Kazue Itoh, Mai Tsujimura, Yoshitaka Takano, Toshiyuki Sakai, Hiroaki Adachi, Atsushi Ohta, Mina Ohtsu, Takuma Ishizaki, Toru Terachi, Ryohei Terauchi

## Abstract

Dioecious plants, which have distinct male and female individuals, constitute ∼5% of angiosperm species and have emerged frequently and independently from hermaphroditic ancestors. Although recent molecular studies of sex determination have started to reveal the diversity of the genetic systems underlying dioecy, research on the evolution of dioecy is limited, especially in monocots. Here, we describe the molecular basis of sex determination in the monocot *Dioscorea tokoro*, a dioecious wild yam endemic to East Asia. Chromosome-scale and haplotype-resolved genome assemblies and linkage analysis suggested that this plant has a male heterogametic sex determination (*XY*) system, with sex determination regions located on chromosome 3. Sequence read coverage analysis of the sex chromosomes revealed *X*- and *Y*-specific regions in putative pericentromeric chromosome regions. Within the *Y*-specific region, we identified two candidate genes that are likely involved in sex determination: *BLH9*, encoding a homeobox protein, and *HSP90*, encoding a molecular chaperone. BLH9 has similar functions to AtBLH9 in *Arabidopsis thaliana*. BLH9 is thought to suppress female organ development, whereas HSP90 might be required for pollen development. These results shed light on the complex evolution of dioecy in plants.

**Author summary:** Sexual reproduction is a nearly universal, indispensable feature of evolution in eukaryotes. The molecular mechanisms underlying sex determination vary depending on the taxon. However, most information about this process was derived from studies of model organisms and/or domesticated species. To elucidate the diversity and evolution of sex determination, we need to expand the taxonomic breadth of these investigations. Here, we focused on the monocot genus *Dioscorea*, which contains species with multiple sex-determination systems, suggesting that frequent evolutionary transitions occur between the different sex-determination systems. We investigated the genetics of sex determination in *Dioscorea tokoro*, a dioecious wild yam endemic to East Asia. Whole-genome assembly and genetic analysis, along with transcriptome analysis, suggested that this species employs a male heterogametic sex determination (*XY*) system, with two *Y*-chromosome-specific genes that might be involved in male and female differentiation. These findings enhance our understanding of the complex evolution of dioecy.

## Introduction

Dioecy is a mating system in which male and female flowers are borne on separate individuals. This system enforces outcrossing and enables efficient pollen and seed production due to the specialization of male or female functions [1–3]. Approximately 30,000 dioecious plant species have been identified, representing 5–10% of angiosperm species. The sex phenotype of dioecious plant species is thought to be genetically controlled [4–6]. The wide and scattered taxonomic distribution of dioecious plants points to the frequent independent emergence of genetic systems controlling the male and female traits [7–9].

The genetic control of sex phenotype is usually based on the presence of male or female heterogametic sex chromosomes (*XY*/*XX* or *ZZ*/*ZW*). The *Y* or *W* chromosome is generally thought to carry one or two genetic components for male or female sex determination; however, some plants employ an *X*-autosome balance system, in which sex is determined based on the ratio of sex chromosomes to autosomes [3,10–12]. In the lotus persimmon (*Diospyros lotus*) [13–15], *Oppressor of meGI* (*OGI*) on the *Y* chromosome encodes a small RNA that silences *Male Growth Inhibitor* (*MeGI*) and thereby promotes male development. Heterologous expression of *MeGI* in *Arabidopsis thaliana* and *Nicotiana tabacum* inhibits anther development, resulting in female flower development. Thus, in this plant, sex appears to be determined by a single genetic switch, *OGI* [13–16]. In garden asparagus (*Asparagus officinalis*), two genes on the *Y* chromosome, *SUPPRESSOR OF FEMALE FUNCTION* (*SOFF*) and *TAPETAL DEVELOPMENT AND FUNCTION1* (*aspTDF1*), function independently to suppress pistil development and promote anther development, respectively [17–20]. The genetic basis of sex determination has been revealed or predicted in ten other angiosperm genera. Variable genetic mechanisms for sex determination have been reported for different genera and even for different species within the same genus [6,12,21,22]. To understand the evolution of dioecy, the molecular basis of sex determination in a wide range of dioecious genera must be elucidated.

*Dioscorea* is the largest genus in the monocot family Dioscoreaceae and consists of approximately 630 species [23]. Most *Dioscorea* species are perennial herbaceous climbers that are widely distributed in tropical and temperate regions [24–26]. This genus includes the important tuber crop yam; reference genome sequences are available for four yam species [27–31]. Most *Dioscorea* species are dioecious, and multiple sex-determination systems have been proposed for this genus based on cytological observations and genetic linkage analyses: a male heterogametic sex-determination system *XY*/*XX* in *D. tokoro, Dioscorea gracillima, Dioscorea bulbifera, Dioscorea dumetorum, Dioscorea tomentosa, Dioscorea pentaphylla, Dioscorea spinosa*, *Dioscorea alata*, and *Dioscorea japonica* [32–36]; a female heterogametic sex-determination system *ZZ*/*ZW* in *Dioscorea deltoidea* and *Dioscorea rotundata* [27,37]; extra chromosomes in *XO*/*XX* females in *Dioscorea sinuata* and *Dioscorea reticulata* [38,39]; and tetraploid sex chromosomes with a male heterogametic (*XXYY* or *XXXY*) and female homogametic (*XXXX*) system in *Dioscorea floribunda* [40]. Different sex-determination systems are distributed across different sections of the genus. For example, the sections Stenophora and Enantiophyllum each contain species with *XY*/*XX* as well as *ZZ*/*ZW* systems [41,42], suggesting that genomic regions involved in sex determination arise via evolutionary transitions or independent evolution.

Recent genome analyses revealed genomic regions associated with sex phenotypes in *D. rotundata* and *D. alata*. Sex-linked genomic regions of *D. rotundata* were identified by QTL-seq analysis of F_1_ progeny segregating for male and female plants [27]. A region of chromosome 11 in *D. rotundata* is associated with female heterogametic SNP markers and the presence of a female-specific (*W*-) genomic region suggest that sex determination in *D. rotundata* involves the *ZZ*/*ZW* system. Sex-linked regions of *D. alata* were identified by linkage analysis of F_1_ progeny using SNP markers [35]. Linkage between DNA markers and sex was detected on the distal part of linkage group 6 in the male consensus genetic maps, suggesting that *D. alata* employs an *XY*/*XX* sex-determination system. A genome-wide association study (GWAS) in *D. alata* identified ∼3 Mb sex-linked regions [43]. A recently assembled phased, telomere-to-telomere genome assembly of *D. alata* contains a ∼7.6 Mb potential sex-determination region (SDR) including a pericentric inversion [44]. The authors identified 88 sex-linked candidate genes and suggested that the jasmonic acid biosynthesis and signaling pathways may be involved in sex determination. However, the evolutionary relationship of the SDR across *Dioscorea* species is unclear. For instance, the *D. alata* SDR corresponds to chromosome 6 [35,44], while that of *D. rotundata* was identified on chromosome 11 [27], which does not share homologous genes and gene order similarity with *D. alata* chromosome 6. These findings strongly suggest that sex determination systems have transitioned from one type to another during the evolution of this genus [35,42].

In this study, we focused on the dioecious wild yam *D. tokoro* Makino (Fig 1A-D; Fig S1), a diploid species (2*n* = 2*x* = 20) that is widely distributed in temperate East Asia, including Japan, Korea, and China [23,45]. A previous DNA marker linkage study suggested that this species employs a *XY*/*XX* male heterogametic sex-determination system [34]. Here, we obtained chromosome-scale and haplotype-resolved genome assemblies from female and male *D. tokoro* plants. Association analysis of the F_1_ progeny and a whole-genome comparison between female and male individuals revealed an SDR residing in the middle of chromosome 3 and containing *X*- and *Y*-specific regions. Based on transcriptome analysis, we identified the homeobox gene *BLH9* and the molecular chaperone gene *HSP90* as candidate genes for sex determination in *D. tokoro*.

**Fig 1.**
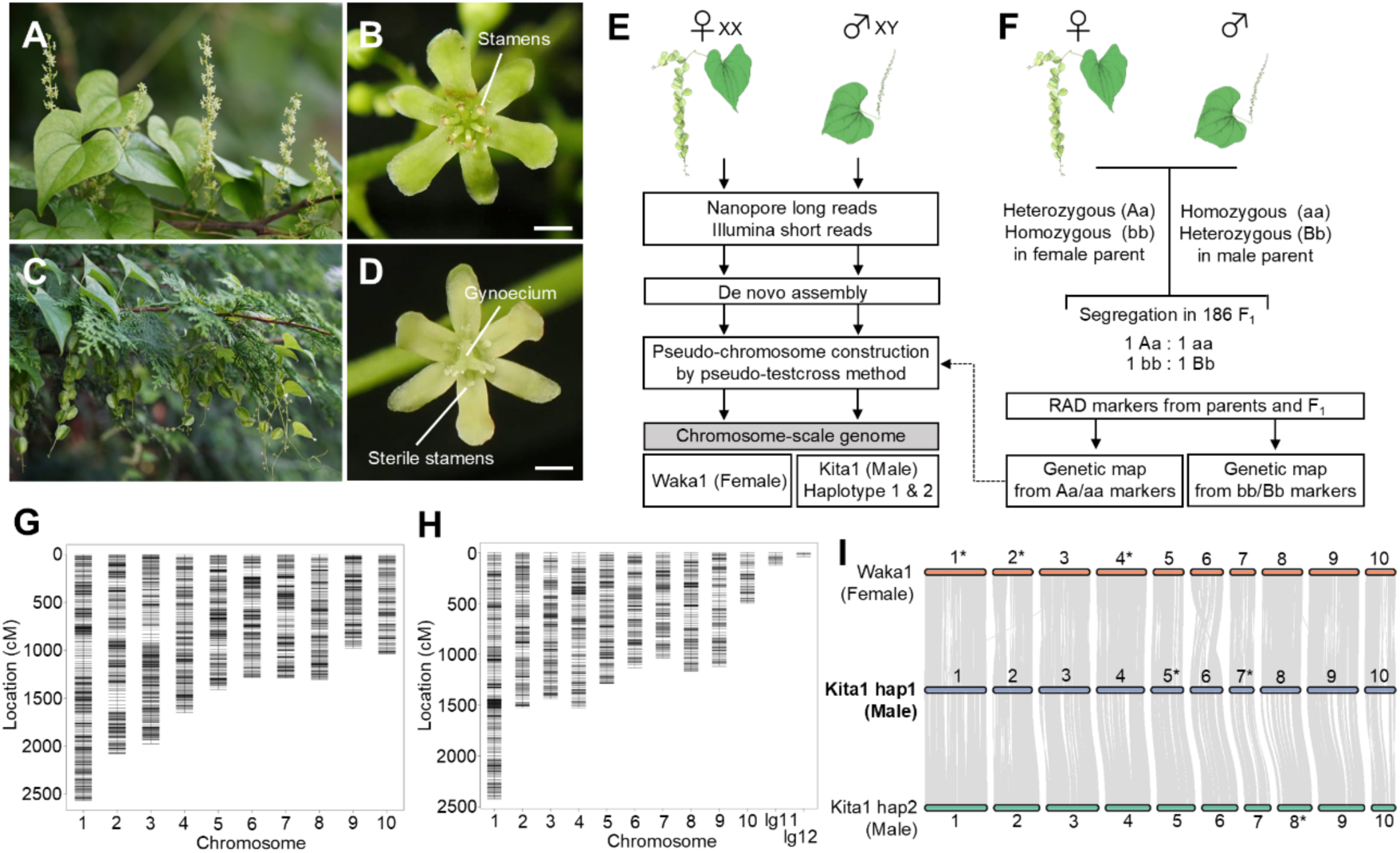
Haplotype-resolved genome assemblies constructed from male and female individuals of the dioecious wild yam *Dioscorea tokoro.* Male and female flowers are borne on separate individuals in *D. tokoro*. (**A-D**) The images show male inflorescences **(A)**, a male flower with stamens **(B)**, female inflorescences with capsular fruits **(C)**, and a female flower with a gynoecium and sterile stamens **(D)**. Scale bars, 1 mm. **(E)** A simplified scheme for generating chromosome-scale female and male reference genomes using long reads generated by Oxford Nanopore Technologies and Illumina sequencing. **(F)** A simplified scheme for anchoring the assemblies using RAD-seq-based linkage maps generated by the pseudo-testcross method using 186 F_1_ progenies. **(G, H)** Linkage maps based on female-parent-heterozygous markers **(G)** and based on male-parent-heterozygous markers **(H)**. **(I)** Chromosomal synteny of the female reference genome and two haplotypes of the male reference genome. The Kita1 (male) haplotype 1 reference genome has the most robust foundation, with ten chromosomes constructed from 18 contigs and two other reference genomes showing high collinearity with this genome. The chromosomes marked with asterisks were subjected to reverse complementation. The illustrations were obtained with TogoTV (copyright 2016 DBCLS TogoTV / CC-BY-4.0).

## Results

### Reconstructing the genome sequence of *Dioscorea tokoro* using genome assembly and linkage analysis

We reconstructed a chromosome-scale, haplotype-resolved genome assembly from female and male individuals of *D. tokoro* using whole-genome sequencing reads obtained using the Oxford Nanopore Technologies (ONT) and Illumina sequencing platforms (Fig 1E). The genome size of *D. tokoro* was estimated to be 388 Mb by flow cytometry analysis (Fig S2). We generated a consensus assembly from the filtered ONT reads from a female Waka1 individual (*XX* genotype). We also obtained separate assemblies for haplotype 1 and 2 of a male Kita1 individual (*XY* genotype) using filtered ONT reads. After polishing the draft assemblies using Illumina reads, we obtained a final assembly of 438.1 Mb containing 1,821 contigs for Waka1 (female). Likewise, for Kita1 (male), we obtained a haplotype 1 assembly with 172 contigs (414.4 Mb in total) and a haplotype 2 assembly with 440 contigs (300.4 Mb in total). Notably, the Kita1 haplotype 1 contained six telomere-to-telomere assemblies. BUSCO analysis [46] showed that the percentages of complete embryophyte BUSCOs were 97.6% in the Waka1 (female) assembly, 98.0% in the haplotype 1 Kita1 (male) assembly, and 76.0% in the haplotype 2 Kita1 (male) assembly (Table 1). The relatively low assembly size of the haplotype 2 assembly was probably caused by the use of the PECAT program, which prioritize the assembly of haplotype 1; similar biases were previously reported [47].

**Table 1.** Summary of the genome assemblies of Waka1 (female) and Kita1 (male) haplotypes 1 and 2.

|  |  | Waka1 (Female) | Kita1 (male) |  |
| --- | --- | --- | --- | --- |
| Chromosome number |  | 2n = 20 | 2n = 20 |  |
| Assembly type |  | Pseudo-haploid | Haplotype-resolved |  |
|  |  |  | Haplotype 1 | Haplotype 2 |
| Genome assembly | Total number of contigs | 1,821 | 172 | 440 |
|  | Total base pairs (bp) | 438,105,977 | 414,436,429 | 300,420,452 |
|  | Average contig size (bp) | 240,585 | 2,409,514.10 | 682,773.80 |
|  | Longest contig (bp) | 22,951,365 | 56,338,240 | 5,180,395 |
|  | Shortest contig (bp) | 600 | 1,051 | 868 |
|  | N50 (bp) | 981,934 | 33,850,955 | 1,379,077 |
| Genome evaluation<br>(BUSCO:<br>eudicots_odb10) | Total BUSCO groups searched | 1,614 | 1,614 | 1,614 |
|  | Complete | 97.60% | 98.00% | 76.00% |
|  | Complete and single copy | 92.90% | 96.00% | 73.50% |
|  | Complete and duplicated | 4.70% | 1.90% | 2.50% |
|  | Fragmented | 1.60% | 1.30% | 1.50% |
|  | Missing | 0.80% | 0.70% | 22.50% |

To anchor the assembled sequences to chromosomes by linkage analysis, we used a pseudo-testcross approach [48] (Fig 1F). We obtained 186 F_1_ progeny from a cross between Waka1 (female) and Kita1 (male) (Fig S3) and genotyped these progeny and the two parent plants by restriction-site-associated DNA sequencing (RAD-seq) [49]. We selected SNP and presence/absence polymorphism (PA) markers that were heterozygous in the female parent and homozygous in the male parent as female-parent-heterozygous markers. We also obtained SNP and PA markers that were homozygous in the female parent and heterozygous in the male parent as male-parent-heterozygous markers. Using the pseudo-testcross scheme, we constructed two separate linkage maps using the female-parent-heterozygous markers and the male-parent-heterozygous markers, respectively (Fig 1G, H; Fig S4, S5, S6). Finally, we combined the two linkage groups using the shared assemblies between the two linkage maps and generated pseudo-chromosomes 1–10 (Fig S7, S8, S9); for simplicity, we refer to the pseudo-chromosomes as chromosomes hereafter. The chromosome-assigned sequence using the Kita1 (Male) haplotype 1 reference appears to be the most complete, with ten chromosomes containing only 18 contigs.

To annotate the gene models, we obtained RNA-seq data from 18 different organs of *D. tokoro* and used protein homology information for *D. alata* TDa95/00328 and *D. rotundata* TDr96_F1. Based on the combination of transcriptome-based gene identification and *ab initio* gene prediction, we identified 37,712 genes for the Waka1 (female) assembly, 33,530 genes for the Kita1 (male) haplotype 1 assembly, and 26,242 genes for the Kita1 (male) haplotype 2 assembly (Table 2). Chromosomal synteny analysis based on collinear blocks with predicted genes revealed high collinearity between the Kita1 (male) haplotype 1-based chromosomes with the largest number of telomere-to-telomere assemblies and the chromosomes inferred using the two other reference genomes (Fig 1I). We judge that the chromosome-scale genome assemblies obtained from female and male individuals are useful to infer the structures of *X*- and *Y*-specific genomic regions.

**Table 2.** Summary of predicted genes in the genome assemblies of Waka1 (female) and Kita1 (male) haplotypes 1 and 2.

|  |  | Waka1 (female) | Kita1 (male) |  |
| --- | --- | --- | --- | --- |
| Chromosome number |  | 2n = 20 | 2n = 20 |  |
| Assembly type |  | Pseudo-haploid | Haplotype-resolved |  |
|  |  |  | haplotype 1 | haplotype 2 |
| Gene prediction | Number of predicted genes |  |  |  |
|  | StringTie | 34,943 | 33,530 | 26,242 |
|  | BRAKER2 | 17,264 | 16,965 | 12,983 |
|  | Total number of genes | 37,712 | 36,483 | 28,345 |
|  | (Total transcript variants) | (43,961) | (42,915) | (33,306) |
| Gene evaluation<br>(BUSCO: eudicots_odb10) | Total BUSCO groups searched | 1,614 | 1,614 | 1,614 |
|  | Complete | 92.70% | 92.80% | 72.20% |
|  | Complete and single-copy | 62.50% | 64.10% | 48.00% |
|  | Complete and duplicated | 30.20% | 28.70% | 24.20% |
|  | Fragmented | 4.60% | 4.20% | 3.60% |
|  | Missing | 2.70% | 3.00% | 24.20% |

### Linkage study and mapping coverage analysis of Illumina short reads identify the *X*- and *Y*-specific genomic regions on chromosome 3

To identify the genomic region linked to the sex phenotype, we performed association analysis using the 186 F_1_ progeny, comprising 38 females, 89 males, and 59 non-flowering individuals. Based on RAD-seq data from the 127 flowering F_1_ progeny, we examined the association between the genotypes and sex phenotype of the F_1_ individuals using Fisher’s exact test (Fig 2A). The log-transformed *q*-values (–log_10_(*q*)) revealed significant associations between sex phenotype and the male-parent-heterozygous markers in the middle of chromosome 3, but did not detect any association using the female-parent-heterozygous markers (Fig 2B, C; Fig S10). Linkage analysis using composite interval mapping performed with R/qtl [50] also detected linkage between sex phenotype and the male-parent-heterozygous markers on chromosome 3, whereas no linkage was detected with the female-parent-heterozygous markers (Fig S11). These results suggest that the sex of *D. tokoro* is determined by the component on chromosome 3 and that the species has a male heterogametic sex-determination system of *XY*/*XX*.

**Fig 2.**
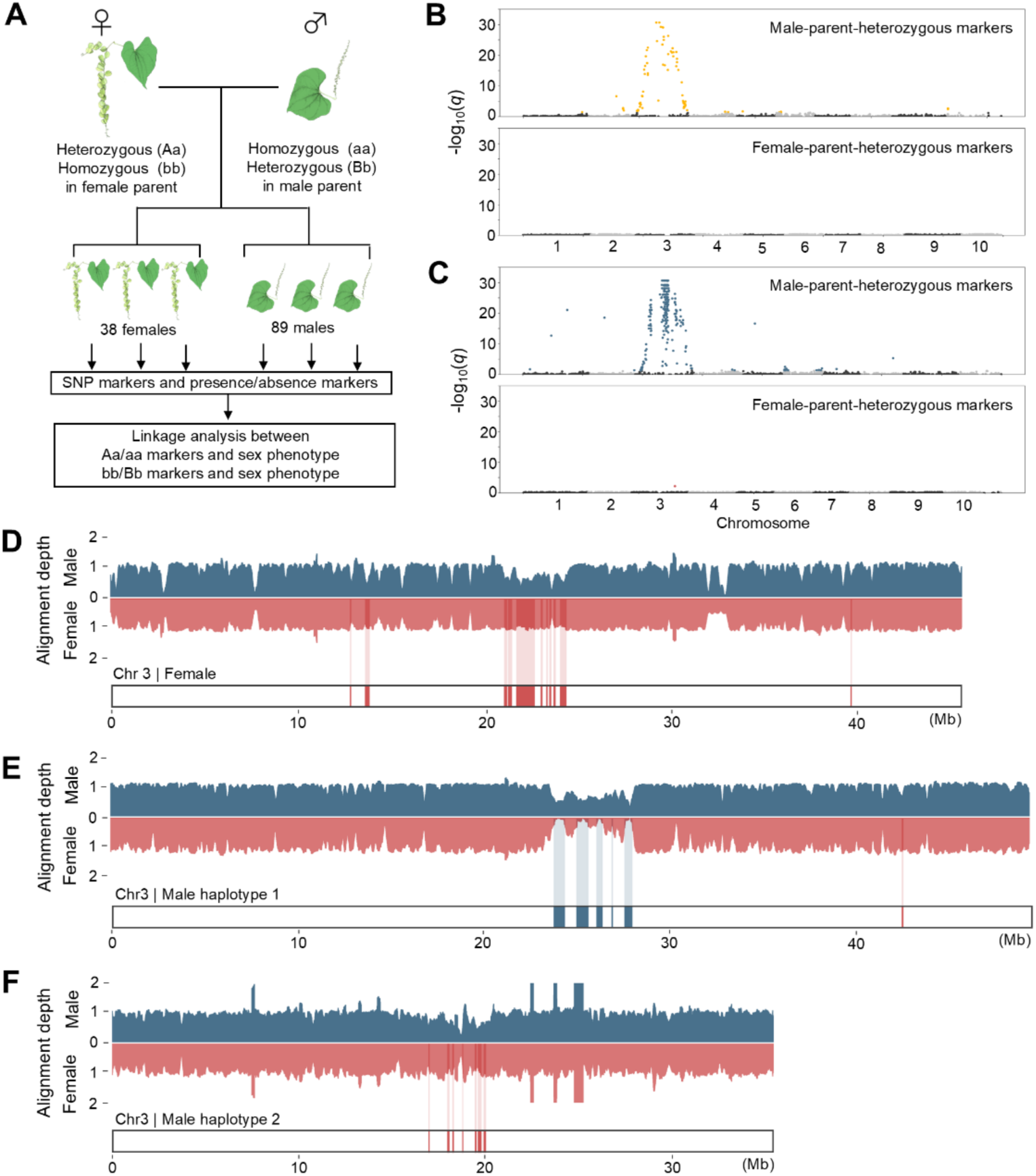
The dioecious wild yam *Dioscorea tokoro* contains *X*- and *Y*-specific regions on chromosome 3. **(A)** Schematic diagram of association analysis to identify genomic regions associated with sex phenotype. SNP and presence/absence (PA) markers were detected in 38 females and 89 males in the F_1_ progeny. **(B)** Manhattan plots of –log_10_(*q*) values between SNP-type markers and sex phenotype on the haplotype 1 assembly from Kita1 (male). The –log_10_(*q*) values were obtained by Fisher’s exact test. The upper and lower plots show male-parent-heterozygous markers and female-parent-heterozygous markers, respectively. The yellow points in the upper plot indicate SNP-type markers significantly associated with sex phenotype. A significance threshold of 5% false discovery rate was adjusted by Benjamini-Hochberg correction: adjusted threshold = 1.31 for plots of male-parent-heterozygous markers and adjusted threshold = 2.18 for plots of male-parent-heterozygous markers. **(C)** Manhattan plots of –log_10_(*q*) values between PA markers and sex phenotype. The red and blue plots show PA markers significantly associated with sex phenotype. The male-parent-heterozygous markers in the middle of chromosome 3 show significant associations. **(D)** Mapping coverage analysis to differentiate between putative *X*- and *Y*-specific regions. The bar graphs indicate female and male mapping depth, obtained by mapping Waka1 (female) and Kita1 (male) Illumina sequences onto the assembly from Waka1 (female). Coverage analysis revealed *X*-specific regions of chromosome 3, shown as red regions below the bar graphs. Chromosome 3 was defined as the *X* chromosome. **(E)** Bar graphs showing female and male mapping depth, obtained by aligning Waka1 (female) and Kita1 (male) Illumina sequences onto the haplotype 1 assembly from Kita1 (male). Coverage analysis revealed *Y*-specific regions on chromosome 3, shown as blue regions below the bar graphs. Chromosome 3 was defined as the *Y* chromosome. The non-zero coverage in female mapping within the *Y*-specific regions might have been caused by misalignment of repeat-rich regions (suggested in Fig 3B). **(F)** Bar graphs showing female and male alignment depth, obtained by aligning Waka1 (female) and Kita1 (male) Illumina sequences onto the haplotype 2 assembly from Kita1 (male). Coverage analysis revealed *X*-specific regions on chromosome 3, shown as red regions below the bar graphs.

In the genomic region associated with sex phenotype, we detected a higher concentration of PA markers (188 markers in the 10 Mbp region around the marker showing the peak association value) than SNP markers (20 markers in the 10 Mbp region) (Fig 2B, C), indicating structural differences in the SDR. We examined the major structural differences in this region by performing coverage analysis of Illumina short-read sequences of the male and female genomes mapped to the reference genome sequence. In diploid plants with male heterogametic sex determination (*XY*) and substantial structural differentiation between *X* and *Y* chromosomes, the *Y*-specific region is predicted to be absent from females. Furthermore, the dosage of the male *X*-specific region is thought to be one-half that of females (hemizygous) [51]. To assess whether this is the case for *D. tokoro*, we aligned the Illumina short reads obtained from female and male individuals to the reference genome of chromosome 3 and studied the mapping depth. As the reference genome sequences for mapping, we used three different chromosome 3 assemblies: the Waka1 (female) assembly and the Kita1 (male) haplotype 1 and 2 assemblies. We identified regions covered by male reads at 0.5× depth and female reads at 1× depth in the center of chromosome 3 in the Waka1 (female) reference genome (Fig 2D). When we mapped the Illumina short reads to the Kita1 (male) haplotype 1 assembly, we identified regions covered by male reads at 0.5× depth and female reads at 0× depth (Fig 2E). We also identified a region covered by male reads at 0.5× depth and female reads at 1× depth in the haplotype 2 assembly (Fig 2F). We also observed such differences in depth between male and female reads when male and female plants collected from three different locations were used for mapping (Fig S12, S13).

Based on the results of our association analysis (Fig 2B, C) and mapping coverage analysis, we defined chromosome 3 of the assembly from Waka1 (female) as the *X* chromosome and chromosome 3 of the haplotype 1 assembly from Kita1 (male) as the *Y* chromosome.

### *X*- and *Y*-specific regions are likely located in the peri-centromeric regions

Recombination suppression is a key step in SDR evolution. This suppression is caused by various mechanisms, such as the accumulation of repetitive DNA and chromosome inversion [6,52]. The *X*- and *Y*-specific regions were located in a genomic region with lower gene density and higher density of repetitive DNA sequences, represented by gypsy LTRs (Fig 3A, B). Centromeres and pericentromeric (near the centromere) regions are known to have suppressed recombination [53] and contain a higher density of retrotransposons and transposons [54,55]; particularly high levels of gypsy LTRs are reported in multiple monocot species [56–58], suggesting that *D. tokoro X*- and *Y*-specific regions likely correspond to the pericentromeric regions (near the centromere) of chromosome 3.

**Fig 3.**
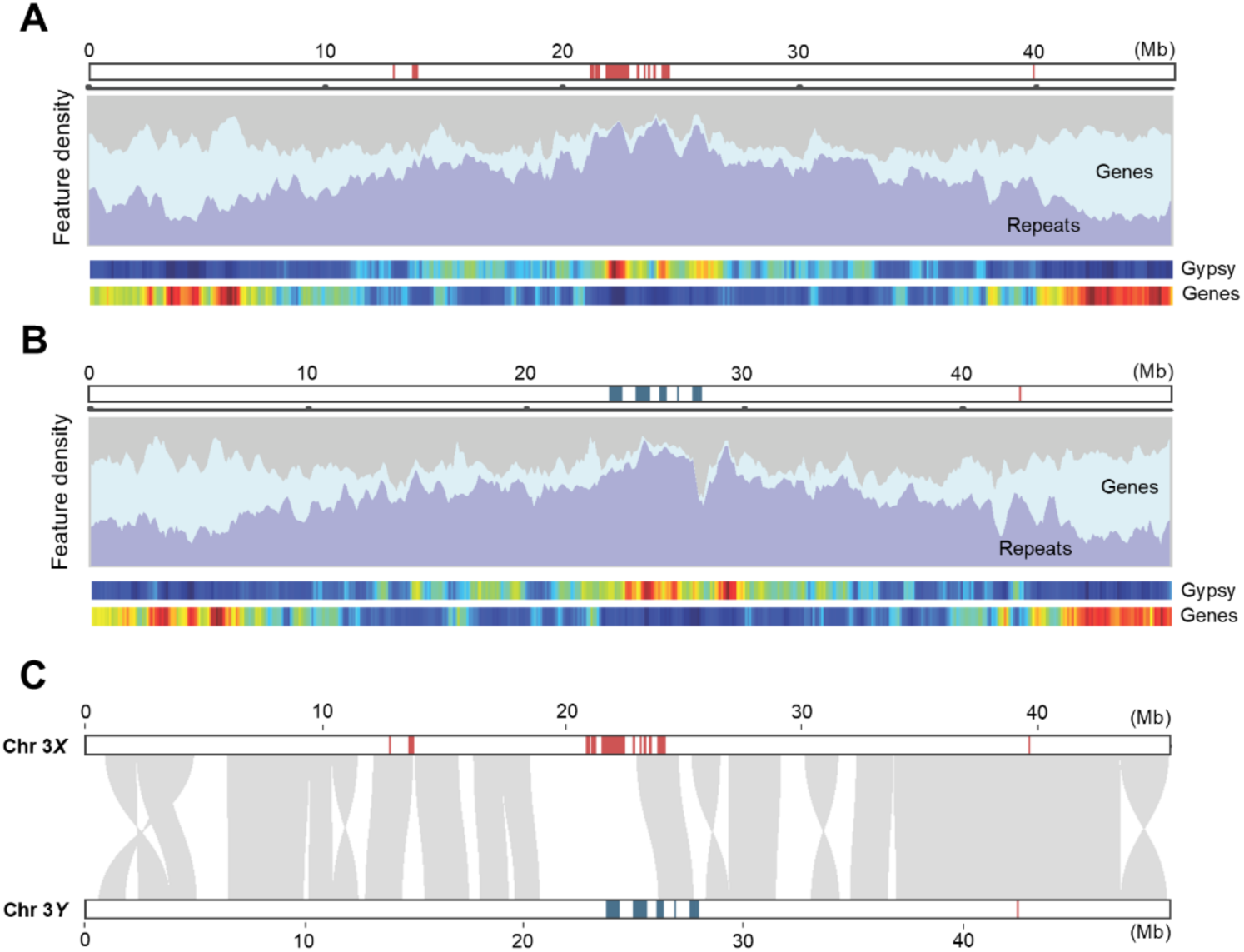
*X*- and *Y*-specific regions locate in peri-centromeric regions with no major inversions. **(A)** Repetitive DNA sequences accumulate in *X*-specific regions on chromosome 3, suggesting that these regions are located in the peri-centromeric regions. The bar graphs indicate gene and retrotransposon density on chromosome 3. The heatmap indicates gene and Gypsy retrotransposon density. **(B)** Repetitive DNA sequences accumulate in *Y*-specific regions of chromosome 3, suggesting that these regions are located in the peri-centromeric regions. **(C)** Chromosomal synteny analysis of the *X* and *Y* chromosomes showing no major inversions in the chromosomes and the differentiation of the *X*- and *Y*-specific regions.

To investigate possible inversions and the accumulation of repetitive DNA sequences around the *X*- and *Y*-specific regions, we performed synteny analysis and visualization of gene and repetitive sequence density. Analysis of chromosomal synteny between the *X* and *Y* chromosomes revealed no major inversions. However, the lack of a synteny block in the middle position, around the *X*- and *Y*-specific regions, indicates differentiation of the *X* and *Y* chromosomes (Fig 3C)

We also evaluated the recombination rate around the *X*- and *Y*-specific regions by focusing on SNP markers segregating among the F_1_ progeny and comparing the relationship between the linkage distances (in centimorgans, cM) and physical distances (in base pairs, bp) (Fig S14, S15). Although we observed a little reduction of linkage distances in the center of the chromosome 3, its pattern of relationship between linkage distances and physical distances was not greatly different from those of other chromosomes. This result suggests that there is no chromosome-wide large-scale suppression of recombination in *D. tokoro* sex chromosomes. These results suggest *D. tokoro* SDR is located in the vicinity of centromere and recombination suppression around the centromere has contributed to the evolution of *X*- and *Y*-regions.

### The *Y-*specific genes *BLH9* and *HSP90* are specifically expressed during early male flower development

The male heterogametic sex determination (*XY*) in *D. tokoro* suggests that *Y*-specific genes or miRNAs function in sex determination. To identify the candidate genes or miRNAs involved in sex determination, we conducted transcriptome analysis via RNA-seq and small RNA-seq. In *D. tokoro*, male and female flowers show sex-specific phenotypes after the bud stage (Fig 4A, B), indicating that the determination of sex phenotypes takes place before this stage. Therefore, we obtained RNA-seq and small RNA-seq data from male and female flowers at five stages of development as well as non-reproductive organs (Fig 4C, D). We mapped the RNA-seq reads to the haplotype 1 Kita1 (male) assembly, which includes the *Y* chromosome.

**Fig 4.**
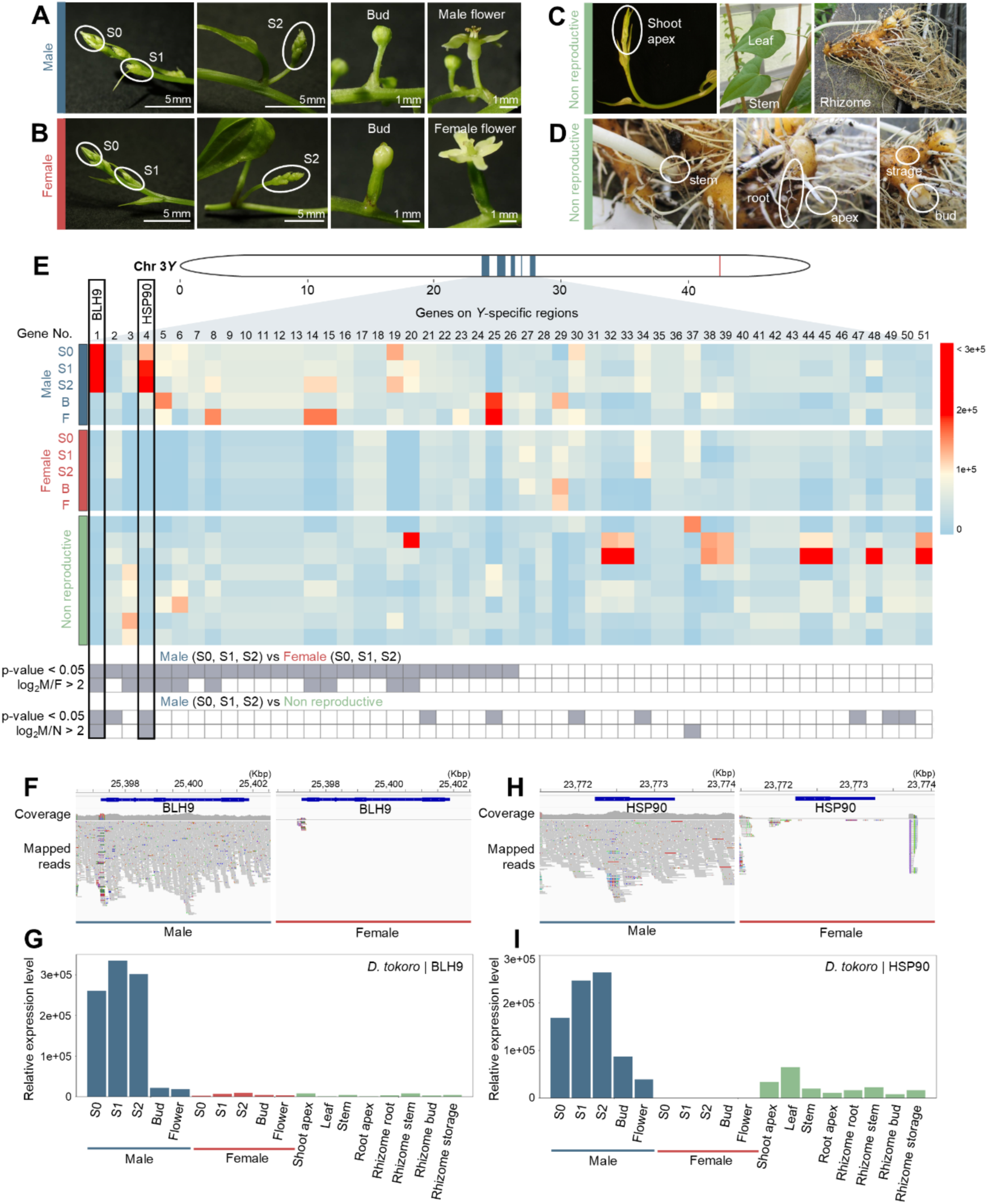
The candidate genes *BLH9* and *HSP90* are located on *Y*-specific regions and are highly expressed during early male flower development. (A) Five stages of male flower development used for RNA-seq and small RNA-seq analyses. **(B)** Five stages of female flower development. **(C)** Non-reproductive parts of the plant, including the shoot apex, leaf, stem, and rhizome. The shoot apex is indicated with ellipses. **(D)** Parts of the rhizome, including the rhizome stem, apex, root, bud, and storage region. Each rhizome region is indicated with ellipses. **(E)** Heatmap of the expression levels of genes in the *Y*-specific region. The boxes under the heatmap show the DESeq2 results for each gene. The gray boxes indicate *p*-value < 0.05 based on differential expression testing by negative binomial generalized linear model or log_2_FC > 2 based on the log_2_ ratio of the mean of normalized counts in each group in two comparisons: male (S0, S1, S2) vs. female (S0, S1, S2) and male (S0, S1, S2) vs. non-reproductive organs. *BLH9* and *HSP90*, highlighted by rectangles in the heatmap, are highly expressed during early stages of male flower development. **(F)** Coverage of short reads from male and female individuals on *BLH9*. *BLH9* was covered only by male short reads. **(G)** Relative expression levels of *BLH9* in male and female flowers at five stages of development and in non-reproductive organs. **(H)** Coverage of short reads on *HSP90* from male and female individuals. *HSP90* was covered only by male short reads. **(I)** Relative expression levels of *HSP90*. All expression levels are shown as normalized TPM values.

The *Y*-specific regions contained 51 potential protein-coding genes (Fig 4E; Table 3) and 15 miRNAs (Table 4). Of these, 26 genes (genes no. 1-26 in Fig 4E) were more highly expressed during the three stages of early male vs. female flower development based on the hypothesis test for differential expression of the RNA-seq data using negative binomial generalized linear models (*p*-value < 0.05), whereas no miRNAs showed differential expression between male and female flowers. A sequence similarity search with BLASTX revealed that the 26 genes included eight genes encoding hypothetical proteins without functional information and 18 genes encoding proteins with functional information: *BEL1-LIKE HOMEODOMAIN PROTEIN 9, L10-INTERACTING MYB DOMAIN-CONTAINING PROTEIN*, *HEAT SHOCK PROTEIN 81-1*, seven *SMALL RIBOSOMAL SUBUNIT PROTEIN MS47* genes, two *RIBONUCLEOSIDE-DIPHOSPHATE REDUCTASE SMALL CHAIN A* genes, *SPLICING FACTOR CACTIN*, *Dr1-associated corepressor*, *SERINE/THREONINE-PROTEIN KINASE SAPK1*, two *PROTEIN PHOSPHATASE 2C 70*, and *EXOCYST COMPLEX COMPONENT EXO70C1* (Table 3).

**Table 3.**
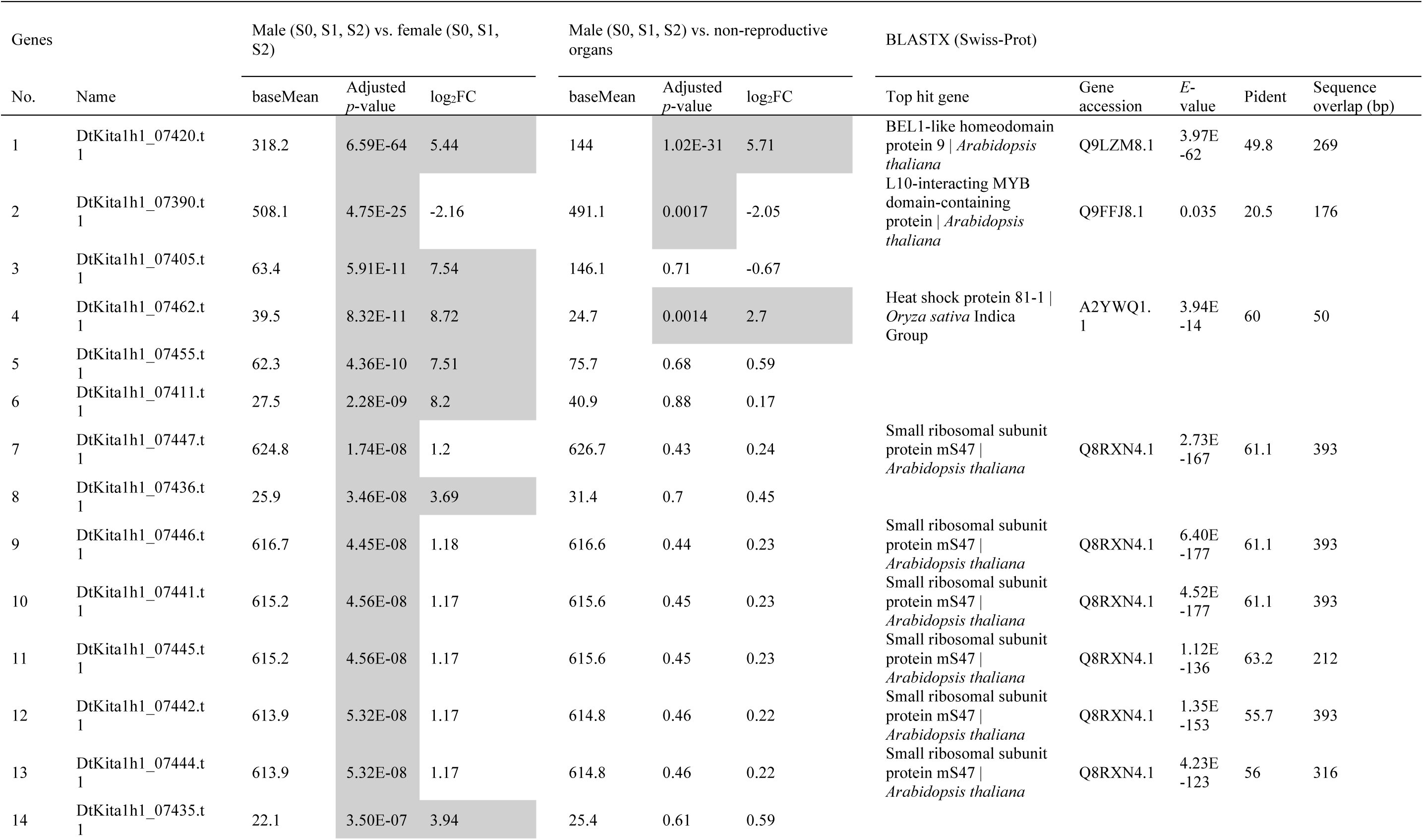
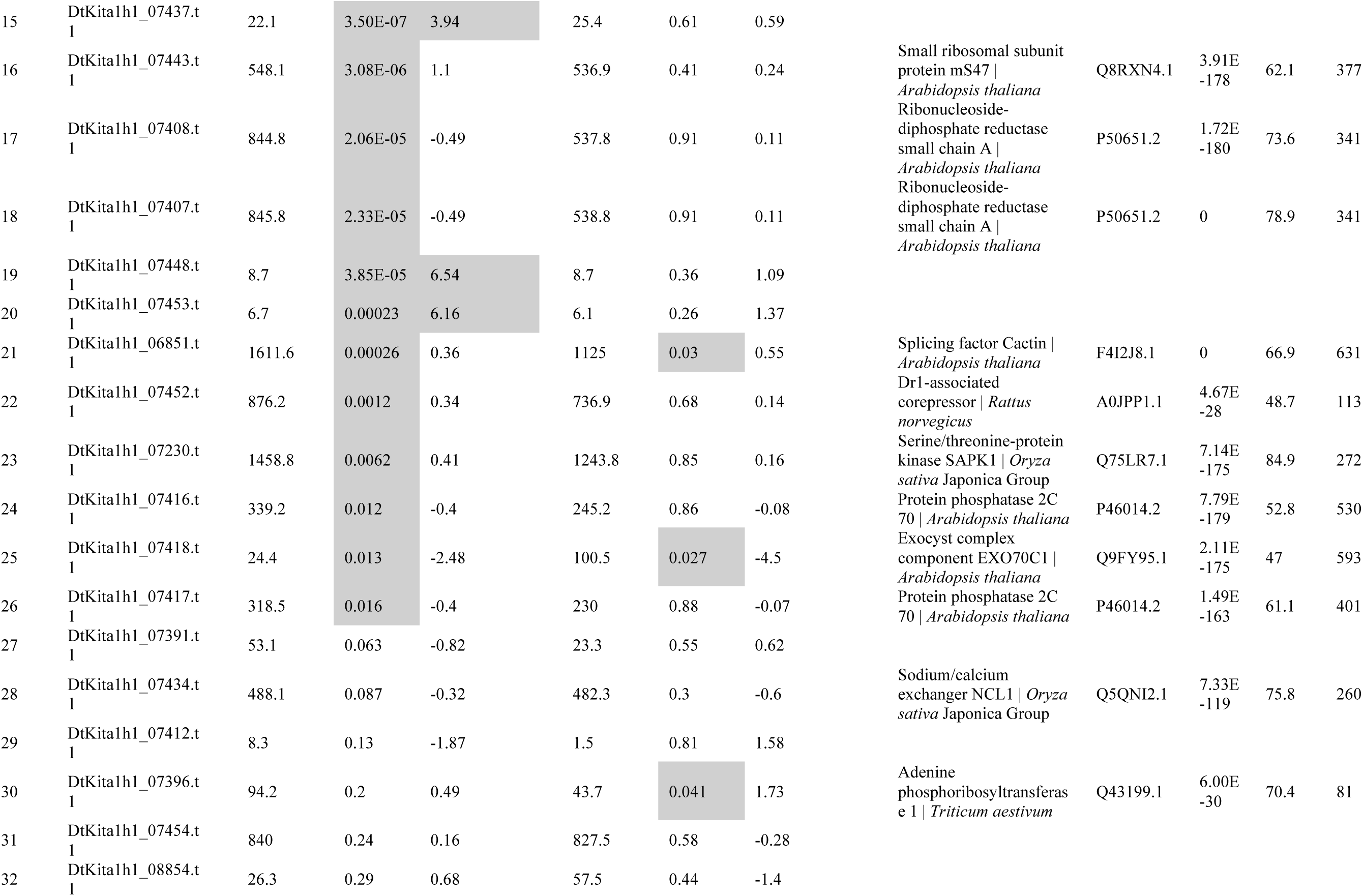
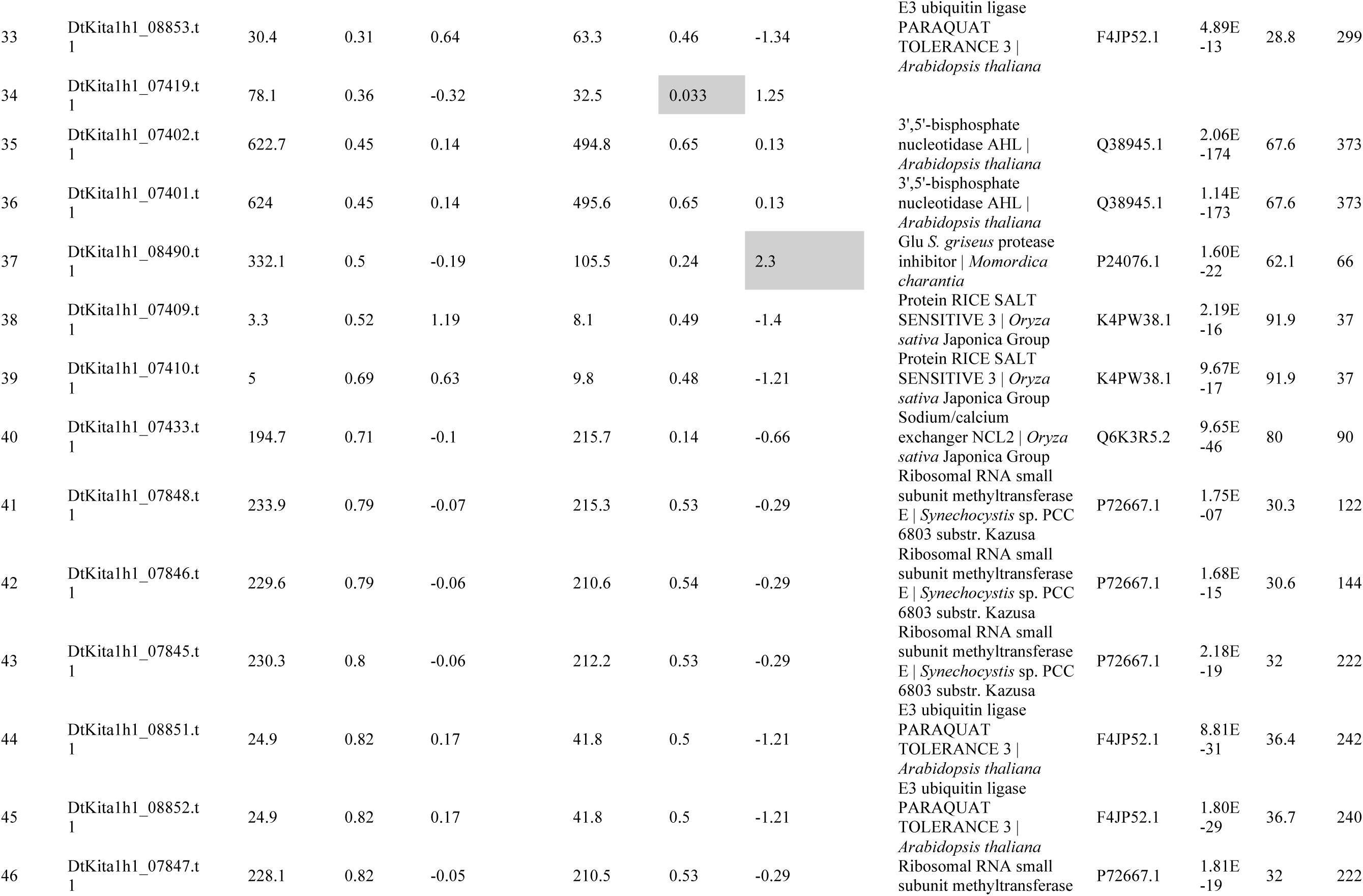
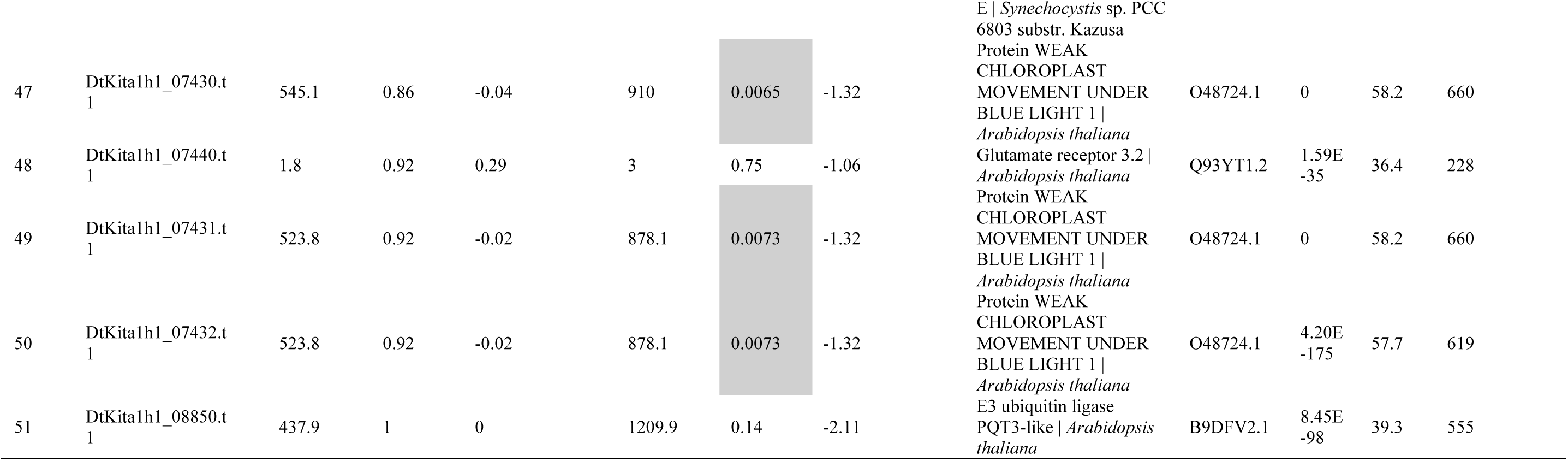
Differential expression analysis of genes located in *Y*-specific regions.

**Table 4.** Differential expression analysis of miRNAs located on *Y*-specific regions.

| miRNA no. | Name | Length | Male (S0, S1, S2) vs. female (S0, S1, S2) |  |  | Candidate position |  |  |  |  |  |  |
| --- | --- | --- | --- | --- | --- | --- | --- | --- | --- | --- | --- | --- |
|  |  |  | baseMean | Adjusted *p*-value | log<sub>2</sub>FC | Start position of *Y*-specific region | End position of *Y*-specific region | Start position of miRNA | End position of miRNA | Matched length | Mismatch | Gap open |
| 1 | Dtokoro_miRDP2_mature_88 | 20 | 12.75139983 | 0.181348204 | -1.616847994 | 24040219 | 24040234 | 5 | 20 | 16 | 0 | 0 |
| 2 | Dtokoro_miRDP2_mature_33 | 24 | 77.13486872 | 0.274648861 | 0.905878248 | 8961492 | 8961507 | 2 | 17 | 16 | 0 | 0 |
|  |  |  |  |  |  | 14843719 | 14843734 | 1 | 16 | 16 | 0 | 0 |
| 3 | Dtokoro_miRDP2_mature_319 | 21 | 45.39343813 | 0.386928618 | 0.873273701 | 25082756 | 25082772 | 5 | 21 | 17 | 0 | 0 |
| 4 | Dtokoro_miRDP2_mature_494 | 21 | 5.394040488 | 0.744233479 | -1.154677187 | 24387813 | 24387833 | 1 | 21 | 21 | 0 | 0 |
|  |  |  |  |  |  | 24387857 | 24387877 | 1 | 21 | 21 | 0 | 0 |
| 5 | Dtokoro_miRDP2_mature_341 | 21 | 1.094048623 | 0.989796571 | 0.047071366 | 24389458 | 24389477 | 2 | 21 | 20 | 0 | 0 |
| 6 | Dtokoro_miRDP2_mature_105 | 23 | 0 | NA | NA | 24369832 | 24369850 | 4 | 22 | 19 | 1 | 0 |
| 7 | Dtokoro_miRDP2_mature_149 | 21 | 0 | NA | NA | 24257718 | 24257740 | 1 | 21 | 23 | 0 | 2 |
|  |  |  |  |  |  | 25680716 | 25680731 | 1 | 16 | 16 | 0 | 0 |
|  |  |  |  |  |  | 25683373 | 25683388 | 1 | 16 | 16 | 0 | 0 |
|  |  |  |  |  |  | 25686041 | 25686056 | 1 | 16 | 16 | 0 | 0 |
| 8 | Dtokoro_miRDP2_mature_185 | 24 | 0 | NA | NA | 26935365 | 26935383 | 3 | 21 | 19 | 0 | 0 |
|  |  |  |  |  |  | 26937818 | 26937836 | 3 | 21 | 19 | 0 | 0 |
|  |  |  |  |  |  | 26366483 | 26366501 | 3 | 21 | 19 | 1 | 0 |
| 9 | Dtokoro_miRDP2_mature_242 | 22 | 0.111576562 | NA | -1.860020433 | 25384647 | 25384662 | 3 | 18 | 16 | 0 | 0 |
| 10 | Dtokoro_miRDP2_mature_303 | 23 | 0 | NA | NA | 24457752 | 24457770 | 3 | 21 | 19 | 1 | 0 |
| 11 | Dtokoro_miRDP2_mature_353 | 21 | 0 | NA | NA | 25294304 | 25294323 | 2 | 21 | 20 | 0 | 0 |
|  |  |  |  |  |  | 27131539 | 27131558 | 2 | 21 | 20 | 0 | 0 |
|  |  |  |  |  |  | 25274241 | 25274259 | 1 | 19 | 19 | 0 | 0 |
|  |  |  |  |  |  | 25227145 | 25227165 | 1 | 21 | 21 | 1 | 0 |
|  |  |  |  |  |  | 25235180 | 25235200 | 1 | 21 | 21 | 1 | 0 |
|  |  |  |  |  |  | 25273788 | 25273808 | 1 | 21 | 21 | 1 | 0 |
|  |  |  |  |  |  | 25273900 | 25273920 | 1 | 21 | 21 | 1 | 0 |
|  |  |  |  |  |  | 25274012 | 25274032 | 1 | 21 | 21 | 1 | 0 |
|  |  |  |  |  |  | 26343213 | 26343233 | 1 | 21 | 21 | 1 | 0 |
|  |  |  |  |  |  | 27514790 | 27514810 | 1 | 21 | 21 | 1 | 0 |
|  |  |  |  |  |  | 27515573 | 27515593 | 1 | 21 | 21 | 1 | 0 |
|  |  |  |  |  |  | 25266511 | 25266530 | 2 | 21 | 20 | 1 | 0 |
|  |  |  |  |  |  | 25232780 | 25232798 | 3 | 21 | 19 | 1 | 0 |
|  |  |  |  |  |  | 25274124 | 25274142 | 3 | 21 | 19 | 1 | 0 |
|  |  |  |  |  |  | 25293969 | 25293987 | 3 | 21 | 19 | 1 | 0 |
|  |  |  |  |  |  | 27514460 | 27514478 | 3 | 21 | 19 | 1 | 0 |
|  |  |  |  |  |  | 27514680 | 27514698 | 3 | 21 | 19 | 1 | 0 |
| 12 | Dtokoro_miRDP2_mature_445 | 24 | 0 | NA | NA | 24457752 | 24457770 | 4 | 22 | 19 | 1 | 0 |
| 13 | Dtokoro_miRDP2_mature_473 | 23 | 0 | NA | NA | 25384647 | 25384662 | 3 | 18 | 16 | 0 | 0 |
| 14 | Dtokoro_miRDP2_mature_485 | 21 | 0 | NA | NA | 25550894 | 25550913 | 2 | 21 | 20 | 1 | 0 |
| 15 | Dtokoro_miRDP2_mature_533 | 21 | 0 | NA | NA | 25384647 | 25384662 | 2 | 17 | 16 | 0 | 0 |

We also compared the expression levels of genes between male and female flowers based on the log_2_ ratio of the mean of normalized counts (log_2_[fold change], log_2_FC). We identified ten genes (genes no. 1, 3–6, 8, 14, 15, 19, 20 in Fig 4E) that showed more than 4-fold higher expression (log_2_FC > 2) during the early stages of male vs. female flower development; these genes were also included in the 26 genes identified by differential expression analysis described above. The ten genes included two genes encoding proteins with functional annotations: *A. thaliana BEL1-LIKE HOMEODOMAIN PROTEIN 9* (*AtBLH9*) and rice (*Oryza sativa*) *HEAT SHOCK PROTEIN 81-1*. The eight remaining genes encode hypothetical proteins without functional information.

When we compared the expression levels of the ten genes on the *Y*-specific region between male flowers and non-reproductive organs, only two genes (genes no. 1 and 4 in Fig 4E) had higher expression in male flowers than in non-reproductive organs (*p*-value < 0.05), and both these genes showed significantly higher expression in male flowers based on normalized counts (log_2_FC > 2). Thus, we identified two *Y*-chromosome-specific genes showing significantly higher expression in male flowers during the three stages of early development (*q*-value < 0.05 and log_2_FC > 2) compared to female flowers and non-reproductive organs. The first gene, named *BLH9*, is a male-specific gene covered only by male-specific short-read sequences (Fig 4F) that is upregulated in male flowers during the three stages of early development (Fig 4G). A sequence similarity search by BLASTX showed that *BLH9* is closely related to the homeobox gene *BEL1-LIKE HOMEODOMAIN PROTEIN 9* (*BLH9*) in the *BELL* family of *A. thaliana* (Table S5). The second gene, *HSP90*, is also male-specific (Fig 4H) and was upregulated in male flowers during the three stages of early development (Fig 4I). *HSP90* shares similarity with the *A. thaliana* chaperone gene *HEAT SHOCK PROTEIN 90-4* (*AtHSP90.4*) (Table 5). The male-specificity of both genes was confirmed by PCR amplification of the DNA fragment using gene-specific primers and DNA templates from individuals of two natural *D. tokoro* populations: the HNMK population (northern Japan) and the KMMT population (southern Japan) (Fig S16). In summary, The *Y*-specific genes *BLH9* and *HSP90,* specifically expressed during early male flower development, were identified as the primary candidate sex-determination genes.

**Table 5.**
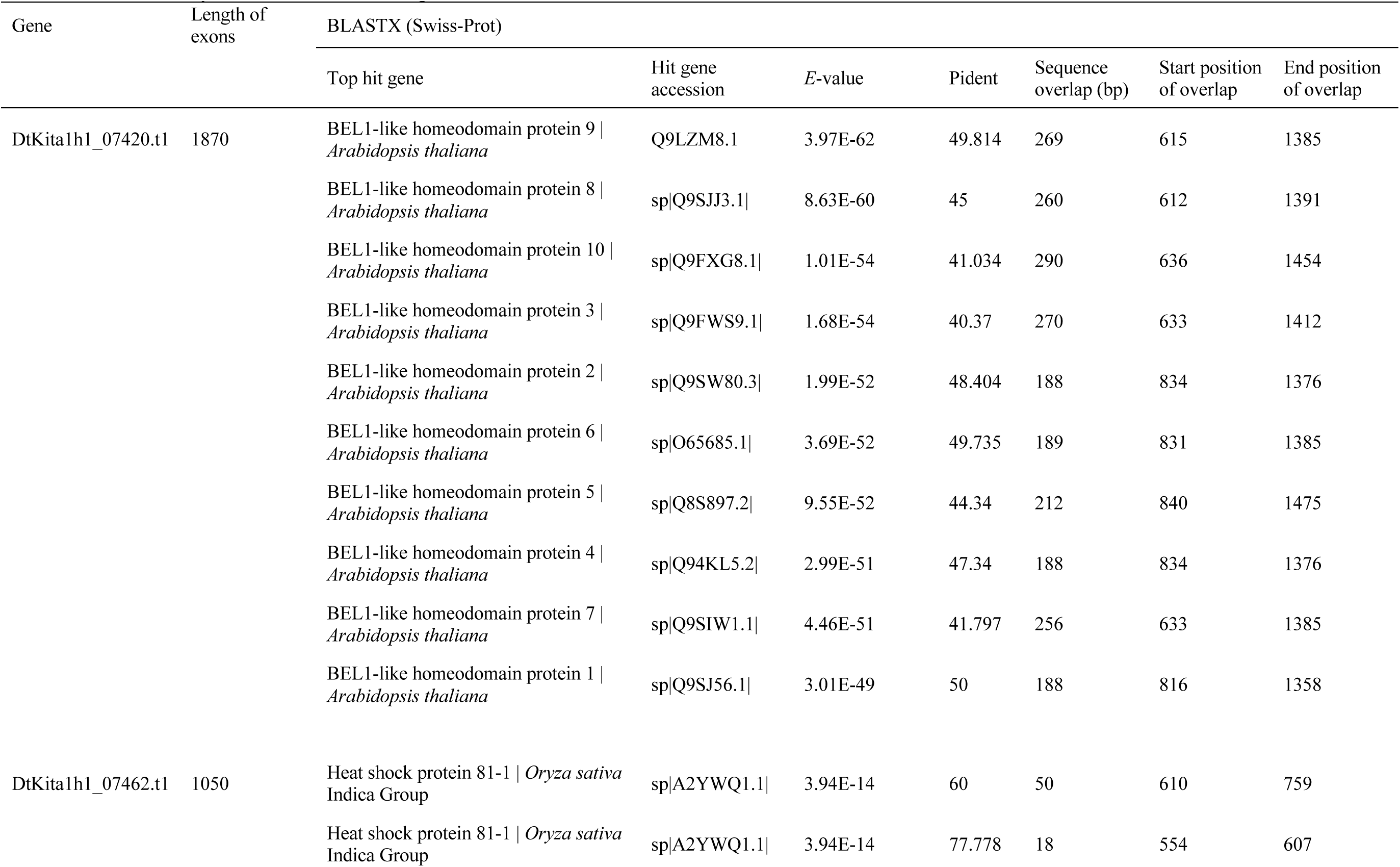
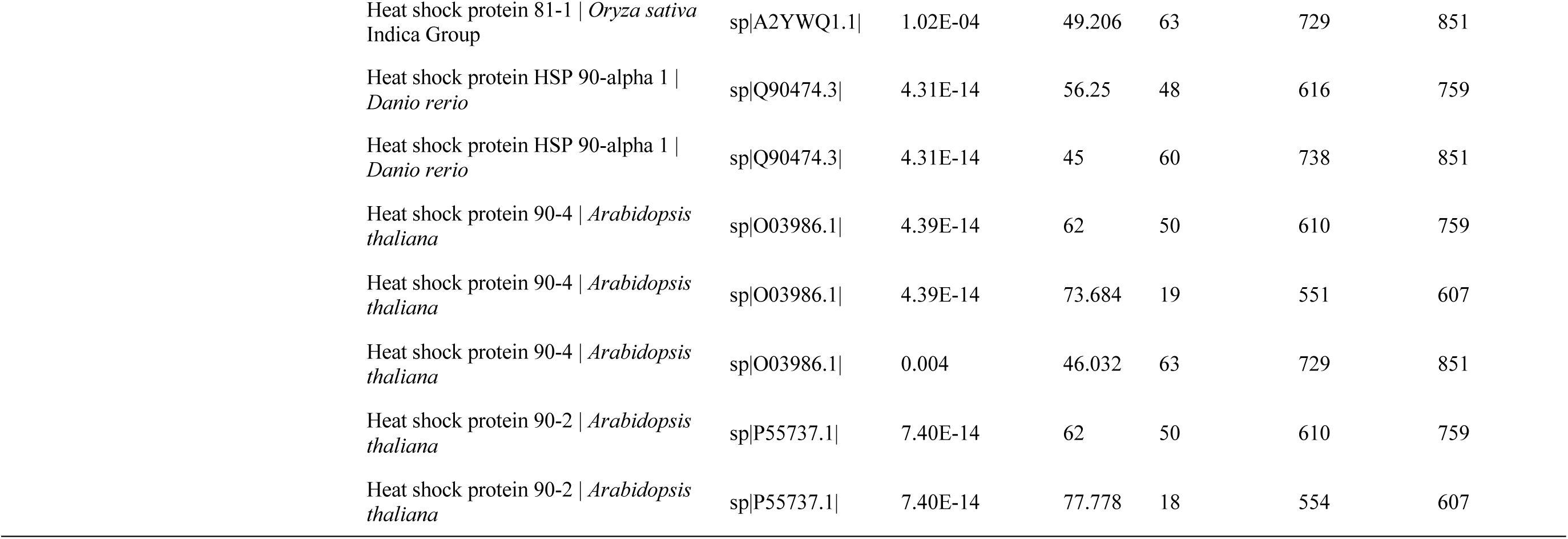
BLASTX analysis of the two candidate genes.

### *BLH9* shares similar functions with *AtBLH9* and suppresses fruit development in *Arabidopsis thaliana*

A genetic transformation method for *D. tokoro* has not yet been established. Among plants in the genus *Dioscorea*, successful transformation has been reported only in *D. rotundata* [59], which belongs to a section (Sect. Enantiophyllum) distant from *D. tokoro* (Sect. Stenophora) and possesses a *ZW/ZZ* sex determination system [27]. Therefore, it is difficult to validate the functions of *D. tokoro Y*-specific genes by transformation of *Dioscorea* species.

To explore the biological role of the candidate gene *BLH9*, we overexpressed this gene and *AtBLH9* in *A. thaliana*. Phylogenetic analysis of BLH9 and *A. thaliana* TALE (Three-amino-loop-extension) superfamily proteins showed that AtBLH9 shared the greatest amino acid sequence similarity with BLH9 (Fig 5A). AtBLH9 is involved in inflorescence architecture and fruit development, as overexpression or knock-out of *AtBLH9* led to abnormal inflorescence development, including reduced inflorescence height and shorter fruit length, as well as irregular internode elongation, with extremely short and long internodes, respectively [60–65]. To compare the biological roles of *AtBLH9* and *BLH9*, we overexpressed *AtBLH9* and *BLH9* in *A. thaliana* plants driven by the CaMV35S promoter. For phenotypic evaluation, we used T_2_ plants transformed with a construct expressing *BLH9* or *AtBLH9*. Wild-type *A. thaliana* Col-0 plants were used as a control.

**Fig 5.**
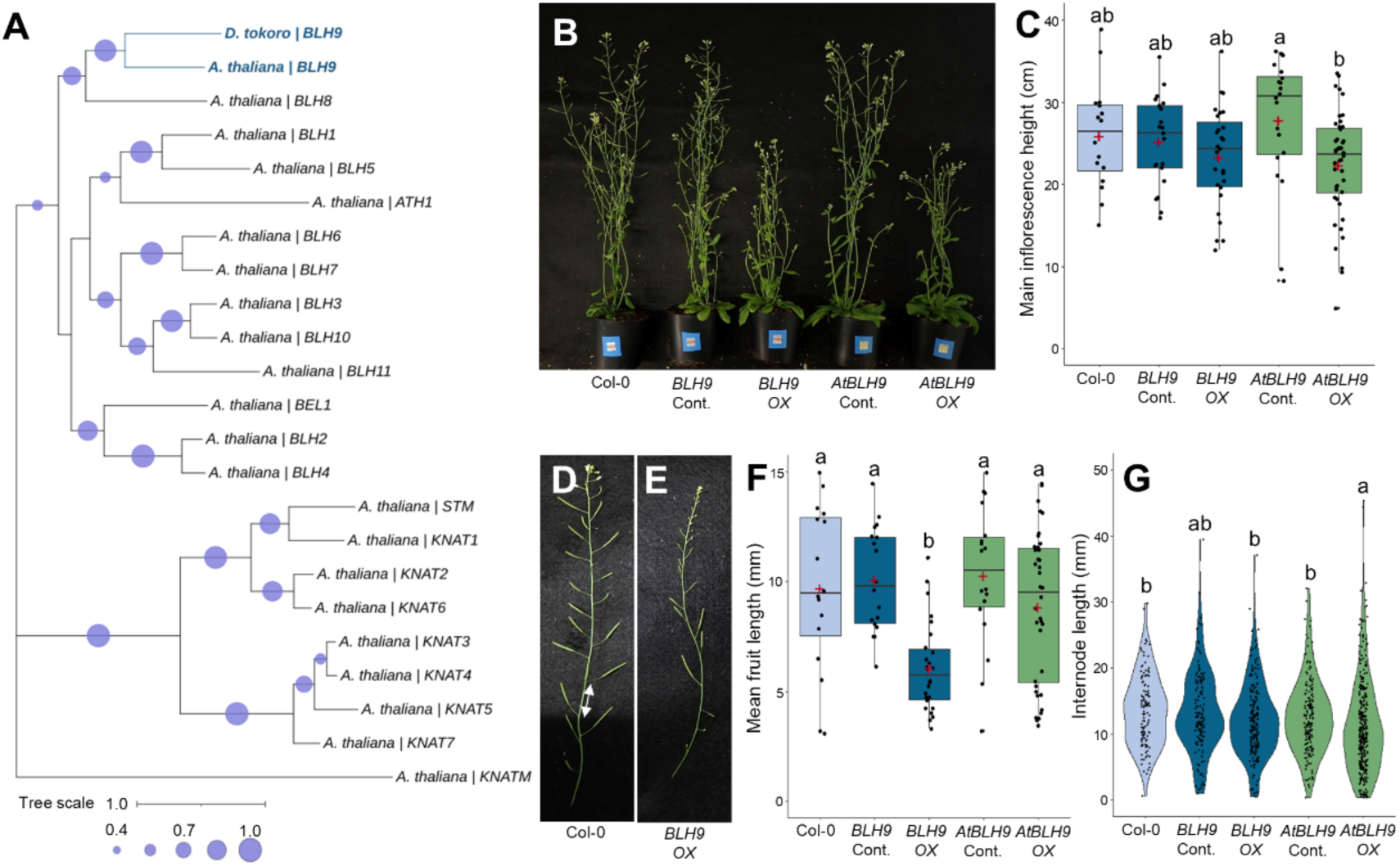
The *D. tokoro Y*-specific gene *BLH9* plays similar roles to *AtBLH9* in inflorescence development. **(A)** Phylogenetic tree of *BLH9* from *D. tokoro* and TALE superfamily proteins from *Arabidopsis thaliana.* The circles in the tree nodes indicate bootstrap values. Tree scale indicates amino acid substitutions per site. **(B)** Whole-plant phenotypes of Col-0, control (Cont.) and overexpression (*OX*) lines of *BLH9,* and control (Cont.) and overexpression (*OX*) lines of *AtBLH9.* Relative expression levels were calculated using the 2^−ΔΔCT^ method, with gene expression in the overexpression lines (*OX*) defined as 2^−ΔΔCT^ > 2 and low expressing lines defined as 2^−ΔΔCT^ ≤ 2 (Fig S17). The low expressing lines were used as control (Cont.). **(C)** Main inflorescence heights of the five lines. Red plus symbol, average; center line, median; box limits, upper and lower quartiles; whiskers, 1.5x interquartile range; points, each measurement. **(D)** Inflorescence of Col-0. White arrow indicates an internode. **(E)** Inflorescence of the *BLH9* overexpression line. **(F)** Mean fruit lengths of the five lines. **(G)** Variation in internode length of the five lines.

We evaluated the relative expression levels of *BLH9* and *AtBLH9* in all T_2_ plants by RT-qPCR using the 2^−ΔΔCT^ method (Fig S17). The T_2_ plants were categorized into two groups: the low-expressing (2^−ΔΔCT^ ≤ 2; indicated as Cont.) and the overexpressing (2^−ΔΔCT^ > 2; indicated as *OX*) groups. Compared to Col-0 plants and the low-expressing T_2_ plants (*BLH9* Cont. and *AtBLH9* Cont.), the T_2_ overexpressing plants (*BLH9 OX* and *AtBLH9 OX*) showed shorter average of main inflorescence height, but the differences between *BLH9 OX* and Col-0 plants and that between *BLH9 OX* and *BLH9* Cont. were not statistically significant (Fig 5B, C). *BLH9 OX* had shorter fruits than the Col-0 plants and the *BLH9* Cont. (Fig 5D, E, F). We also evaluated irregular internode elongation by examining variation in internode length in the main inflorescence. A large variation in internode length was observed in *AtBLH9 OX* (Fig 5G), but not in *BLH9 OX.* These results suggest that *BLH9* from *D. tokoro* shares some but not all functions with *AtBLH9* from *A. thaliana* and may be involved in suppression of fruit development.

## Discussion

In this study, we constructed haplotype-resolved genome assemblies and used them to explore the sex determination mechanism of the wild yam *D. tokoro.* We found that this species has a male heterogametic sex-determination system (*XY =* male*, XX =* female) and that the sex determination locus is situated on chromosome 3. This chromosome is differentiated into *X*-specific and *Y*-specific regions spanning ∼3.4 kb and ∼2.5 kb, respectively. Within the *Y*-specific region, we identified two genes, *BLH9* and *HSP90*, as the primary candidate sex-determination genes.

*D. tokoro* contains homomorphic sex chromosomes without microscopically detectable size differentiation [34], suggesting this species might be in an early stage of sex chromosome evolution [6]. Chromosome 3 is differentiated into *X* and *Y* chromosomes with small *X*- and *Y*-specific regions (Fig 2D, E, F). These regions are located in the middle of chromosome 3 in a region with low gene density, and a high frequency of gypsy LTR retrotransposons (Fig 3A, B), indicating that it is a pericentromeric region [54–58]. These results suggest that the initial SDR of *D. tokoro* occurred in a pericentromeric region with suppressed recombination, a pattern reminiscent of several dioecious species: garden asparagus [66], the poplar *P. tremuloides* [67], and the papaya relative *Vasconcellea parviflora* [68]. *D. alata*, a member of section Enantiophyllum, the most derived group within the genus *Dioscorea*, contains a large pericentric inversion in its sex chromosome that includes a ∼7.6 Mb SDR, accounting for ∼44% of the sex chromosome [44]. The sex chromosome of *D. alata* is collinear with half of the sex chromosome of *D. tokoro*, which belongs to section Stenophora, the most ancestral section in the genus [44]. These findings suggest that several *Dioscorea* species initially contained the same SDR as *D. tokoro*, which expanded during species differentiation. In contrast, the SDR of *D. rotundata* is located in the terminal region of chromosome 11 and shows no collinearity with the sex chromosomes of *D. tokoro* or *D. alata* [27,44]. These results point to multiple transitions of the sex determination system within a single genus.

Despite the many studies on sex determination systems in the genus *Dioscorea,* the detailed molecular basis for its sex determination has not been reported. In this study, we identified two *Y*-specific genes, *BLH9* and *HSP90,* as candidate sex-determination genes related to the regulation of female and male floral organ development. The developmental program leading to carpel initiation is induced by one of the floral homeotic genes *AGAMOUS* (*AG*) [69]. The newly identified gene *BLH9* shares sequence similarity with *AtBLH9* of *A. thaliana* which functions as a repressor of *AG* together with *LEUNIG* and *SEUSS* [70]. *AtBLH9* plays multiple roles in inflorescence architecture and fruit development [60–64]. *AtBLH9-*overexpressing plants have reduced inflorescence height and irregular inflorescence architecture [65]. We also observed reduced inflorescence height and irregular internode elongation in *A. thaliana AtBLH9* overexpression lines (Fig 5B, C, G). The short fruit length phenotype occurred in overexpression lines of *BLH9* (Fig 5D, E, F). Therefore, similar to AtBLH9, the *Y*-encoded BLH9 of *D. tokoro* might function as a repressor of *AG* and suppress female development in *D. tokoro*. Future research should aim to identify the mechanism downstream of *BLH9* expression that controls dioecious floral formation.

Another candidate gene, *HSP90*, shares sequence similarity with the *A. thaliana* chaperone gene *AtHSP90.4.* In *A. thaliana,* AtHSP90.1 to AtHSP90.4 function in the nucleus and cytoplasm and share high sequence similarity [71]. AtHSP90 forms a complex with BRASSINOSTEROID INSENSITIVE 1-EMS-SUPRESSOR 1 (BES1) and BRASSINAZOLE RESISTANT 1 (BZR1) in the nucleus and cytoplasm [72–74], which is required for their downstream function [71]. BZR proteins, including BES1 and BZR1, are indispensable for pollen production; the simultaneous knockout of *BES1*, *BZR1*, and their homologs resulted in male sterility [75]. The *Y*-encoded gene *HSP90* of *D. tokoro* might also be required for successful male flower development.

The identification of two primary candidate genes for sex determination in *D. tokoro* support a major hypothesis for the evolution of dioecy: Two sequential mutations affecting male and female fertility led to the evolution of separate sexes controlled by *Y*-linked maleness-promoting and femaleness-suppressing factors. This two-factor model is also supported by studies in multiple dioecious plants, including garden asparagus and kiwifruit (*Actinidia chinensis*). In garden asparagus, *SUPPRESSOR OF FEMALE FUNCTION* (*SOFF*) and *TAPETAL DEVELOPMENT AND FUNCTION1* (*aspTDF1*) on the *Y* chromosome independently act to suppress pistil development and promote anther development, respectively [17–20]. Kiwifruit also employs a two-gene system for sex determination: *Shy Girl* (*SyGl*) and *Friendly boy* (*FrBy*) independently suppress feminization and promote male sex determination in tapetal cells, respectively [76,77]. The candidate *D. tokoro* genes identified in the current study belong to different classes of genes previously identified in dioecious plants, suggesting that mutations in various genes related to flower development led to the evolution of dioecy.

Key genomic regions and genes for sex determination have been identified in various plant species in recent years; however, the downstream signaling mechanisms resulting in the differential development of female and male floral organs remain elusive [78]. The *Y*-specific region in *D. tokoro* identified in this study includes genes that are upregulated in male flowers during late development, e.g., gene no. 25 in Fig 4E. This gene shares sequence similarity with *A. thaliana EXO70C1*, encoding a component of the exocyst complex (Table 3). *EXO70C1* is expressed in pollen and functions in pollen tube growth [79]. Following sex determination conferred by *BLH9* and *HSP90* expression, *Y*-specific downstream genes such as *EXO70C1* might function in the completion of successful male fertility. The *Y*-specific region of *D. tokoro* also includes genes that are highly expressed in non-reproductive organs (Fig 4E); these genes might be involved in the development of as-yet-unrecognized male-specific characteristics. Further study of these *Y*-specific or *X*-specific genes should help reveal the downstream signaling mechanisms that function after sex determination and play roles in the evolution of sex chromosomes in dioecious plants.

## Conclusion

In summary, the reconstructed *X*- and *Y*-specific regions of chromosome 3 in *D. tokoro* suggest that the SDR of this species is present in the pericentromeric region of chromosome 3. We identified the *Y*-encoded genes *BLH9* and *HSP90* located in the SDR as prime candidates for genetic components underlying sex determination in *D. tokoro*. These two candidate genes are highly expressed during the early stages of male flower development and are predicted to suppress femaleness and promote maleness. Further molecular analyses of sex-determination components in other *Dioscorea* species will shed light on the complex evolution of dioecy in the genus *Dioscorea*.

## Materials and Methods

### Plant materials

To construct the reference genome, a female *D. tokoro* plant (Waka1; original code: DT49) collected from Tahara, Wakayama Pref., Japan (33°32’16.8”N, 135°51’36.0”E) was crossed with a male *D. tokoro* plant (Kita1; original code: 110628-5) collected from Waga-Sennin, Kitakami, Iwate Pref., Japan; 39°17’42.0”N (140°53’45.6”E). The 186 F_1_ progeny comprised 38 female plants, 89 male plants, and 59 non-flowering plants. To identify sex-linked regions, female and male individuals were collected from three wild populations: a population in northern Japan (KTKM; Kitakami, Iwate Pref., Japan; 39°18’25.0”N, 140°54’07.0”E); a population in central Japan (SHG; Koka, Shiga Pref., Japan; 34°56’24.1”N, 136°13’05.3”E); and a population in southern Japan (FKOK; Kasuya, Fukuoka Pref., Japan; 33°38’08.7”N, 130°30’38.2”E). For transcriptome analysis, RNA-seq data were collected from 18 tissue samples. Male and female flowers were collected from wild populations in Takizawa and Kitakami, Iwate Pref., Japan, and non-reproductive organs were collected from Kita1. For small RNA-seq, 15 samples were collected from the wild population in Koka, Shiga Pref., Japan. To check the male specificity of the candidate genes, five females and five males were collected from two wild populations: one in northern Japan (HNMK; Hanamaki, Iwate Pref., Japan; 39°22’10.0”N 141°09’16.0”E) and one in southern Japan (KMMT; Kumamoto, Kumamoto Pref., Japan; 32°53’34.8”N 130°39’22.7”E).

### DNA/RNA extraction, library preparation, and sequencing

For Oxford Nanopore Technologies (ONT) sequencing, genomic DNA was extracted from fresh leaves of Waka1 (female) and Kita1 (male) using NucleoBond HMW DNA (Macherey-Nagel, Düren, Germany). The DNA was subjected to size selection and purification with Short Read Eliminator XL (Circulomics, Baltimore, MD, USA). Long-read libraries for Waka1 (female) were constructed using a Ligation Sequencing Kit (SQK-LSK109; Oxford Nanopore Technologies, Oxford, UK). Long-read libraries for Kita1 (male) were constructed using a Ligation Sequencing Kit V14 (SQK-LSK114). The Waka1 (female) libraries were sequenced using a MinION Mk1C device, and the Kita1 (male) libraries were sequenced using a PromethION 2 Solo device. The raw sequencing data were subjected to base calling using Guppy v6.1.1 [80] for Waka1 (female) and dorado v0.8.1 for Kita1 (male).

For Illumina sequencing, genomic DNA was extracted from Waka1 (female) and Kita1 (male) using a NucleoSpin Plant II Kit (Macherey-Nagel). Libraries for Waka1 (female) were constructed using a Collibri™ ES DNA Library Prep Kit for Illumina Systems (Invitrogen, Camarillo, CA, USA) and a TruSeq DNA PCR-Free LT Library Prep Kit (Illumina, San Diego, CA, USA). Libraries for Kita1 (male) were constructed using a TruSeq DNA PCR-Free LT Library Prep Kit. The libraries were sequenced using the MiSeq and HiSeq X systems. Genomic DNA was extracted from female and male individuals from northern, central, and southern Japan using a DNeasy Plant Maxi Kit (Qiagen, Hilden, Germany). The DNA was purified by phenol/chloroform extraction and ethanol precipitation. Sequencing libraries were constructed using a Collibri™ ES DNA Library Prep Kit for Illumina Systems and sequenced using the HiSeq X system.

RAD-seq was performed as previously described [27]. Genomic DNA was extracted from fresh leaves of Waka1, Kita1, and 186 F_1_ individuals using a NucleoSpin Plant II Kit (Macherey-Nagel). The DNA was digested with the restriction enzymes PacI and NlaIII and used to prepare libraries, and 75-bp paired-end reads were sequenced on the Illumina NextSeq 500 platform. Adapters and unpaired reads were removed using FaQCs and PRINSEQ lite. The filtered RAD-seq reads were used for linkage mapping and association analysis (Data S1).

The RNA-seq data were obtained from 18 samples, including male and female flowers and non-reproductive organs of *D. tokoro.* The samples included male and female flowers at five stages of development: inflorescence stage 0, inflorescence stage 1, inflorescence stage 2, buds, and flowers. The samples also included eight non-reproductive organs: the vegetative shoot apex, leaf, stem, root apex, rhizome root, rhizome stem, rhizome bud, and rhizome storage tissue. Total RNA was extracted from the samples using a RNeasy Plant Mini Kit (Qiagen). The cDNA library was constructed using a TruSeq RNA Sample Prep Kit V2 (Illumina) and sequenced on the Illumina NextSeq 500 platform.

Small RNA-seq data were obtained from 15 samples, including male and female flowers and non-reproductive organs of *D. tokoro.* The samples included male and female flowers at five stages of development: inflorescence stage 0, inflorescence stage 1, inflorescence stage 2, buds, and flowers. The samples also included three non-reproductive organs: the vegetative shoot apex, male and female leaves, and male and female stems. Total RNA was extracted from the samples using Ambion Plant RNA Isolation Aid (Ambion, Austin, TX, USA), and the small RNA fraction was isolated using a mirVana miRNA Isolation Kit (Ambion). The small RNA libraries were constructed using a NEBNext Multiplex Small RNA Library Prep Set for Illumina (New England BioLabs, Ipswich, MA, USA) and DNA Clean & Concentrator-5 (Zymo Research, CA, USA). The libraries were sequenced on the NovaSeq 6000 platform.

### Reference assembly

The genome size of *D. tokoro* Kita1 was estimated by flow cytometry using nuclei prepared from fresh leaf samples. *D. rotundata* accession TDr96-F1 (570 Mb) [27] was used as an internal reference standard. DNA from isolated nuclei was stained with propidium iodide (PI) and analyzed using a Cell Lab Quanta SC Flow Cytometer (Beckman Coulter, USA). The genome size of *D. tokoro* was estimated to be ∼388 Mb (570 Mb × 0.68) (Fig S2).

To construct the reference genome based on the predicted genome size, the ONT sequencing data for Waka1 (female) were filtered using NanoLyse v1.2.0 [81] and Nanofilt v2.8.0 [81]. The ONT sequencing data for Kita1 (male) were filtered using chopper v0.8.0 [82] (Table SM7). The filtered reads from Waka1 (female) were assembled with Flye v2.9.2 [83], resulting in 1,880 contigs with N50 of 1,049,091 bp and a total size of 438.1 Mb. The filtered reads from Kita1 (male) were assembled using PECAT [47], resulting in 128 contigs with N50 of 33,851,599 bp and a total size of 415.0 Mb for haplotype 1 and 415 contigs with N50 of 1,379,312 bp and a total size of 300.8 Mb for haplotype 2. The three assemblies were polished using Racon v1.5.0 [84] and Medaka v1.7.2 (Oxford Nanopore Technologies, 2018). The three assemblies were also polished twice with Hypo v1.0.3 [85] using Illumina short reads from Waka1 (female) and Kita1 (male) filtered using FaQCs v2.08 [86].

### TE and gene annotation

To annotate repetitive sequences, *de novo* repeat libraries were constructed for each assembly using EDTA (The Extensive de novo TE Annotator) v2.1.0 [87]. Using the resulting files, each genome FASTA file was soft-masked with the perl script make_masked.pl provided in EDTA.

For transcriptome-based gene identification using RNA-seq data from 18 samples of *D. tokoro*, poly(A) sequences and reads shorter than 50 bp were removed from the raw RNA-seq reads with FaQCs v2.10 [86]. Low-quality bases on the ends of reads with an average quality score of <20 were trimmed using PRINSEQ lite v0.20.4 [88]. Low-quality reads with an average read quality score of <20 were also removed using PRINSEQ lite v0.20.4. The filtered RNA-seq reads were aligned to the assembled contigs with HISAT2 v2.2.1 [89]. The aligned transcriptomes were assembled using StringTie v3.0.0 [90], and ORF regions were identified using TransDecoder v5.5.0 [91]. Gene models were excluded from further analysis if their coding sequences (CDS) contained an internal stop codon, if the CDS lengths were not multiples of three, or if they encoded proteins shorter than 50 amino acids.

BRAKER2 v2.1.6 [92] was used for *ab initio* gene prediction using the filtered RNA-seq data from *D. tokoro* and protein homology information for *D. alata* and *D. rotundata*. Protein sequences of *D. alata* TDa95/00328 and *D. rotundata* TDr96_F1 were downloaded from NCBI (accession number GCA_020875875.1 and GCF_009730915.1, respectively). The predicted genes were categorized into three groups: gene models fully supported by protein hints, gene models at least partially supported by protein hints, and gene models without any support using the python script selectSupportedSubsets.py in BRAKER. Gene models fully supported by protein hints were selected, and gene models in which the CDS contained an internal stop codon, was comprised of a number of nucleotides that was not a multiple of three, or encoded a protein of <50 amino acids were removed.

Finally, all predicted genes were merged using GffCompare v0.12.6 [93]. The final annotations include 37,712 genes in the Waka1 (female) assembly, 36,483 genes in the Kita1 (male) haplotype 1 assembly, and 28,345 genes in the Kita1 (male) haplotype 2 assembly. To evaluate the completeness of the gene sets in the final scaffolds, BUSCO (Bench-Marking Universal Single Copy) v5.3.2 [46] was used with “genome” as the assessment mode and Embryophyta odb10 as the database (Table S2).

### Generation of chromosomes using the pseudo-testcross approach

The pseudo-testcross approach [48] was used to construct chromosomes from the assembled contigs. SNP-type heterozygous markers and presence/absence-type heterozygous markers were obtained from RAD-seq data from Waka1, Kita1, and 186 F_1_ individuals. Parental-line-specific heterozygous markers were identified as previously described [29] with several modifications (S1 Supplementary Materials and Methods). Linkage maps were constructed based on the heterozygous SNP-type and heterozygous presence/absence-type markers. For each marker set (female-parent-heterozygous marker set and male-parent-heterozygous marker set), the markers were converted to genotype-formatted data to construct genetic linkage maps using MSTmap v1.0 [94]. After trimming the orphan linkage groups, linkage groups and ordered markers in each linkage group were reconstructed using R/qtl [50]. Finally, two parental-specific linkage maps were constructed and visualized using Asmap [95]. Based on the two parental-specific linkage maps, the contigs were anchored and linearly ordered as chromosomes using ALLMAPS [96]. Gene-order-based synteny of the three constructed reference genomes was detected using MCscan [97].

### Identification of sex-linked regions by association analysis and coverage analysis

To identify sex-determination systems and sex-linked regions, association analysis was conducted using RAD-seq data from the 127 flowering F_1_ progeny, which segregated into 38 females and 89 males. The RAD-seq data were aligned to the newly constructed female and male chromosomes using BWA v0.7.18-r1243-dirty [98]. SNP-type markers and presence/absence-type markers were identified based on these alignments. The presence/absence-type markers were defined based on the aligned read depth of F_1_ individuals following a cutoff of depth ≥ 3 indicating presence and depth = 0 indicating absence.

The associations between the genotypes and sex phenotypes of the 127 flowering F_1_ individuals were calculated using Fisher’s exact test. The *q*-value of Fisher’s exact test was obtained for each marker by comparing the frequencies of particular alleles and sex phenotypes categorized as female or male. The false discovery rate (FDR) was set to 0.05 and was corrected by Benjamini-Hochberg correction. The log-transformed *q*-values (– log10(*q*)) for each position were visualized as Manhattan plots. The associations between the genotypes and sex phenotypes were also checked by composite interval mapping (CIM) using R/qtl [50].

Regions with major gene losses on each sex chromosome were detected by coverage analysis using Illumina short reads from four combinations of male and female individuals: male parent Kita1 and female parent Waka1 and male and female individuals collected from northern, central, and southern Japan. The Illumina short reads were filtered using FaQCs v2.10 [86] and aligned to each newly constructed reference genome: Waka1 (female), Kita1 (male) haplotype 1, and Kita1 (male) haplotype 2. The alignment was conducted using BWA v0.7.17-r1188 [98]. The alignment depths were calculated with Samtools v1.16.1 [99]. The alignment depths were normalized by dividing the values with the mean depth of all positions on the chromosomes. To reduce the effects of error on the alignment depths, a sliding window approach was employed (window size = 150 kb, step size = 10 kb). *X*- and *Y*-specific regions were then identified using the region depths. Regions in which the male depth was 0.5 (half of mean normalized depth) ± 0.15 and the female depth was 1 (mean normalized depth) ± 0.15 were selected as *X*-specific regions. Regions in which the male depth was 0.5 (half the mean normalized depth) ± 0.15 and the female depth was <0.15 were selected as *Y*-specific regions.

### Genome structures of the *X* and *Y* chromosomes

Gene-order-based synteny of chromosomes 3*X* and 3*Y* was detected using MCscan [97]. Gene density and repetitive sequence accumulation values were obtained from gene annotation and the predicted repetitive sequences of the Waka1 (female) assembly and the Kita1 (male) haplotype 1 assembly. Repetitive sequences included Copia LTR retrotransposons, Gypsy LTR retrotransposons, hAT TIR transposons, and Helitrons predicted by EDTA v2.1.0. Gene density and retrotransposon accumulation were visualized using the jcvi package [97].

To evaluate recombination suppression around the SDR in the *D. tokoro* genome, SNP-type heterozygous markers were obtained from RAD-seq data from Waka1, Kita1, and 186 F_1_ individuals. The RAD-seq markers were aligned to the Kita1 (male) haplotype 1 and Waka1 (female) reference genomes. Male-parent-heterozygous SNP markers and female-parent-heterozygous SNP markers showing 1:1 segregation of homozygous and heterozygous genotypes in the F_1_ progeny were selected. After marker selection, linkage distances were obtained based on genetic linkage maps constructed using MSTmap v1.0 [92]. The linkage distances and physical distances were compared and visualized using ALLMAPS [96].

### Identification of candidate genes and miRNAs for sex determination

For transcriptome analysis of genes, the filtered RNA-seq data from 18 samples, including male flowers, female flowers, and non-reproductive organs, were aligned to the assembled contigs with HISAT2 v2.2.1 [89], and the aligned reads were counted using the featureCounts function in Subread v2.0.1 [100]. Differential expression analysis was performed with DESeq2 v3.15 [101] for two comparisons: male vs. female flowers at three stages of early development (inflorescence stages 0, 1, and 2); and male flowers at three stages of early development vs. non-reproductive organs. The false discovery rate threshold was set to 0.05.

For transcriptome analysis of miRNAs, small RNA-seq data from 15 samples, including male flowers, female flowers, and non-reproductive organs, were filtered using FaQCs v2.10 [86] and Seqkit v2.3.0 [102]. Based on the small RNA-seq data, miRNAs were predicted using miRDeep-P2 v1.1.4 [103] as described previously [104]. The trimmed small RNA-seq reads were then aligned to the predicted miRNA references with Bowtie v1.3.1 [105]. The aligned reads were counted using the perl script bam2ref_counts.pl [104]. Differential expression analysis was performed with DESeq2 v3.15 [101] to compare male vs. female flowers at three stages of early development (inflorescence stage 0, 1, and 2). The false discovery rate threshold was set to 0.05.

In the first step of candidate gene and miRNA identification, 51 genes and 15 miRNAs located on *Y*-specific regions were identified. In this step, *Y*-specific regions were detected with liberal thresholds in sliding windows (window size = 50 kb, step size = 1 kb). In the second step, within the *Y*-specific genes, 26 genes (and no miRNAs) were detected that were highly expressed during these three stages of early male vs. female flower development (*q*-value < 0.05). Ten genes were also identified that were significantly upregulated in male vs. female flowers during the same three stages of development (log_2_FC > 2). In the third step, within the *Y*-specific genes, 10 genes were identified that were highly expressed in male flowers compared to non-reproductive organs during these three stages (*q*-value < 0.05). Three genes that were significantly upregulated in male flowers compared to non-reproductive organs during the three stages also identified (log_2_FC > 2). Finally, two *Y*-specific genes were selected with significantly upregulated expression in male flower during the early three stages of development (*q*-value < 0.05 and log_2_FC > 2) in both transcriptome comparisons. The similarity of the candidate gene products were compared to known proteins using BLASTX with the Swiss-Prot function. The male-specific expression of the two candidate genes was confirmed by PCR amplification from five females and five males from two wild populations, one from northern Japan and one from southern Japan.

### Overexpressing BLH9 and AtBLH9 in Arabidopsis thaliana

The homologous genes of *BLH9* were subjected to phylogenetic analysis using protein sequences of BLH9 and TALE superfamily members in *A. thaliana.* The sequences were aligned with Molecular Evolutionary Genetics Analysis (MEGA) v11.0.13 [106], and a phylogenic tree was constructed using the maximum likelihood method in MEGA using the Jones-Taylor-Thornton model. The tree was visualized using Interactive Tree Of Life (iTOL) v7.0 [107].

To overexpress *BLH9* and *AtBLH9* in *A. thaliana*, full-length cDNA clones were used for In-Fusion cloning. Total RNA was extracted from *D. tokoro* and *A. thaliana* using a Maxwell RSC Plant RNA Kit (Promega, Madison, WI, USA), and was reverse-transcribed into cDNA using a ReverTra Ace qPCR RT Kit FSQ-101 (TOYOBO, Osaka, Japan). *BLH9* fragment was amplified from the *D. tokoro* cDNA using KOD FX Neo (TOYOBO). *AtBLH9* was amplified from *A. thaliana* cDNA using PrimeSTAR GXL DNA Polymerase (Takara Bio). Each cDNA fragment was cloned into EcoRI and BamHI sites of the pBICP35 binary vector containing the CaMV35S promoter [108] using In-Fusion HD Cloning Kit (Takara Bio). The pBICP35::BLH9 and pBICP35::AtBLH9 were transformed into *Agrobacterium tumefaciens* strain GV3101::pMP90 by electroporation. *A. thaliana* transformation was performed by the floral dip method with *A. tumefaciens* strains carrying the binary expression plasmids, and kanamycin-resistant T_1_ plants were selected. T_2_ lines that showed a 3:1 segregation ratio for kanamycin resistance were selected, as they were thought to contain a single T-DNA.

To examine inflorescence phenotypes, 16 plants each were grown from eight lines: Col-0, three T_2_ lines transformed with the construct for *BLH9*, and four T_2_ lines transformed with the construct for *AtBLH9*. To eliminate the effects of kanamycin selection on plant growth, all seeds of T_2_ plants and Col-0 were grown in Murashige and Skoog (MS) medium without kanamycin. The expression levels of *BLH9* and *AtBLH9* in T_2_ plants were confirmed by RT-qPCR (Data S2). The primers used in RT-qPCR are listed in S1 Supplementary Materials and Methods. The T_2_ plants were separated into four groups based on the relative expression levels of the inserted genes calculated by the 2^−ΔΔCT^ method (Fig S17): *BLH9* Control (2^−ΔΔCT^ ≤ 2), *BLH9 OX* (2^−ΔΔCT^ > 2), *AtBLH9* Control (2^−ΔΔCT^ ≤ 2), and *AtBLH9 OX* (2^−ΔΔCT^ > 2). At 30 days after the plants were transplanted to soil, three phenotypes were measured: inflorescence height, mean fruit length, and internode length. All phenotypic data are listed in Data S3, S4 and S5. The mean inflorescence heights and mean fruit lengths were compared by Wilcoxon rank sum test between the five groups: Col-0, *BLH9* Control, *BLH9 OX*, *AtBLH9* Control, and *AtBLH9 OX*. The variations in internode lengths between the five groups were compared by *F*-test. The *p*-values were adjusted by Bonferroni correction.

All detailed materials and methods are shown in S1 Supplementary Materials and Methods. All sequencing data are available under the Umbrella BioProject number PRJNA1223176. The genome assemblies and annotation files are available at zenodo: 10.5281/zenodo.14899024.

## Acknowledgements

We would like to thank members of the Crop Evolution Laboratory at Kyoto University and Hiroki Matsuo of the Plant Pathology Laboratory at Kyoto University for advice on genome analysis. We also thank Yui Niihara for the support with the experiment performed at Kyoto Sangyo University. We are grateful to Katsuyoshi Kubota and the staff of the Kasuya Research Forest and Kyushu University and the Kosaji villagers in Koka, Shiga Pref., for their cooperation with sample collection. Genome analyses were conducted on the supercomputer at ACCMS, Kyoto University, and the NIG supercomputer at the ROIS National Institute of Genetics.

## Supporting information

### S1 Supplementary materials and methods

1. Materials

2. Reference assembly

3. Generation of chromosomes using pseudo-testcross methods

4. Identification of sex-linked regions by association analysis and coverage analysis

5. Genome structures of the *X* and *Y* chromosomes

6. Identification of candidate genes for sex determination

7. Identification of candidate miRNAs for sex determination

8. Overexpression of *BLH9* and *AtBLH9* in *Arabidopsis thaliana*

**Data S1.** Summary of RAD-seq data from Waka1, Kita1, and 186 F_1_ plants.

**Data S2.** Ct values for all transformed T_2_ plants confirmed by RT-qPCR.

**Data S3.** Main inflorescence lengths of Col-0 and T_2_ plants.

**Data S4.** Fruit lengths of Col-0 and T_2_ plants.

**Data S5.** Internode lengths of Col-0 and T_2_ plants.

---

**Fig S1.**
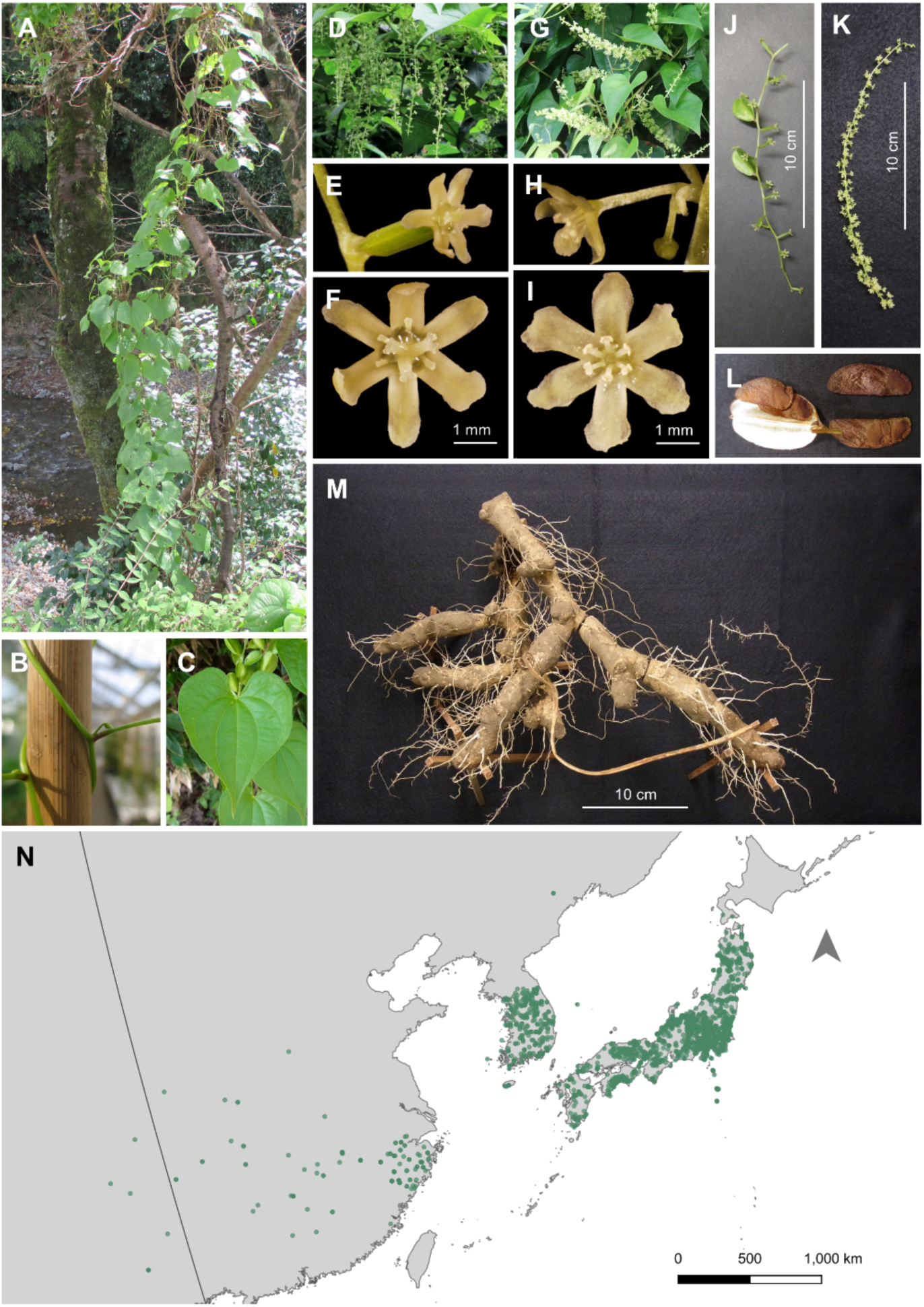
Morphological characteristics of *Dioscorea tokoro*. **(A-C)** Photographs of the perennial herbaceous wild yam *D. tokoro*, including a vine **(A)** with left twining stems **(B)** and heart-shaped alternate leaves **(C)**. **(D-I)** This dioecious species bears female **(D, E, F, J)** and male **(G, H, I, K)** flowers on separate individuals. **(L)** Females produce capsular fruits containing six winged seeds. **(M)** The plant also reproduces via underground rhizomes. **(N)** *D. tokoro* is widely distributed in temperate East Asia: Japan, Korea, and China. The distribution data were provided by the GBIF Occurrence Download (https://doi.org/10.15468/dl.yqn4xt; https://doi.org/10.15468/dl.exatcz). The map was created with Quantum Geographic Information System (QGIS) software version 3.16.0 (QGIS Development Team 2022). The base map was obtained from the Geospatial Information Authority of Japan, 2020.

**Fig S2.**
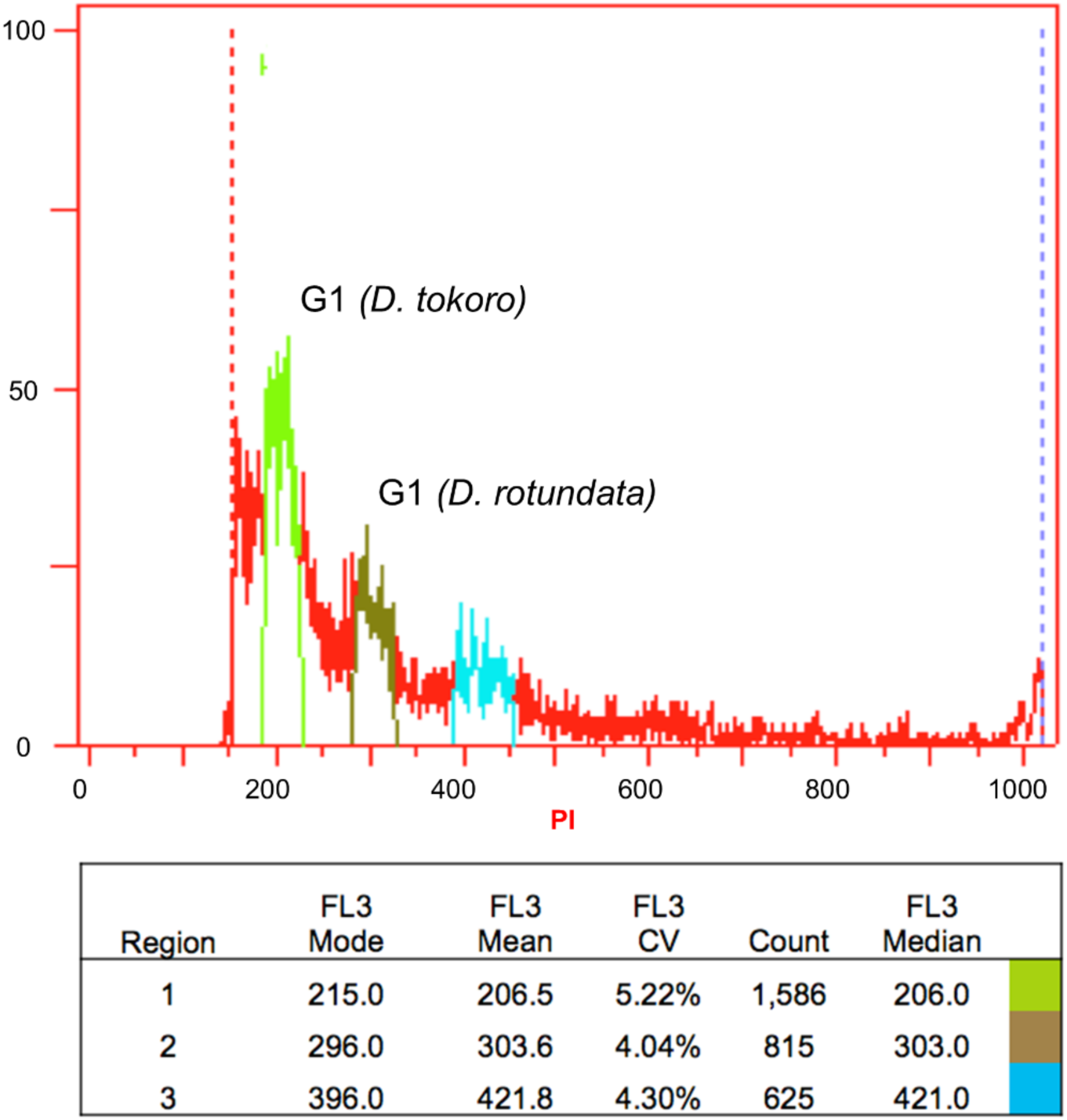
Estimation of genome size by flow cytometry. The genome size of *D. tokoro* was estimated by flow cytometry using nuclei prepared from fresh leaf samples of *D. tokoro* and *D. rotundata* (570 Mb) [27]. Nuclei were isolated and stained with propidium iodide (PI) and analyzed using a Cell Lab Quanta SC Flow Cytometer (Beckman Coulter, USA). The genome size of *D. tokoro* was estimated to be 388 Mb based on the G1 peak mean ratio [*D. tokoro* (206.5):*D. rotundata* (303.6) = 0.680].

**Fig S3.**
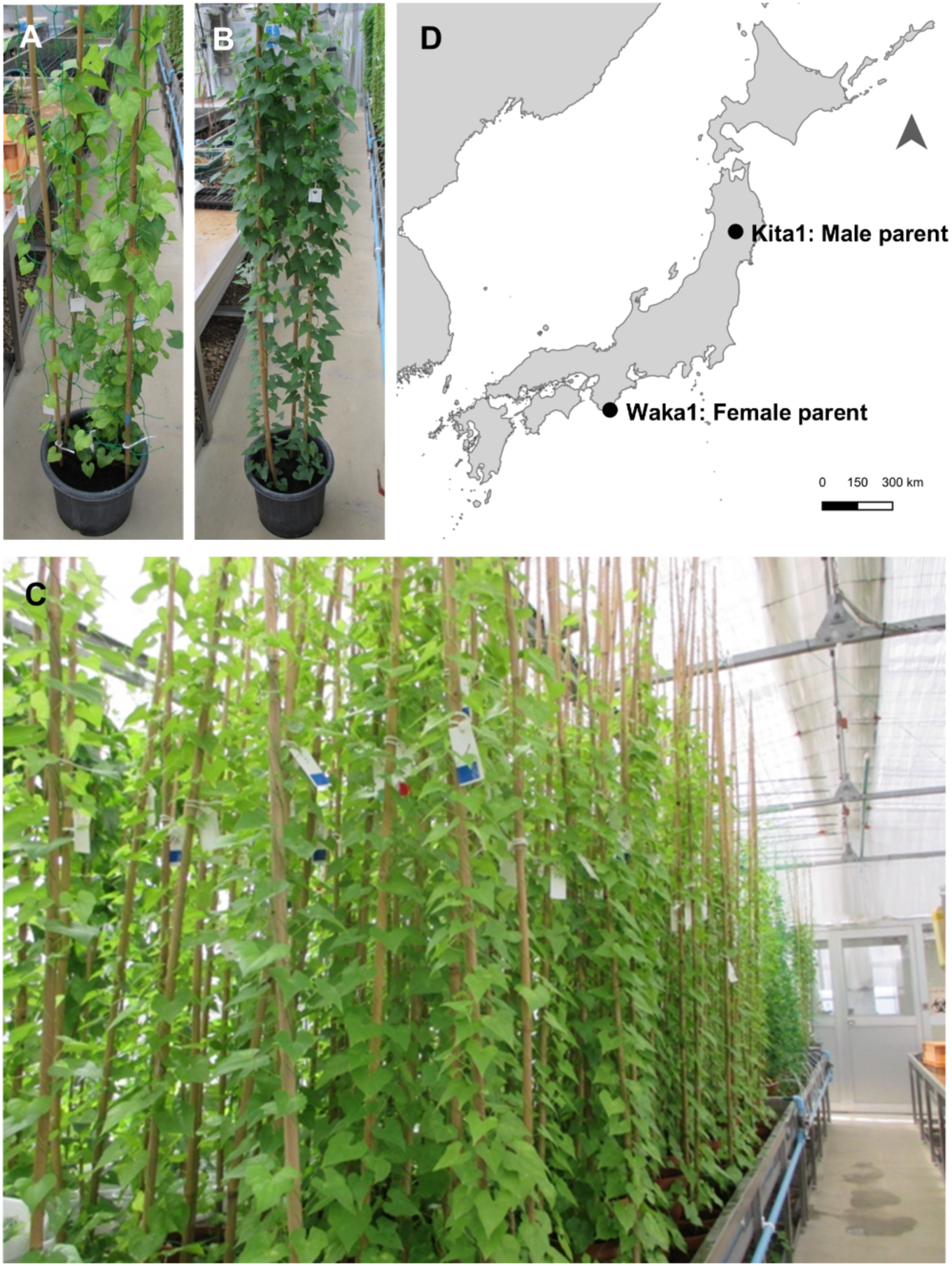
F_1_ progeny used for linkage map construction and linkage analysis. **(A,B)** The 186 F_1_ progeny, comprising 38 females, 89 males, and 59 non-flowering individuals, were derived from a cross between the male parent Kita1 **(A)** and female parent Waka1 **(B)**. **(C)** The F_1_ progeny were grown in Iwate Biotechnology Research Center in Japan. **(D)** The male parent Kita1 was collected from Waga-Sennin in Kitakami, Iwate Pref., Japan. The female parent Waka1 was collected from Tahara, Wakayama Pref., Japan. The map was created with Quantum Geographic Information System (QGIS) software version 3.16.0 (QGIS Development Team 2022). The base map was obtained from the Geospatial Information Authority of Japan, 2020.

**Fig S4.**
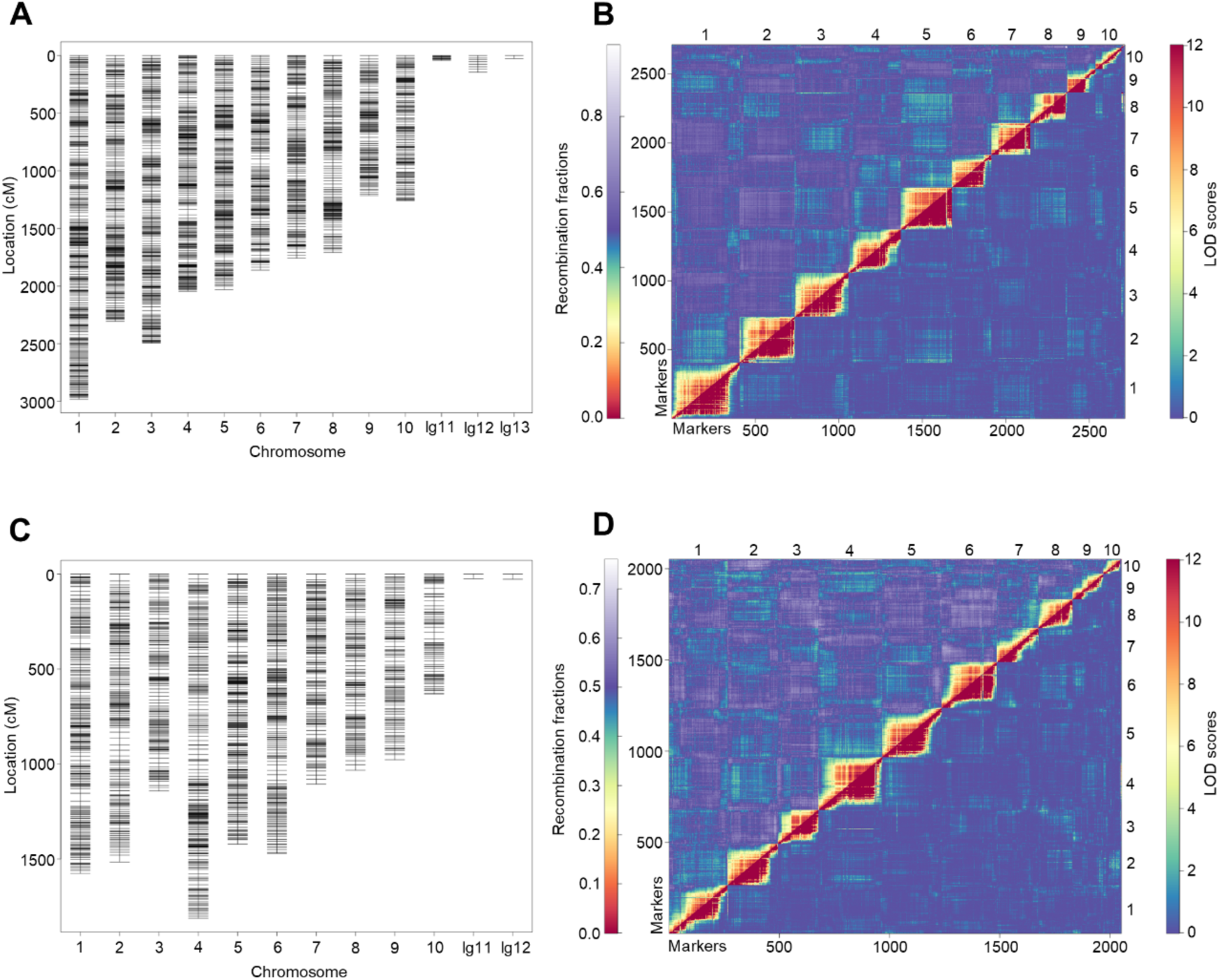
RAD-seq-based linkage map of the Waka1 (female) reference genome generated by the pseudo-testcross method using 186 F_1_ progeny. **(A)** Genetic map based on female-parent-heterozygous markers. **(B)** Plots of estimated recombination fractions and LOD scores for the genetic map based on female-parent-heterozygous markers. **(C)** Genetic map based on male-parent-heterozygous markers. **(D)** Plots of estimated recombination fractions and LOD scores for the genetic map based on male-parent-heterozygous markers.

**Fig S5.**
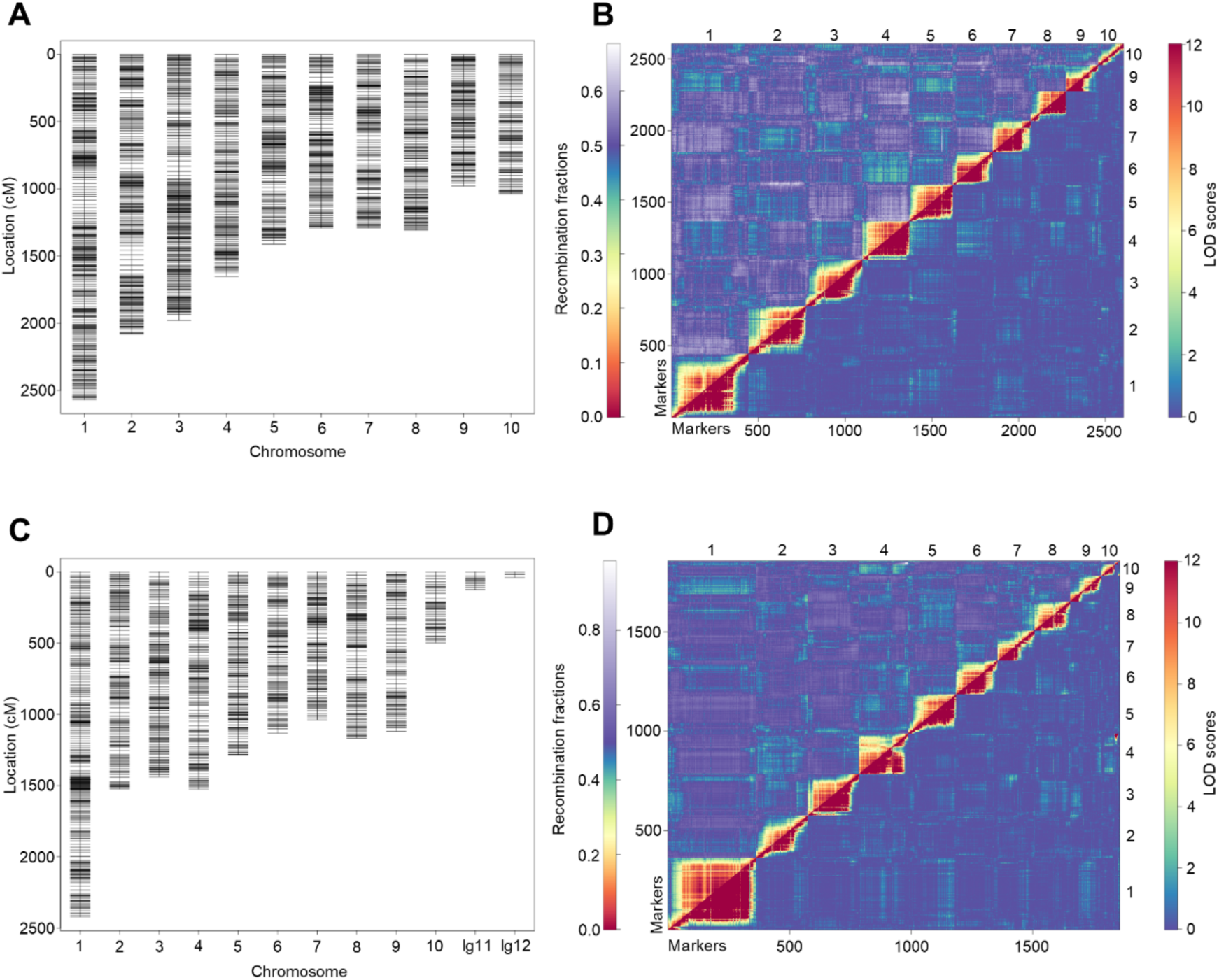
RAD-seq-based linkage map of the haplotype 1 Kita1 (male) reference genome generated by the pseudo-testcross method using 186 F_1_ progeny. **(A)** Genetic map based on male-parent-heterozygous markers. **(B)** Plot of estimated recombination fractions and LOD scores for the genetic map based on male-parent-heterozygous markers. **(C)** Genetic map based on female-parent-heterozygous markers. **(D)** Plot of estimated recombination fractions and LOD scores for the genetic map based on female-parent-heterozygous markers.

**Fig S6.**
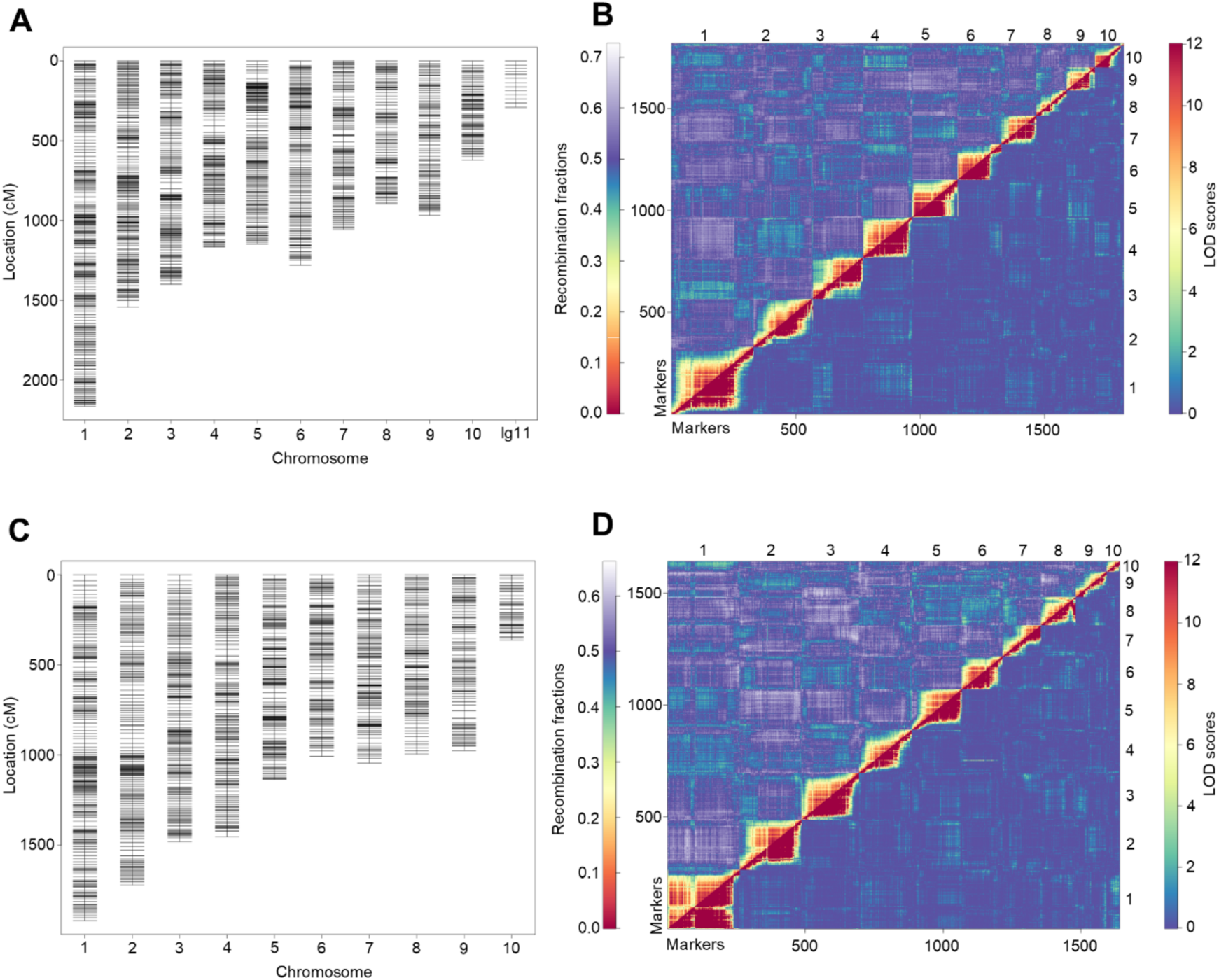
RAD-seq-based linkage map of the haplotype 2 Kita1 (male) reference genome generated by the pseudo-testcross method using 186 F_1_ progeny. **(A)** Genetic map based on male-parent-heterozygous markers. **(B)** Plot of estimated recombination fractions and LOD scores for the genetic map based on male-parent-heterozygous markers. **(C)** Genetic map based on female-parent-heterozygous markers. **(D)** Plot of estimated recombination fractions and LOD scores for the genetic map based on female-parent-heterozygous markers.

**Fig S7.**
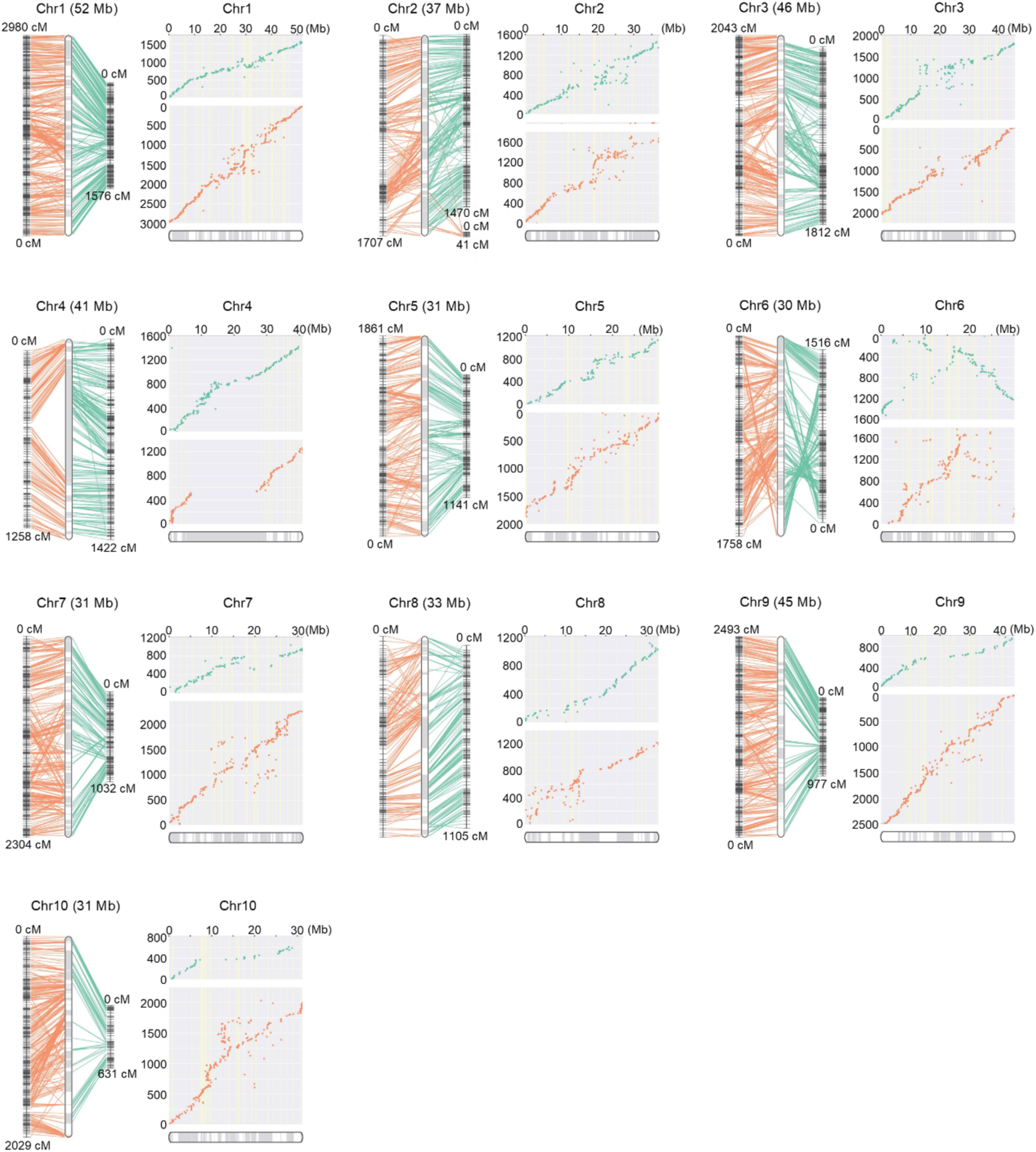
Integrated genetic and physical maps of the Waka1 (female) reference genome. Ten chromosome-scale pseudo-molecules were generated using a RAD-based genetic map generated with 186 F_1_ progeny. For each chromosome, the left panel shows the physical positions on the newly constructed chromosome based on the genetic map using male-parent-heterozygous markers (green) and the genetic map using female-parent-heterozygous markers (orange). The right panels show scatter plots representing the physical positions on the chromosome (*x*-axis) versus the map location (*y*-axis). Scaffold ordering and orientation were conducted using ALLMAPS [96].

**Fig S8.**
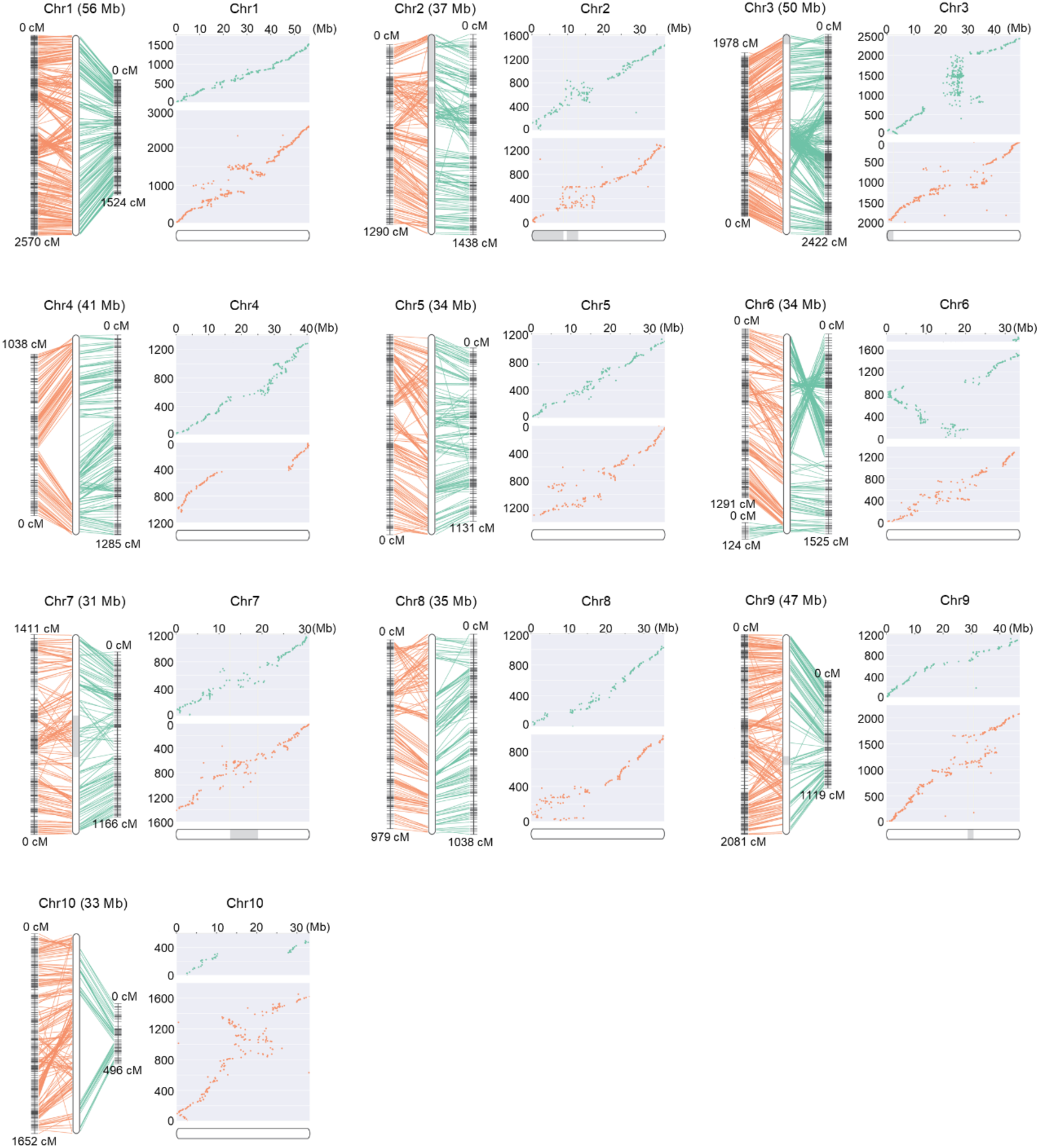
Integrated genetic and physical maps of the haplotype 1 Kita1 (male) reference genome. Ten chromosome-scale pseudo-molecules were generated using a RAD-based genetic map generated with 186 F_1_ progeny. For each chromosome, the left panel shows the physical positions on the newly constructed chromosome based on the genetic map using male-parent-heterozygous markers (green) and the genetic map using female-parent-heterozygous markers (orange). The right panels show scatter plots representing the physical positions on the chromosome (*x*-axis) versus the map location (*y*-axis). Scaffold ordering and orientation were conducted using ALLMAPS [96].

**Fig S9.**
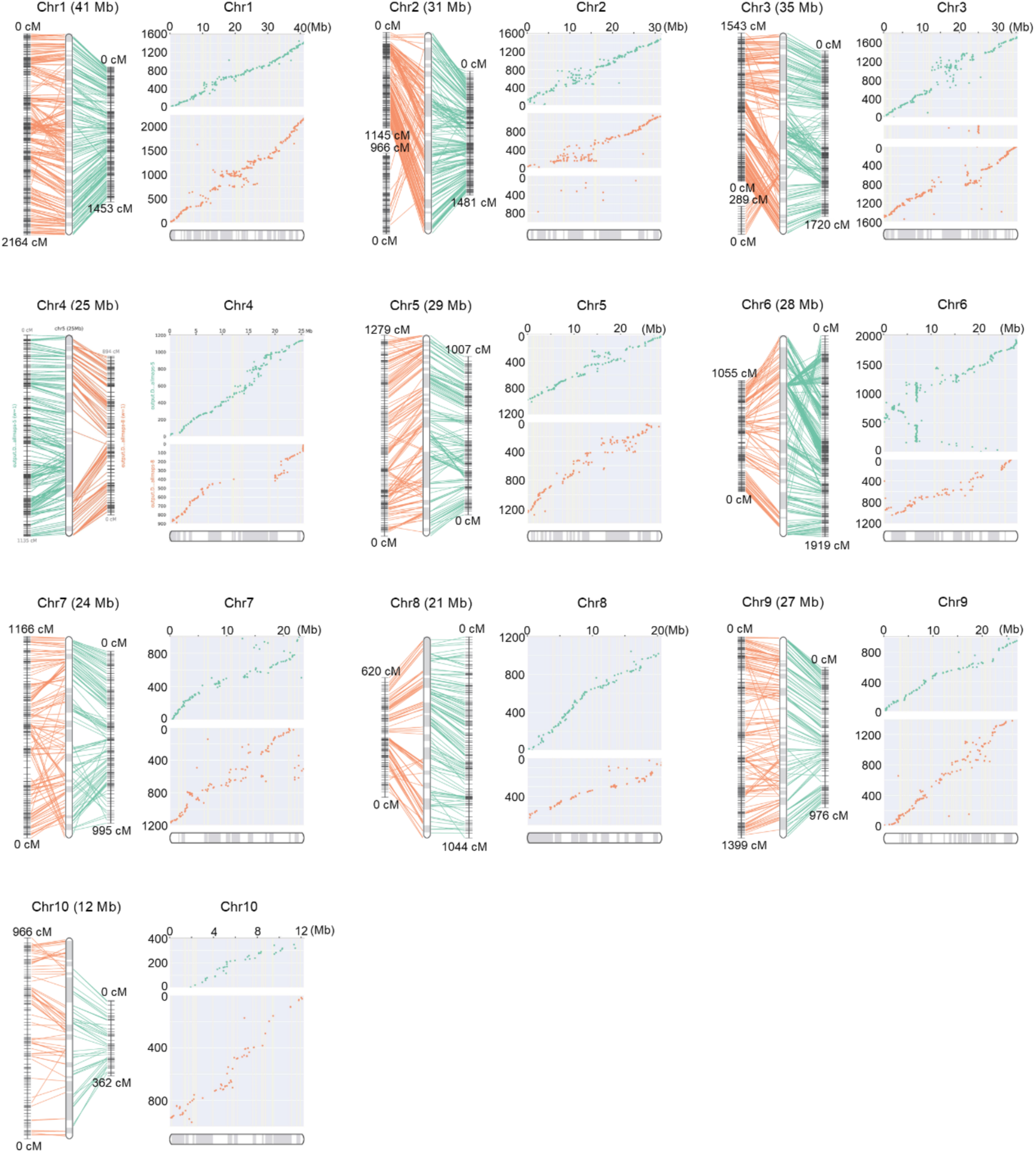
Integrated genetic and physical maps of the haplotype 2 Kita1 (male) reference genome. Ten chromosome-scale pseudo-molecules were generated using a RAD-based genetic map generated with 186 F_1_ progeny. For each chromosome, the left panel shows the physical positions on the newly constructed chromosome based on the genetic map using male-parent-heterozygous markers (green) and the genetic map using female-parent-heterozygous markers (orange). The right panels show scatter plots representing the physical positions on the chromosome (*x*-axis) versus the map location (*y*-axis). Scaffold ordering and orientation were conducted using ALLMAPS [96].

**Fig S10.**
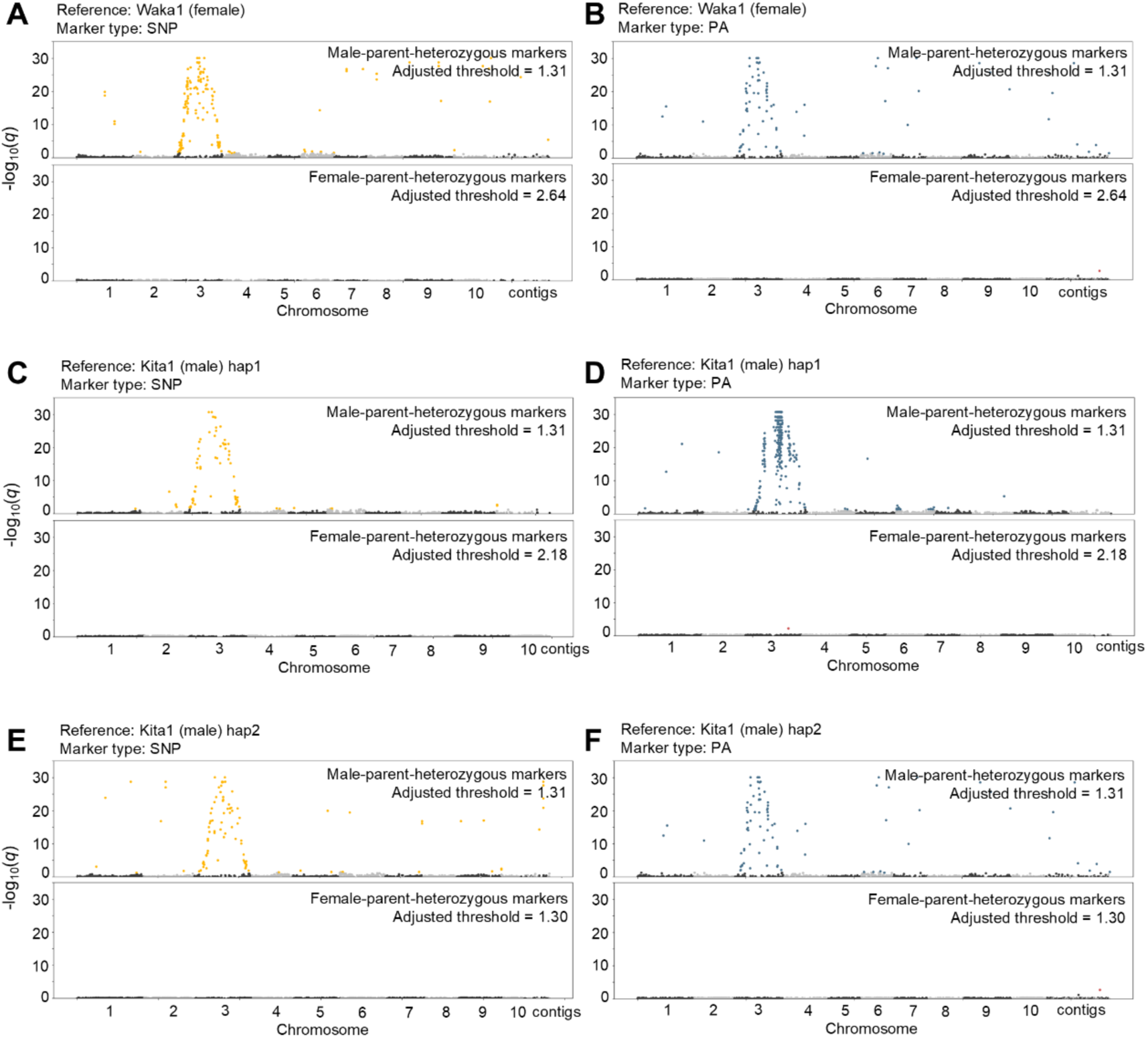
Manhattan plots of –log_10_(*q*) values between markers and sex phenotype showing genomic regions associated with sex phenotype. The –log_10_(*q*) values were obtained by Fisher’s exact test. In each figure, the upper plot was obtained using male-parent-heterozygous markers and the lower plot was obtained using female-parent-heterozygous markers. A significance threshold of 5% false discovery rate was adjusted by Benjamini-Hochberg correction. **(A)** Manhattan plot of –log_10_(*q*) values between SNP-type markers and sex phenotype in the Waka1 (female) assembly. The yellow plots in the upper panels indicate SNP-type markers with significant associations with sex phenotype. **(B)** Manhattan plot of –log_10_(*q*) values between PA markers and sex phenotype in the Waka1 (female) assembly. The blue and red plots indicate PA markers with significant associations with male and female phenotypes, respectively. **(C)** Manhattan plot of –log_10_(*q*) values between SNP-type markers and sex phenotype in the Kita1 (male) haplotype 1 assembly. **(D)** Manhattan plot of –log_10_(*q*) values between PA markers and sex phenotype in the Kita1 (male) haplotype 1 assembly. **(E)** Manhattan plot of –log_10_(*q*) values between SNP-type markers and sex phenotype in the Kita1 (male) haplotype 2 assembly. **(F)** Manhattan plot of –log_10_(*q*) values between PA markers and sex phenotype in the Kita1 (male) haplotype 2 assembly.

**Fig S11.**
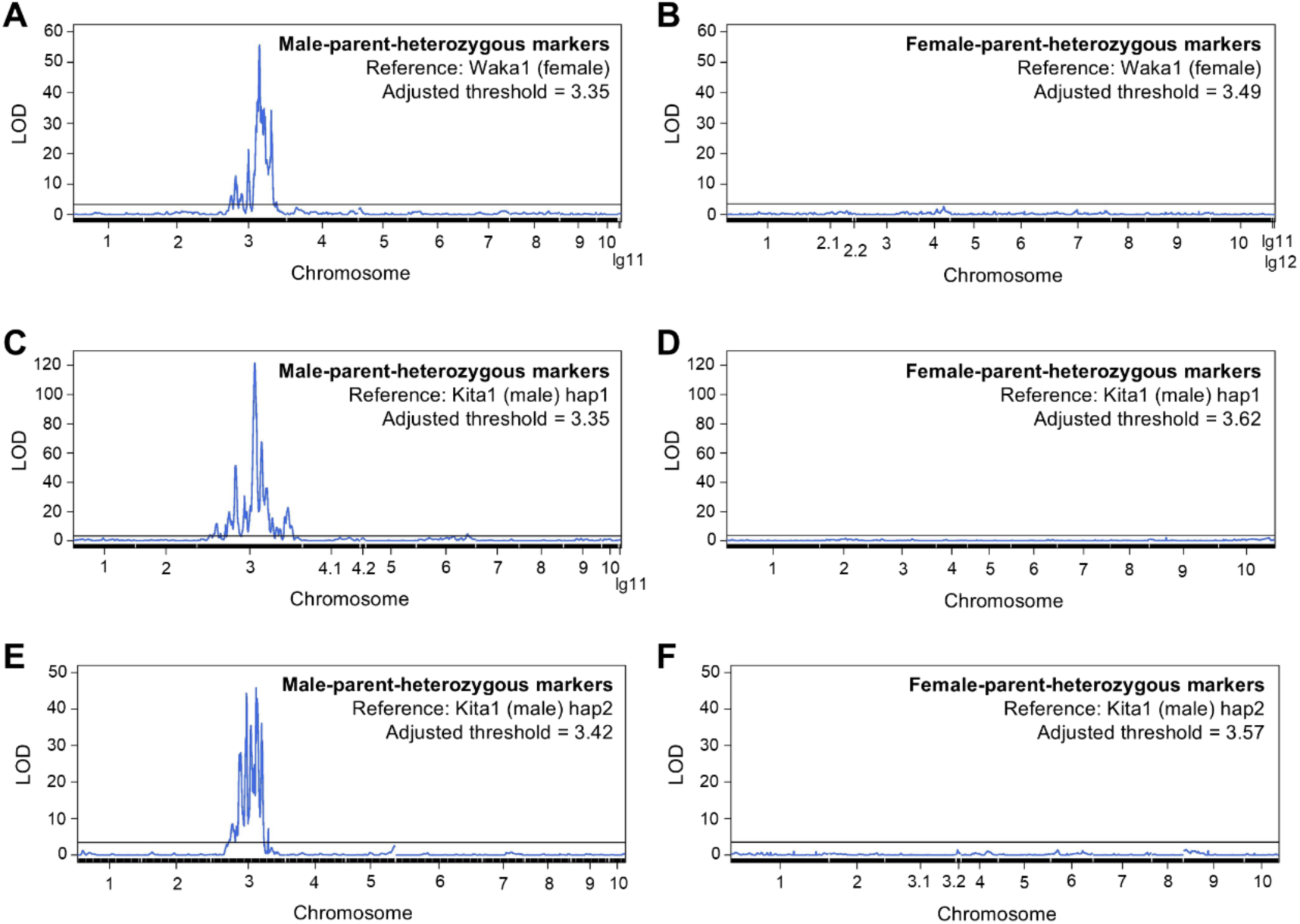
Composite interval mapping (CIM) curves detecting quantitative trait loci (QTL) related to sex phenotype. LOD scores were obtained to detect QTL related to the sex phenotypes of 186 F_1_ progeny and visualized as CIM curves. **(A)** CIM curve of male-parent-heterozygous markers obtained from RAD markers aligned to the Waka1 (female) assembly. **(B)** CIM curve of female-parent-heterozygous markers obtained from RAD markers aligned to the Waka1 (female) assembly. **(C)** CIM curve of male-parent-heterozygous markers obtained from RAD markers aligned to the Kita1 (male) haplotype 1 assembly. **(D)** CIM curve of female-parent-heterozygous markers obtained from RAD markers aligned to the Kita1 (male) haplotype 1 assembly. **(E)** CIM curve of male-parent-heterozygous markers obtained from RAD markers aligned to the Kita1 (male) haplotype 2 assembly. **(F)** CIM curve of female-parent-heterozygous markers obtained from RAD markers aligned to the Kita1 (male) haplotype 2 assembly. The horizontal lines indicate adjusted thresholds.

**Fig S12.**
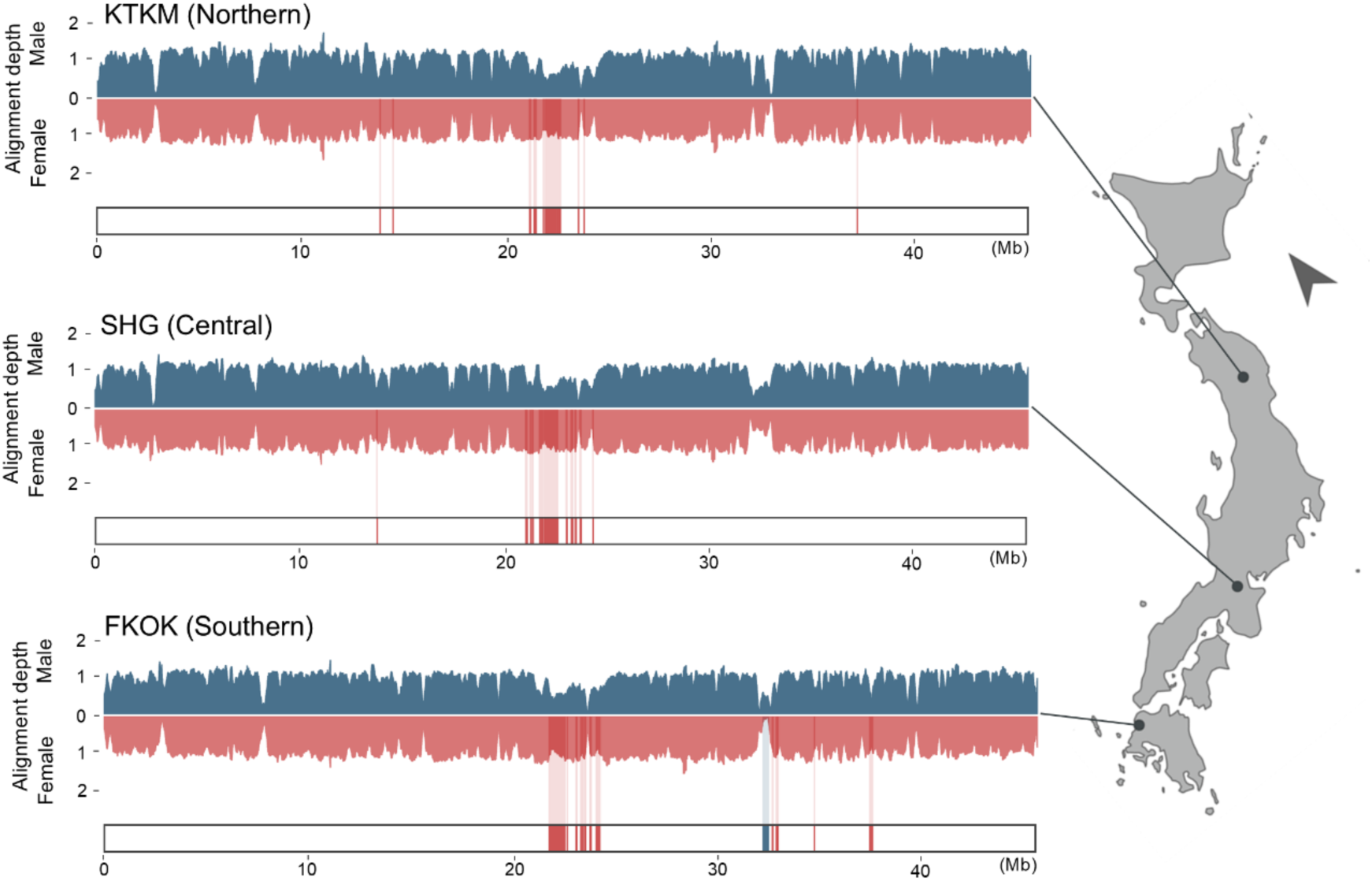
Alignment depths on the *X* chromosomes of male and female individuals from northern, central, and southern Japan. The bar graphs show alignment depths of short reads from male and female individuals. The red regions below the bar graphs show *X-*specific regions. The map was created with Quantum Geographic Information System (QGIS) software version 3.16.0 (QGIS Development Team 2022). The base map was obtained from the Geospatial Information Authority of Japan, 2020.

**Fig S13.**
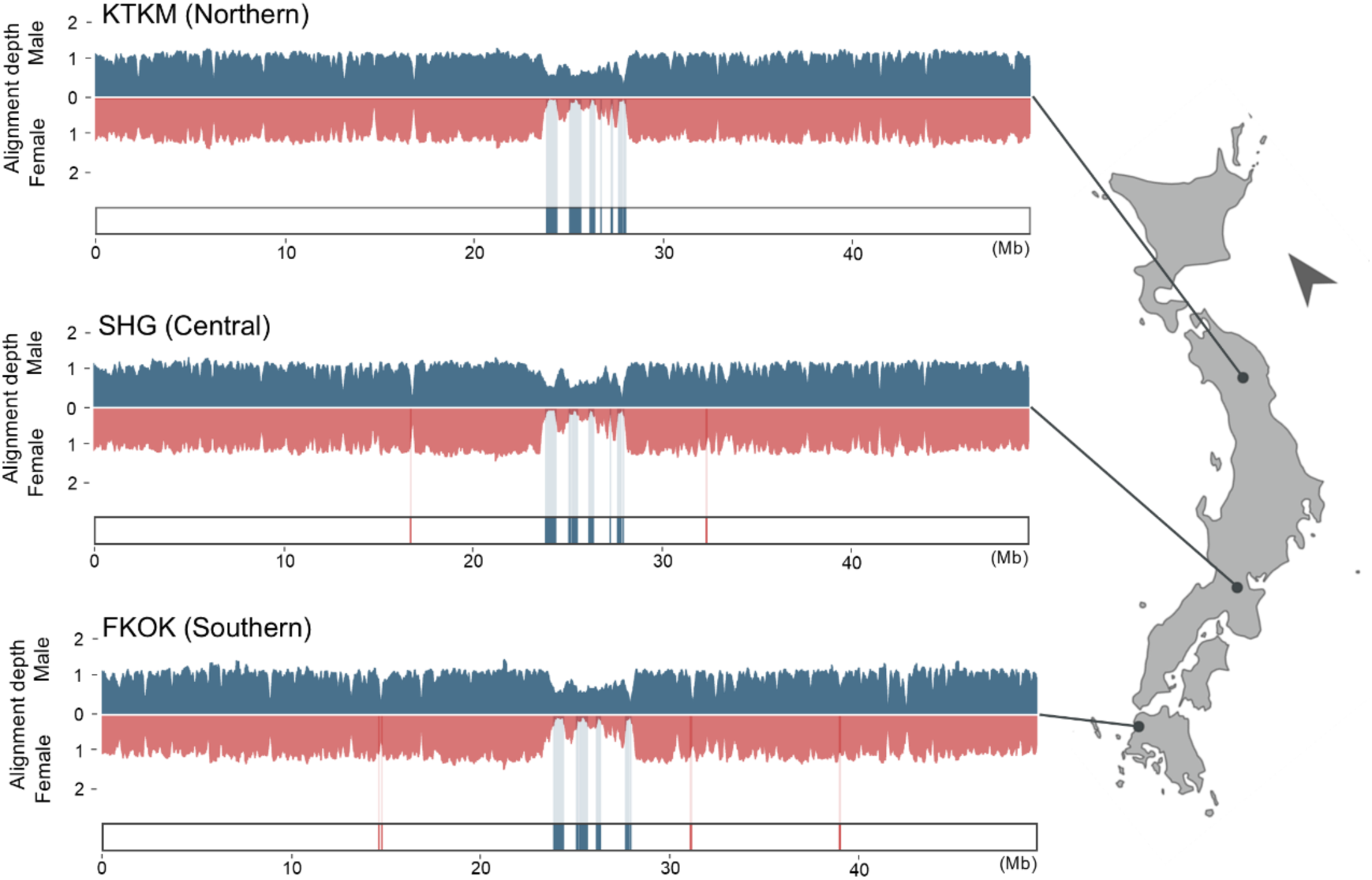
Alignment depths on the *Y* chromosomes of male and female individuals from Northern, Central, and Southern Japan. The bar graphs show the alignment depths of short reads from male parent and female individuals. The blue regions show *Y-*specific regions. The map was created with Quantum Geographic Information System (QGIS) software version 3.16.0 (QGIS Development Team 2022). The base map was obtained from the Geospatial Information Authority of Japan, 2020.

**Fig S14.**
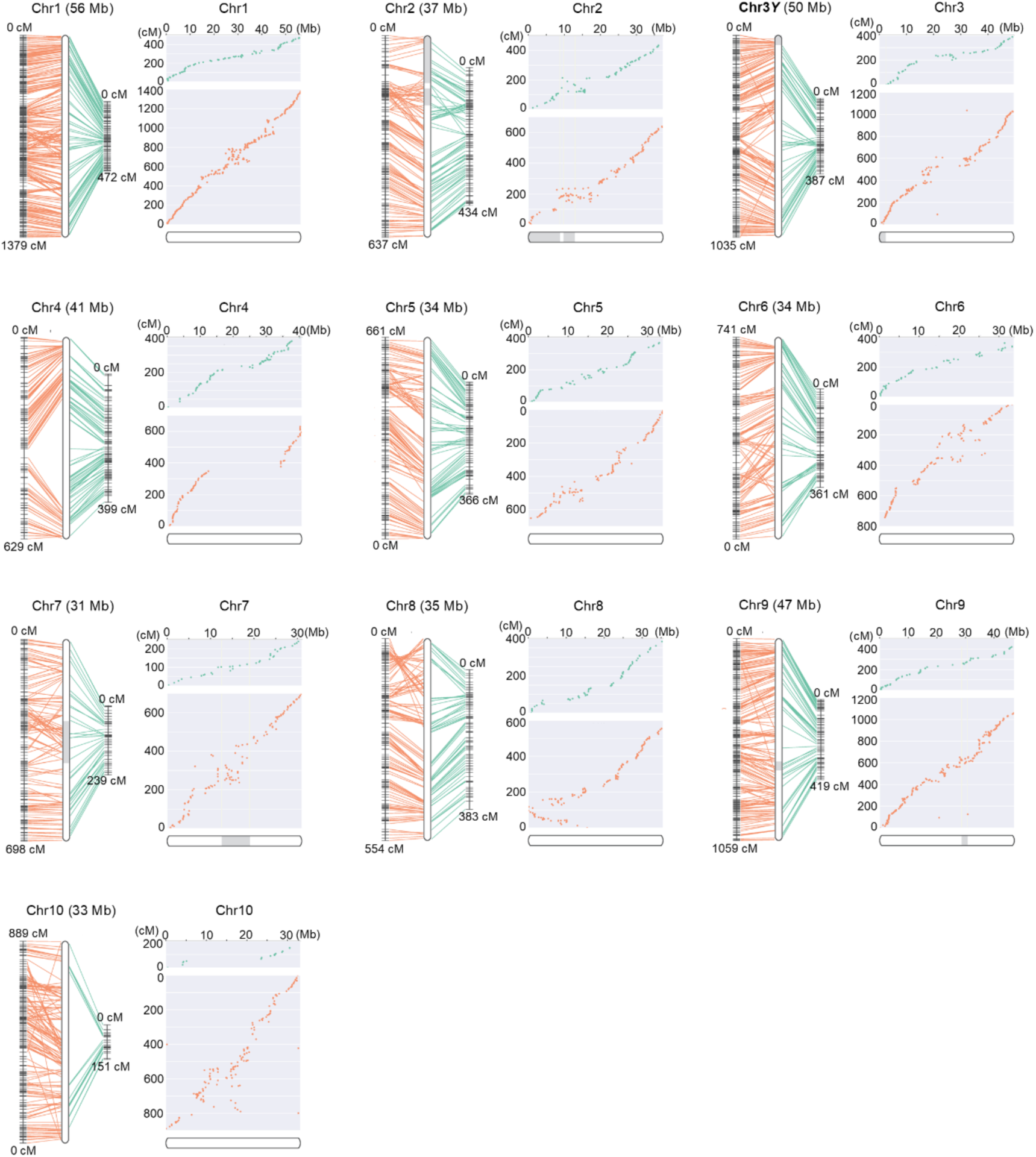
Comparison of genetic and physical maps of chromosomes from the Kita1 (male) haplotype 1 reference genome. For each chromosome, the left panel shows the physical positions of male-parent-heterozygous SNP markers (green) and female-parent-heterozygous SNP markers (orange) and linkage distances between each marker. The right panel shows scatter plots representing the physical positions on the chromosome (Mb; *x*-axis) versus the linkage distances (cM; *y*-axis).

**Fig S15.**
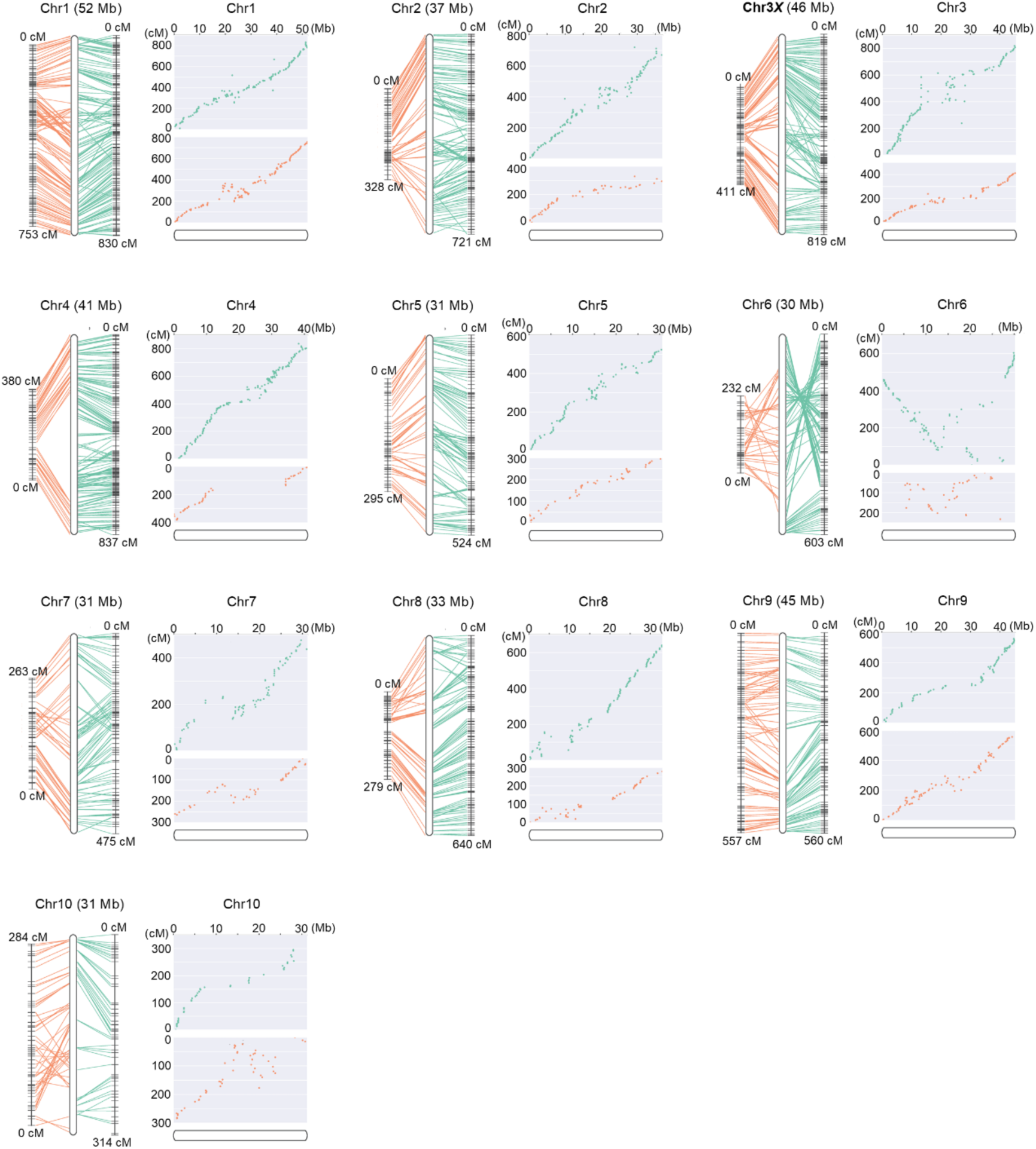
Comparison of genetic and physical maps of chromosomes from the Waka1 (male) reference genome. For each chromosome, the left panel shows the physical positions of male-parent-heterozygous SNP markers (green) and female-parent-heterozygous SNP markers (orange) and linkage distances between each marker. The right panel shows scatter plots representing the physical positions on the chromosome (Mb; *x*-axis) versus the linkage distances (cM; *y*-axis).

**Fig S16.**
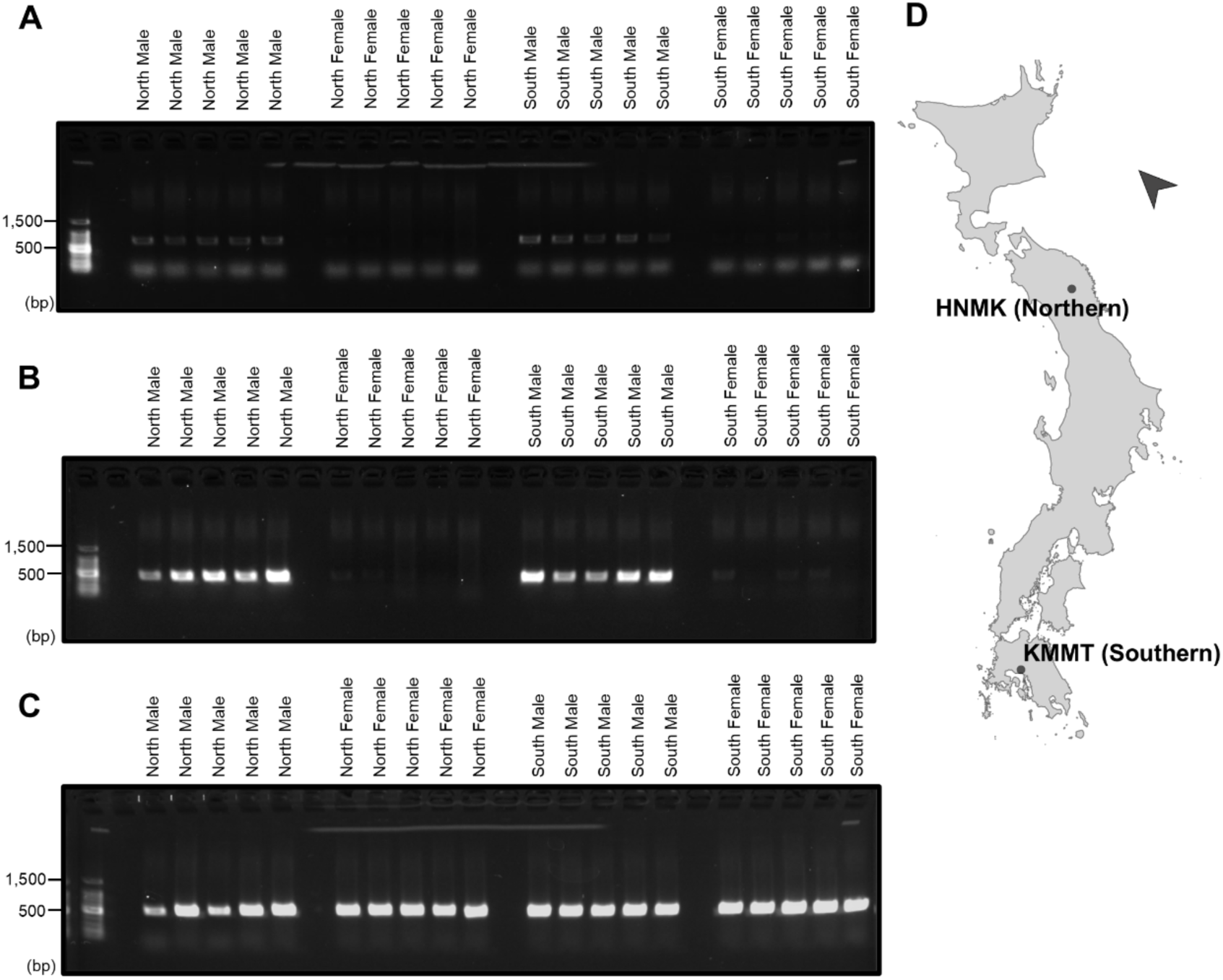
*BLH9* and *HSP90* are male specific in natural populations. **(A)** Amplification of *BLH9* in HNMK (Northern) males, HNMK (Northern) females, KMMT (Southern) males, and KMMT (Southern) females. Each group included five individuals as a replicate. **(B)** Amplification of *HSP90*. **(C)** Amplification of *xanthine dehydrogenase* (*Xdh*) located on the pseudoautosomal regions of chromosome 3 as a control. **(D)** Distribution of the HNMK (Northern) and KMMT (Southern) populations. The map was created with Quantum Geographic Information System (QGIS) software version 3.16.0 (QGIS Development Team 2022). The base map was obtained from the Geospatial Information Authority of Japan, 2020.

**Fig S17.**
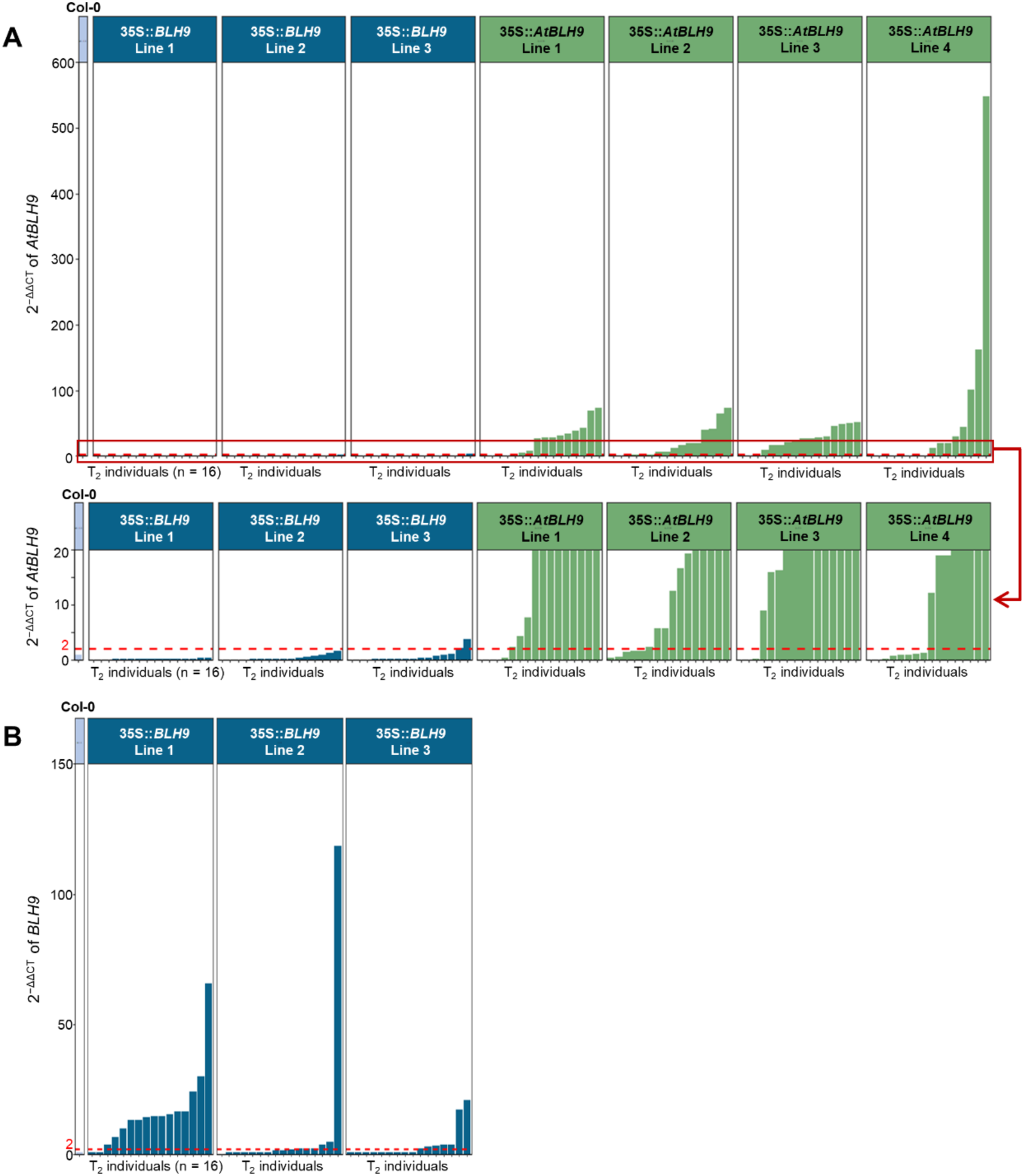
Relative expression levels of *BLH9* and *AtBLH9* in T_2_ individuals harboring 35S::*BLH9* and 35S::*AtBLH9*. Sixteen plants were grown from each of eight lines: Col-0, three T_2_ lines transformed with the construct for *BLH9* from *D. tokoro,* and four T_2_ lines transformed with the construct for *AtBLH9* from *A. thaliana*. The expression levels of *BLH9* and *AtBLH9* in T_2_ plants were confirmed by RT-qPCR. **(A)** 2^−ΔΔCT^ values of *BLH9* in Col-0 and T_2_ individuals harboring 35S::*BLH9* and 35S::*AtBLH9.* Red line indicates 2^−ΔΔCT^ = 2. **(B)** 2^−ΔΔCT^ values of *BLH9* in Col-0 and T_2_ individuals harboring 35S::*BLH9* and 35S::*AtBLH9*.

