## Supplementary Materials and Methods for "Whole-genome sequencing reveals the molecular basis of sex determination in the dioecious wild yam *Dioscorea tokoro*"

**S1 Supplementary materials and methods**

Table of contents

1. Materials

  1.1 Plant materials

  1.2 Whole-genome sequencing of female and male individuals using Oxford Nanopore Technology

  1.3 Illumina library construction and sequencing of female and male individuals

  1.4 RAD-seq for linkage mapping and association analysis

  1.5 RNA library construction and sequencing

  1.6 Small RNA library construction and sequencing

2. Reference assembly

  2.1 Estimation of genome size

  2.2 Quality control

  2.3 *De novo* assembly

  2.4 Polishing and removal of duplicated contigs

  2.5 TE annotation

  2.6 Transcriptome-based gene identification

  2.7 *Ab initio* gene prediction

  2.8 Gene annotation

3. Generation of chromosomes using pseudo-testcross methods

  3.1 Identification of parental-line-specific heterozygous markers

  3.2 Linkage mapping

  3.3 Integration of two parental-specific linkage maps into the chromosome-scale physical map

4. Identification of sex-linked regions by association analysis and coverage analysis

  4.1 Association analysis

  4.2 Coverage analysis

5. Genome structures of the *X* and *Y* chromosomes

  5.1 Gene density and repetitive sequence accumulation in *X*- and *Y*-specific regions

  5.2 Synteny comparison between the *X* and *Y* chromosomes

  5.3 Recombination frequency

6. Identification of candidate genes for sex determination

  6.1 Identification of highly expressed genes in male flowers during early stages of development

  6.2 Identification of candidate genes for sex determination

  6.3 PCR amplification of the candidate genes

7. Identification of candidate miRNAs for sex determination

7.1 Sequence processing
7.2 miRNA prediction and annotation
7.3 Identification of highly expressed miRNAs in male flowers
7.4 Identification of candidate miRNAs for sex determination
8. Overexpression of *BLH9* and *AtBLH9* in *Arabidopsis thaliana*
8.1 Phylogenetic analysis of BLH9 and TALE superfamily proteins
8.2 Plant materials
8.3 Cloning and plasmid construction
8.4 Plant transformation
8.5 RT-qPCR
8.6 Measurement of inflorescence phenotypes

### 1. Materials

#### 1.1 Plant materials

To construct the reference genome, a female *D. tokoro* Waka1 (original code: DT49) plant was collected from Tahara, Wakayama Pref., Japan. A male *D. tokoro*, Kita1 (original code: 110628-5) plant was collected from Waga-Sennin, Kitakami, Iwate Pref., Japan. The 186 $F_1$ progeny, comprising 38 females, 89 males, and 59 non-flowering individuals, were derived from a cross between female parent Waka1 and male parent Kita1. $F_1$ seeds were obtained from the cross in 2011, and 206 $F_1$ individuals were planted in 2012 at Iwate Biotechnology Research Center. The sex phenotypes of the 186 $F_1$ individuals were obtained in 2014 and 2015. To analyze sex-linked regions, female and male individuals were collected from three wild populations: one each in northern Japan (KTKM; Kitakami, Iwate Pref., Japan), central Japan (SHG; Koka, Shiga Pref., Japan), and southern Japan (FKOK; Kasuya, Fukuoka Pref., Japan). To conduct transcriptome analysis, 18 tissue samples were collected for RNA-seq. Male and female flowers were collected from wild populations in Takizawa and Kitakami, Iwate Prefecture, Japan, and non-reproductive organs were collected from Kita1. Fifteen samples were collected for small RNA-seq from the wild population in Koka, Shiga Pref., Japan. To check the male specificity of the candidate genes, five females and five males were collected from two wild populations: one in northern Japan (HNMK; Hanamaki, Iwate Prefecture, Japan) and one in Southern Japan (KMMT; Kumamoto, Kumamoto Prefecture, Japan). The latitudes and longitudes of the sampling sites are listed in Table SM1.

#### 1.2 Whole-genome sequencing of female and male individuals using Oxford Nanopore Technology

Genomic DNA was extracted from fresh leaves of Waka1 (female) and Kita1 (male) plants using NucleoBond HMW DNA (Macherey-Nagel, Düren, Germany) The DNA was subjected to size selection and purification with Short Read Eliminator XL (Circulomics, Baltimore, MD, USA). Libraries for Waka1 (female) long reads were constructed using a Ligation Sequencing Kit SQK-LSK109 (Oxford Nanopore Technologies, Oxford, UK). Libraries for Kita1 (male) long reads were constructed using a Ligation Sequencing Kit V14 SQK-LSK114. The Waka1 (female) library was sequenced on the MinION Mk1C device, and the Kita1 (male) library was sequenced on the PromethION 2 Solo device at Iwate Biological Research Center. The raw sequencing data for Waka1 (female) was subjected to base calling using Guppy v6.1.1 [1] with the option: "-s fastq_guppy422_sup --min_qscore 8 -r". The raw sequencing data for Kita1 (male) were subjected to base calling using dorado v0.8.1 with the dna_r10.4.1_e8.2_400bps_sup@v5.0.0 model and the option: "--min-qscore 8 --emit-fastq -c 10000 -r".

#### 1.3 Illumina library construction and sequencing of female and male individuals

Genomic DNA was extracted from Waka1 (female) and Kita1 (male) using a NucleoSpin Plant II Kit (Macherey-Nagel). Libraries for Waka1 (female) were constructed using a Collibri™ ES DNA Library Prep Kit for Illumina Systems (Invitrogen, Camarillo, CA, USA) and a TruSeq DNA PCR-Free LT Library Prep

Kit (Illumina, San Diego, CA, USA). Sequencing libraries for Kita1 (male) were constructed using a TruSeq DNA PCR-Free LT Library Prep Kit. The libraries were sequenced via MiSeq and HiSeqX. Genomic DNA was extracted from female and male individuals from the northern, central, and southern Japan using a DNeasy Plant Maxi Kit (Qiagen, Hilden, Germany) following the manufacturer's protocol. Contaminating proteins in the extracted DNA lysate were removed by phenol/chloroform extraction. The DNA was purified by ethanol precipitation. Sequencing libraries were constructed using a Collibri™ ES DNA Library Prep Kit for Illumina Systems. The quality and quantity of the sequencing libraries were assessed using a Qubit fluorometer (Invitrogen), an Agilent Bioanalyzer with Agilent High Sensitivity DNA Kit (Agilent Technologies, Waldbronn, Germany), and a qPCR with Library Quantification Kit (Takara Bio, Mountain View, CA, USA). The libraries were sequenced on the HiSeqX platform by Rhelixa, Tokyo, Japan (Table SM2).

##### 1.4 RAD-seq for linkage mapping and association analysis

RAD-seq was performed as previously described [2]. Genomic DNA was extracted from fresh leaves of Waka1, Kita1, and 186 $F_1$ individuals using a NucleoSpin Plant II Kit (Macherey-Nagel). The DNA was digested with the restriction enzymes PacI and NlaIII, and the libraries for 75-bp paired-end reads were sequenced on the Illumina NextSeq 500 platform. Adapters and unpaired reads were removed using FaQCs and PRINSEQ lite. The filtered RAD-seq reads were used to construct linkage maps and for association analysis (Data S1).

##### 1.5 RNA library construction and sequencing

RNA-seq data were obtained from 18 samples, including male and female flowers and non-reproductive organs of *D. tokoro.* The samples included male and female flowers at five stages of development: inflorescence stage 0, inflorescence stage 1, inflorescence stage 2, buds, and flowers. The samples also included eight non-reproductive organs: vegetative shoot apex, leaves, stems, root apex, rhizome root, rhizome stem, rhizome bud, and rhizome storage tissues. Total RNA was extracted from the samples using an RNeasy Plant Mini Kit (Qiagen) as described previously [2] with slight modifications. The cDNA library was constructed using a TruSeq RNA Sample Prep Kit V2 (Illumina) and sequenced on the Illumina NextSeq 500 platform (Table SM3).

##### 1.6 Small RNA library construction and sequencing

Small RNA-seq data were obtained from 15 samples, including male and female flowers and non-reproductive organs of *D. tokoro.* The samples included male and female flowers at five stages of development: inflorescence stage 0, inflorescence stage 1, inflorescence stage 2, buds, and flowers. The samples also included three non-reproductive organs: the vegetative shoot apex, male and female leaves, and male and female stems. Total RNA was extracted from the samples using an Ambion Plant RNA

Isolation Aid (Ambion, Austin, TX, USA) following the manufacturer's protocol with slight modifications. The frozen and powdered samples were mixed with 100 µL of Plant RNA Isolation Aid and 1 mL lysis solution and thoroughly homogenized. The homogenized samples were centrifuged at 15,000 g in a microcentrifuge for 5 min at room temperature. The supernatants with total RNA were transferred to new tubes. The small RNA fraction was isolated from total RNA using a mirVana miRNA Isolation kit (Ambion) following the manufacturer's protocol. The small RNA libraries were constructed using a NEBNext Multiplex Small RNA Library Prep Set for Illumina (New England BioLabs, Ipswich, MA, USA) following the manufacturer's protocol from the adapter ligation step to the PCR amplification step. For quality control, the PCR-amplified cDNA constructs were purified using DNA Clean & Concentrator-5 (Zymo Research, CA, USA) following the manufacturer's protocol. After eluting the purified DNA, size selection using AMPure XP Beads was conducted following the NEBNext Multiplex Small RNA Library Prep Set for Illumina (New England BioLabs) protocol. The quality and quantity of the libraries were assessed using a Qubit fluorometer (Invitrogen), Agilent BioAnalyzer with Agilent High Sensitivity DNA Kit (Agilent Technologies), and qPCR with Library Quantification Kit (Takara Bio). The libraries were sequenced on the NovaSeq 6000 platform at Genebay, Yokohama, Japan (Table SM4).

#### 2. Reference assembly

##### 2.1 Estimation of genome size

The genome size of *D. tokoro* individual Kita1 was estimated by flow cytometry using nuclei prepared from fresh leaf samples. *D. rotundata* accession TDr96-F1 (570 Mb) [2] was used as an internal reference. DNA from isolated nuclei was stained with propidium iodide (PI) and analyzed using a Cell Lab Quanta SC Flow Cytometer (Beckman Coulter, USA) following the manufacturer's protocol. The G1 peak mean value of *D. tokoro* was 206.5, whereas that of *D. rotundata* was 303.6. Based on the ratio between the two species of 0.68 (206.5/303.6), the genome size of *D. tokoro* was estimated to be ~388 Mb (570 Mb × 0.68) (Fig S2).

##### 2.2 Quality control

Whole-genome assembly of female and male individuals was conducted using long reads generated by Oxford Nanopore Technology. The long-read data for Waka1 (female) and Kita1 (male) were obtained as described in section 1.2 above. As the first step in the pipeline for the reference assembly, the raw sequencing data were filtered. For Waka1 (female) reads, the lambda phage genome was removed from the raw reads with NanoLyse v1.2.0 [3]. Reads with an average read quality score of <7 and length of <1,000 bases were removed with Nanofilt v2.8.0 [3]. For Kita1 (male) reads, reads with an average read quality score of <10 and length of <1,000 bases were removed with chopper v0.8.0 [4] (Table SM5).

##### 2.3 *De novo* assembly

The filtered long reads of Waka1 (female) were assembled using Flye v2.9.2 [5] with the options --nanoraw and --scaffold. This assembly step generated 1,880 contigs with N50 of 1,049,091 base pairs and a total size of 438.1 Mb. The filtered long reads of Kita1 (male) were assembled using PECAT [6] with the options genome_size=388000000, prep_min_length=3000, prep_output_coverage=80, corr_output_coverage=80. This assembly step generated 128 contigs with N50 of 33,851,599 base pairs and a total size of 415.0 Mb for haplotype 1 and 415 contigs with N50 of 1,379,312 base pairs and a total size of 300.8 Mb for haplotype 2.

#### 2.4 Polishing and removal of duplicated contigs

The three assembled contig sets were polished (Fig SM1, SM2). To correct the assembled contigs, a consensus module was generated using Racon v1.5.0 [7]. The assembled contigs were polished and corrected using Medaka v1.7.2 (Oxford Nanopore Technologies, 2018) with the option "-m r941_prom_hac_g507" for female contigs and "-m r1041_e82_400bps_sup_g615" for male contigs. Finally, the scaffolds were polished twice using each Illumina short read from Waka1 (female) and Kita1 (male) with Hypo v1.0.3 [8]. For quality control of the Illumina short reads, adapters, reads of <50 bp and low-quality reads with an average quality score < 20 were removed using FaQCs v2.08 [9]. In the Hypo step, the coverage was set to 60 for Waka1 (female) contigs based on a calculation by CoverM v0.6.1 [10]. The coverage was set to 146 for Kita1 (male) haplotype 1 contigs and 181 for Kita1 (male) haplotype 2 contigs. The genome size was set to 443 Mb based on a previous assembly available at the DNA Databank of Japan (DDBJ) database under BioProject PRJDB12945 because the newly constructed male primary scaffolds and female scaffolds were larger than 388 Mb, estimated as described in section 2.1. To evaluate the completeness of the gene set in each step, BUSCO (Bench-Marking Universal Single Copy) v5.2.2 [11] was utilized with "genome" as the assessment mode and Embryophyta odb10 as the database (Table SM6).

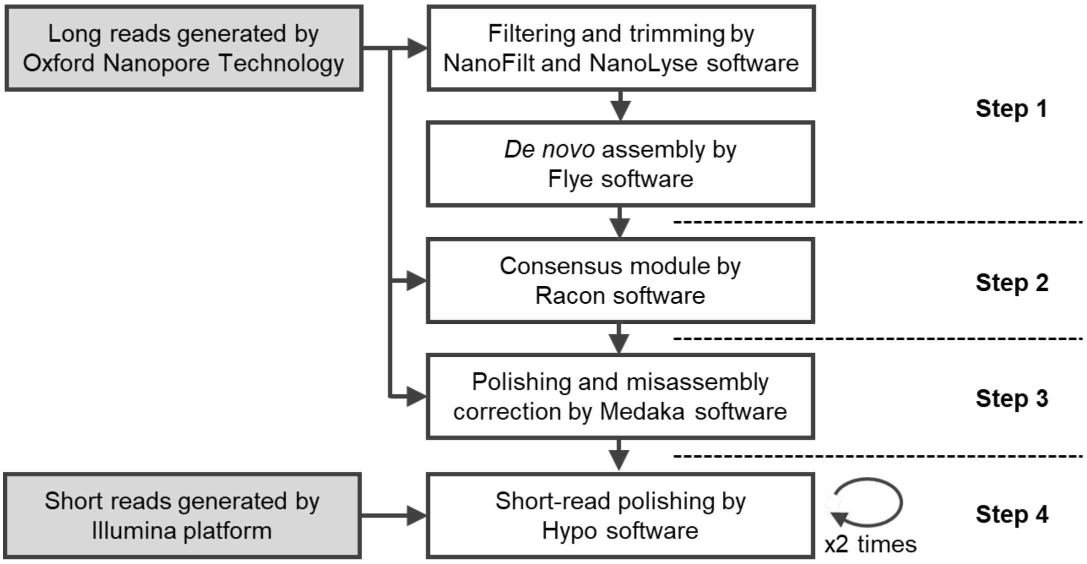


**Fig SM1. Pipeline for genome assembly of Waka1 (female)**

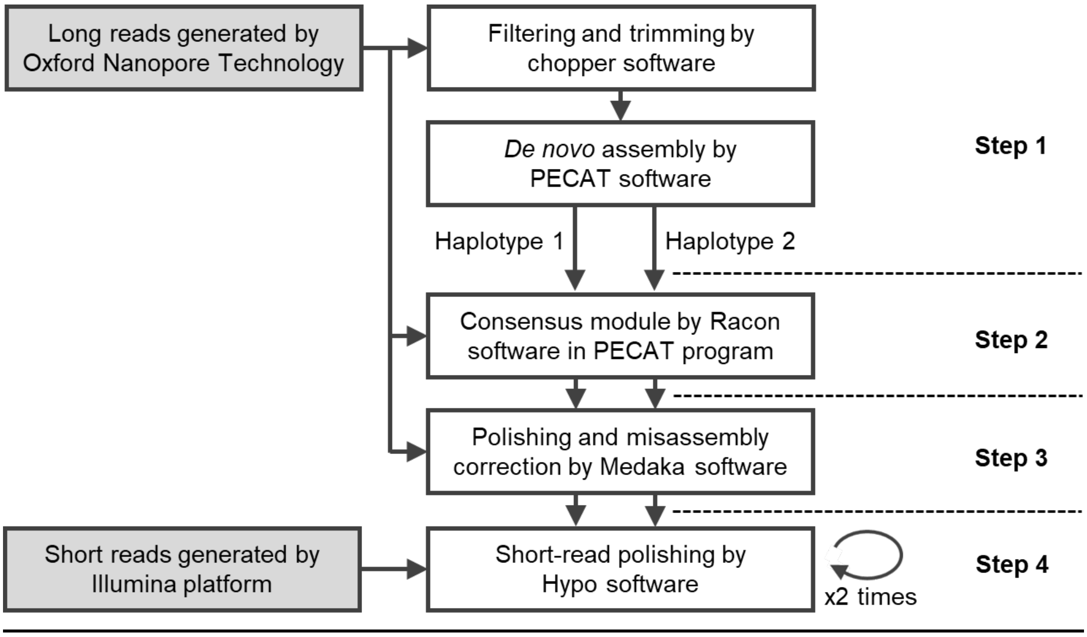


**Fig SM2. Pipeline for genome assembly of Kita1 (male) haplotype 1 and haplotype 2**

#### 2.5 TE annotation

To annotate repetitive sequences, *de novo* repeat libraries were constructed for each assembly using EDTA (The Extensive *de novo* TE Annotator) v2.1.0 [12] with the options --anno 1, --species others, and --step all. Using the resulting files, each genome FASTA file was soft-masked with the perl script make_masked.pl provided in EDTA with the options -maxdiv 30 -minscore 1000 -minlen 1000 -misschar N -hardmask 0.

#### 2.6 Transcriptome-based gene identification

For transcriptome-based gene identification using RNA-seq data from 18 samples of *D. tokoro*, poly(A) sequences and of <50 bp were removed from raw RNA-seq reads with FaQCs v2.10 [9]. Low-quality bases on read ends with an average quality score of <20 were trimmed using PRINSEQ lite v0.20.4 [13] using a window size of 5. Low-quality reads with an average read quality score of <20 were also removed using PRINSEQ lite v0.20.4.

The filtered RNA-seq reads were aligned to the assembled contigs with HISAT2 v2.2.1 [14] with the options "--max-intronlen 15000 --dta". The aligned transcriptomes were assembled using StringTie v3.0.0 [15], and open reading frame (ORF) regions were identified using TransDecoder v5.5.0 [16]. Gene models were excluded from further analysis if their coding sequences (CDS) contained an internal stop codon, if the CDS lengths were not multiples of three, or if they encoded proteins of <50 amino acids.

##### 2.7 *Ab initio* gene prediction

BRAKER2 v2.1.6 [17] was used for *ab initio* gene prediction using the filtered RNA-seq data from *D. tokoro* and protein homology information from *D. alata* and *D. rotundata* with the options --addUTR=on, --softmasking, --etpmode. Protein sequences from *D. alata* TDa95/00328 and *D. rotundata* TDr96_F1 were downloaded from NCBI (accession numbers GCA_020875875.1 and GCF_009730915.1, respectively). The predicted genes were categorized into three groups—gene models fully supported by protein hints, gene models at least partially supported by protein hints, and gene models without any support—using the python script selectSupportedSubsets.py in BRAKER. Gene models fully supported by protein hints were selected, and gene models whose CDS contained an internal stop codon, gene models whose the CDS lengths were not multiples of three, and gene models whose CDS encoded proteins of <50 amino acids were removed.

##### 2.8 Gene annotation

Finally, all predicted genes were merged using GffCompare v0.12.6 [18]. *Ab initio* predicted genes were selected with no or less overlap with genes predicted by StringTie using overlapping class codes: u (no overlap), i (fully contained within a reference intron), and y (contained a reference within its introns). The selected *ab initio* predicted genes were combined with gene annotations generated by gene structure prediction. The final annotations include 37,712 genes in the Waka1 (female) assembly, 36,483 genes in the Kita1 (male) haplotype 1 assembly, and 28,345 genes in the Kita1 (male) haplotype 2 assembly. To evaluate the completeness of the gene sets in the final scaffolds, BUSCO (Bench-Marking Universal Single Copy) v5.3.2 [11] was utilized with "genome" as the assessment mode and Embryophyta odb10 as the database (Table 2).

#### 3. Generation of chromosomes using pseudo-testcross methods

##### 3.1 Identification of parental-line-specific heterozygous markers

Chromosome-scale genome sequences from the assembled contigs were developed using SNP-type heterozygous markers and presence/absence-type heterozygous markers from RAD-seq data from Waka1, Kita1, and 186 $F_1$ individuals. Parental-line-specific heterozygous markers were identified as described previously [19] with the following modifications.

***SNP-type heterozygous markers***

To obtain SNP-type heterozygous markers, the RAD-seq data forWaka1 (female) and Kita1 (male) contigs were aligned using BWA v0.7.18-r1243-dirty [20]. Based on these alignments, SNP-based genotypes were obtained using Bcftools v1.15 [21] with the following commands: (i) mpileup command with the option "-t DP,AD,SP -B -Q 18 -C 50"; (ii) call command with the option "-P 0 -v -m -a GQ,GP"; (iii) filter command with the options "-i 'INFO/MQ≥40, INFO/MQ0F≤0.1, and AVG(GQ)≥10"; and (iv) norm command with the option “-m+any.” Biallelic SNPs were selected with Bcftools view commands with the option “-m 2 -M 2 -v snps”. In addition, variants with low read depth (<10) or low genotype quality scores (<10) in the two parents were removed. Variants with low read depth (<8) or low genotype quality scores (<5) in $F_1$ progenies considered to be missing were also removed, and only variants with low missing rates (<0.3) were retained. After obtaining SNP-based genotypes, heterozygous genotypes with unbalanced allele frequency (out of 0.1-0.9 in $F_1$ progenies) were filtered out. Finally, a binomial test was performed to reject SNPs affected by segregating distortion in the $F_1$ progenies. This binomial test assumes that the probability of success is 0.5 based on the two-side hypothesis, and variants with $p$-values $< 0.001$ were regarded as having segregation distortion. The following SNP-type heterozygous markers were ultimately obtained: 1,000 female-parent-heterozygous SNP markers and 1,983 male-parent-heterozygous SNP markers for the Waka1 (female) reference; 2,303 female-parent-heterozygous SNP markers and 1,194 male-parent-heterozygous SNP markers for the haplotype 1 Kita1 (male) reference; and 1,721 female-parent-heterozygous SNP markers and 1,301 male-parent-heterozygous SNP markers for the haplotype 2 Kita1 (male) reference.

***Presence/absence-type heterozygous markers***

To obtain presence/absence-type heterozygous markers, the RAD-seq data for Waka1 (female) and Kita1 (male) contigs were aligned using BWA v0.7.18-r1243-dirty [20]. Based on the alignment read depth of the two parental plants, Waka1 (female) and Kita1 (male), presence/absence-based genotypes were obtained using Bcftools v1.15 [21] with the following commands: (i) mpileup command with the option “DP,AD,SP,ADF,ADR,INFO/ADF,INFO/ADR -B -Q 18 -C 50”; (ii) call command with the option “-P 0 -A -m -f GQ,GP”; and (iii) view command with the options “-i ‘MAX(FMT/DP)≥4 & MIN(FMT/DP) ≤0’ -g miss -V indels”. These presence/absence-based markers included genotypes in which one parent (Waka1 or Kita1) has sufficient read depth (≥4) and the other parent has no read depth. After obtaining the genotypes in VCF format, continuous positions in VCF format were converted to a feature that provides a region’s start and end coordinate information using the BEDtools v2.31.1 [22] merge command with the option “-d 10 -c 1 -o count”. Only wide features (≥50 bp) were retained in the BED file. The total read base count (i.e., the sum of per-base read depth) was obtained using the Samtools bedcov command. Based on the depth value, presence/absence-based genotypes were determined. For Waka1 (female) and Kita1 (male), genotypes with depth $\geq 4$ were regarded as present genotypes, meaning heterozygosity of presence and absence, and genotypes with depth $= 0$ were regarded as absent genotypes, meaning homozygosity of absence. For $F_1$ progenies, markers with depth $\geq 2$ and depth $= 0$ were classified as present and absent markers, respectively. After obtaining presence/absence-based genotypes, heterozygous genotypes with unbalanced allele frequency (out of 0.1-0.9 in $F_1$ progenies) were filtered out. Finally, a binomial test was performed to reject SNPs affected by segregating distortion in the $F_1$ progenies. This binomial test assumes that the probability of success is 0.5 based on the two-side hypothesis, and variants with $p < 0.001$ were regarded as having segregation distortion. As a result, 3,095 female-parent-heterozygous presence/absence markers and 834 male-parent-heterozygous presence/absence markers were obtained for the Waka1 (female) reference, 1,605 female-parent-heterozygous presence/absence markers and 1,517 male-parent-heterozygous presence/absence markers for the haplotype 1 Kita1 (male) reference, and 1,146 female-parent-heterozygous presence/absence markers and 1,077 male-parent-heterozygous presence/absence markers for the haplotype 2 Kita1 (male) reference.

##### 3.2 Linkage mapping

Linkage maps were constructed based on the SNP-type heterozygous markers and Presence/absence-type heterozygous markers. For each marker set (female-parent-heterozygous marker set and male-parent-heterozygous marker set), the markers were converted into genotype-formatted data for linkage map construction using MSTmap v1.0 [23] with the following parameters: "populationtype DH; distancefunction kosambi; cutoffpvalue 0.000000000001; nomapdist 15.0; nomapsize 0; missingthreshold 25.0; estimationbeforeclustering no; detectbaddata no; objective_function ML". After trimming the orphan linkage groups, the linkage groups were reconstructed and the markers in each linkage group were ordered using R/qtl [24] with the following commands: (i) est.rf command, (ii) switchAlleles command, (iii) formLinkageGroups command with the options "max.rf=0.35, min.lod=6, reorgMarkers=TRUE", (iv) orderMarkers command. Finally, two parental-specific linkage maps were constructed and visualized by Asmap [25] using the mstmap command with the options "bychr = TRUE, anchor = TRUE, dist.fun = 'kosambi'". The linkage maps for the Waka1 (female) assembly and the Kita1 (male) haplotype 1 and 2 assemblies are shown in Fig S4, S5, and S6.

##### 3.3 Integration of two parental-specific linkage maps into the chromosome-scale physical map

Based on two parental-specific linkage maps, the contigs were anchored and linearly ordered as pseudochromosomes using ALLMAPS [26]. Gene-order-based synteny of the three newly constructed reference genomes was detected using MCscan [27] (Tang et al., 2024). The orthologous regions were identified using the "jcvi.compara.catalog ortholog" function with the option "--cscore=.99" and the "jcvi.compara.synteny screen" function with at least 50 collinear gene blocks. The detected collinearity was visualized using the "jcvi.graphics.karyotype" function. The repeat libraries and gene annotations were lifted over from each assembly to the chromosome-scale genome sequence using the liftOver function in ALLMAPS. The telomere repeats were identified using quarTeT v1.2.5 [28] with the option TeloExplorer -c plant. The numbers of anchored contigs and telomere repeats are shown in Table SM7.

#### 4. Identification of sex-linked regions by association analysis and coverage analysis

##### 4.1 Association analysis

To identify sex-linked regions, association analysis was performed using markers obtained by RAD-seq data from the $F_1$ progenies. The RAD-seq data were aligned onto the newly constructed chromosomes from Waka1 (female) and the haplotype 1 and 2 assembly from Kita1 (male) using BWA v0.7.18-r1243-dirty [20]. Based on these alignments, SNP-type markers and presence/absence-type markers were identified as genotype data as described in section 3.1. The presence/absence-type markers were defined based on the aligned read depth of $F_1$ individuals using the cutoffs depth $\geq 3$ as presence and depth = 0 as absence.

The associations between the genotypes and sex phenotypes of the 127 flowering $F_1$ individuals were calculated using Fisher's exact test. The $q$-value for the Fisher's exact test was obtained for each marker by comparing the frequencies of particular alleles and sex phenotypes categorized as female or male. The log transformed $q$-values ($-\log_{10}(q)$) for each position were visualized as Manhattan plots (Fig 2B, C; Fig S10). The false discovery rate (FDR) was set to 0.05 and was corrected by Benjamini-Hochberg correction. Each adjusted threshold is shown in Fig S10. When there were no values for rejected hypotheses before FDR correction in the analysis, threshold = $-\log_{10}(0.05)$ was used. Statistical analysis and visualization were performed using an in-house python script.

The associations between the genotypes and sex phenotypes of the 186 $F_1$ individuals were also checked by composite interval mapping (CIM) using R/qtl v1.70 [24]. The significance of each linkage was tested based on the likelihood-ratio statistic (LOD). The LOD scores were calculated by "cim" with the following options: method="hk", n.marcovar=5, window=10, n.perm=1000. The LOD threshold for linkage was set to a 95% confidence level (alpha = 0.05).

**4.2 Coverage analysis**

To identify regions with different alignment depths between male and female individuals, Illumina short reads from four pairs of male and female were examined: the male parent Kita1 and the female parent Waka1, and three pairs of male and female individuals collected from northern, central, and southern Japan. For quality control of the Illumina short reads, adapters and reads of <50 bp, as well as low-quality reads with an average read quality score <20, were removed with FaQCs v2.10 (Lo & Chain, 2014). The filtered short reads were aligned to each reference genome: Waka1 (female), Kita1 (male) haplotype 1, and Kita1 (male) haplotype 2. Sequence alignment was conducted using BWA v0.7.17-r1188 [20] with the BWA-MEM algorithm setting. The alignment depths were calculated with Samtools v1.16.1 [21]; in this step, reads with mapping quality $\geq 40$ were counted. The alignment depths were normalized by dividing by the mean depth of all positions on the chromosomes. To reduce the effect of error on the depth, a sliding window approach was employed (window size = 150 kb, step size = 10 kb). *X*- and *Y*-specific regions were then identified using the region depths. Regions in which the male depth was 0.5 (half of mean normalized depth) $\pm$ 0.15 and the female depth was 1 (mean normalized depth) $\pm$ 0.15 were selected as *X*-specific regions. Regions in which the male depth was 0.5 (half of mean normalized depth) $\pm$ 0.15 and the female depth was < 0.15 were selected as *Y*-specific regions.

#### 5. Genome structures of the *X* and *Y* chromosomes

##### 5.1 Gene density and repetitive sequence accumulation in *X*- and *Y*-specific regions

Information about gene density and repetitive sequence accumulation was obtained from gene annotations and predicted repetitive sequences of the Waka1 (female) assembly and Kita1 (male) haplotype 1 assembly. Repetitive sequences including Copia LTR retrotransposons, Gypsy LTR retrotransposons, hAT TIR transposons, and Helitrons were predicted using EDTA v2.1.0. Gene density and retrotransposon accumulation were visualized using the "jcvi.graphics.landscape heatmap" function of the jcvi package [27].

##### 5.2 Synteny comparison between the *X* and *Y* chromosomes

Gene-order-based synteny of pseudochromosomes 3*X* and 3*Y* was detected using MCscan [27]. The orthologous regions were identified using the "jcvi.compara.catalog ortholog" function with the option "--cscore=.99" and the "jcvi.compara.synteny screen" function with at least 100 collinear gene blocks. The detected collinearity was visualized using the "jcvi.graphics.karyotype" function.

##### 5.3 Recombination frequency

To evaluate recombination suppression around the SDR in the *D. tokoro* genome, SNP-type heterozygous markers were obtained from RAD-seq data from Waka1, Kita1, and 186 $F_1$ individuals. The RAD-seq data were aligned onto Waka1 (female)contigs and Kita1 (male) haplotype 1 contigs with BWA v0.7.18-r1243-dirty [20]. Based on these alignments, SNP-based genotypes were obtained as described in section 3.1. After SNP-based genotypes were obtained, SNP markers showing 1:1 segregation of homozygous and heterozygous genotypes in the $F_1$ progenies were selected. To ensure the segregation patterns, heterozygous genotypes with unbalanced allele frequency (out of 0.4-0.6 in $F_1$ progenies) were filtered out to select SNP markers. Finally, a binomial test was performed to reject SNPs affected by segregating distortion in the $F_1$ progenies. This binomial test assumes that the probability of success is 0.5 based on the two-side hypothesis; variants with $p$-value < 0.001 were regarded as having segregation distortion. After marker selection, linkage distances were obtained based on genetic linkage maps constructed using MSTmap v1.0 [92] with the following parameters: "populationtype DH; distancefunction kosambi; cutoffpvalue 0.000000000001; nomapdist 15.0; nomapsize 0; missingthreshold 25.0; estimationbeforeclustering no; detectbaddata no; objective_function ML". After trimming the orphan linkage groups, linkage groups were reconstructed and markers in each linkage group were ordered using R/qtl [24] with the following commands: (i) est.rf command, (ii) switchAlleles command, (iii) formLinkageGroups command with the options "max.rf=0.35, min.lod=6, reorgMarkers=TRUE". Finally, the markers were reordered by Asmap

[25] using the mstmap command with the options "bychr = FALSE, anchor = FALSE, dist.fun = 'kosambi'". The linkage distances and physical distances of Kita1 (male) haplotype 1 chromosomes were compared and visualized using ALLMAPS [26].

#### 6. Identification of candidate genes for sex determination

##### 6.1 Identification of highly expressed genes in male flowers during early stages of development

Differential expression analysis was performed using the filtered RNA-seq data from male flowers, female flowers, and non-reproductive organs. The trimmed RNA-seq reads were aligned to the Kita1 (male) haplotype 1 assembly using HISAT v2.2.1 [14]. The mapped reads were counted with the featureCounts function in Subread v2.0.1 [29]. The minimum fragment length was set to 19, and the attribute type was set to transcript id. Differential expression analysis was performed using DESeq2 v3.15 [30] in R v4.1.1 for two comparisons: male vs. female flowers at three early stages of development (inflorescence stages 0, 1, and 2); and male flowers at three stages of early development vs. non-reproductive organs. The false discovery rate threshold was set to 0.05.

##### 6.2 Identification of candidate genes for sex determination

As the first step in candidate gene identification, 51 genes located on *Y*-specific regions were identified (Table 3). In this step, *Y*-specific regions were detected with liberal thresholds in sliding windows (window size = 50 kb, step size = 1 kb). In the second step, among the *Y*-specific genes, 26 genes that were highly expressed during three early stages of male vs. female flower development were selected based on differential expression analysis using negative binomial generalized linear models ($p < 0.05$). Ten genes were also identified that were significantly upregulated in male vs. female flowers during three stages of early development based on the $\log_2$ ratio of the mean of normalized counts in each group ($\log_2 FC > 2$). In the third step, among the *Y*-specific genes, 10 genes were identified that were highly expressed in male flowers during three stages of early development compared to non-reproductive organs ($p < 0.05$). Three genes that were significantly upregulated in male flowers during three stages of early development compared to non-reproductive organs were also identified ($\log_2 FC > 2$). Finally, two *Y*-specific genes were selected with significantly upregulated expression in male flowers during three stages of early development in both transcriptome comparisons ($q < 0.05$ and $\log_2 FC > 2$). The similarity of the candidate gene products was compared to known proteins using BLASTX with the Swiss-Prot function.

##### 6.3 PCR amplification of the candidate genes

The male specificity of the two candidate genes was also confirmed by PCR amplification using five females and five males from two wild populations, one in northern Japan and one in southern Japan. Genomic DNA was extracted from the samples using a Maxwell RSC Plant DNA Kit (Promega, Madison, WI, USA). DNA fragments of *BLH9* and *HSP90* were amplified with Quick Taq HS DyeMix (TOYOBO,

Osaka, Japan) using primers for the first exons of *BLH9* and *HSP90*. The *xanthine dehydrogenase* (*Xdh*) gene located on the pseudoautosomal region of chromosome 3 was also amplified as a control. The *Xdh* gene has been used for nuclear phylogenetic analysis of the genus *Dioscorea* [31], and we obtained the *Xdh* sequence of *D. tokoro* based on the *Xdh* sequence of the related species *D. caucasica* (GenBank: KY712749.1) using BLASTX. All primers used in this step are listed in Table SM8.

#### 7. Identification of candidate miRNAs for sex determination

##### 7.1 Sequence processing

For quality control of the small RNA-seq data, adapters and low-quality reads with an average read quality score > 20 were removed with FaQCs v2.10 [9]. Reads shorter than 19 bp and longer than 25 bp were removed using Seqkit v2.3.0 [32]. To predict miRNAs using miRDeep-P2 v1.1.4 [33], the filtered reads were preprocessed into the designated format. The filtered reads were parsed into FASTA format with Seqkit v2.3.0, and redundant sequences were removed.

##### 7.2 miRNA prediction and annotation

miRNAs were predicted as described previously [34] with several modifications. Novel miRNAs were detected from preprocessed reads using the miRDP2-v1.1.4_pipeline.bash script of miRDeep-P2 v1.1.4. The predicted miRNA sequences and their corresponding precursor information were extracted using the perl script parse_miRDP2_prediction.pl [34]. The miRNA sequences were annotated to the Kita1 (male) haplotype 1 assembly. First, a BLAST database was created from the *D. tokoro* reference genome using the makeblastdb function of Blast+ v2.2.31 [35]. miRNAs hits in the *D. tokoro* BLAST database were selected using the blastn function of BLAST+ v2.2.31 with output format 6. To eliminate missing candidate miRNAs, the word size was set to 7 and the *e*-value threshold was set to 1,000. The output tab file was transferred to gff3 format using the blast2gff.py script of genomeGTFtools v1.3 [36].

##### 7.3 Identification of highly expressed miRNAs in male flowers

The trimmed small RNA-seq reads were aligned to the predicted miRNA references with Bowtie v1.3.1 [37]. The mapped reads were counted using the perl script bam2ref_counts.pl, and the read counts data for each sample were combined using the perl script combine_htseq_counts.pl [34]. Differential expression analysis was performed using DESeq2 v3.15 [30] in R v4.1.1 for two comparisons: male vs. female flowers at three stages of early development (inflorescence stages 0, 1, and 2); and male flowers at three stages of early development vs. non-reproductive organs. The false discovery rate threshold was set to 0.05.

##### 7.4 Identification of candidate miRNAs for sex determination

As the first step in candidate miRNA identification, 15 miRNAs were located on *Y*-specific regions (Table 4). In this step, *Y*-specific regions were detected with liberal thresholds in sliding windows (window size =

50 kb, step size = 1 kb). Among the *Y*-specific miRNAs, no miRNAs were highly expressed in male vs. female flowers during early stages of development ($p < 0.05$).

#### 8. Overexpression of *BLH9* and *AtBLH9* in *Arabidopsis thaliana*

##### 8.1 Phylogenetic analysis of BLH9 and TALE superfamily proteins

Phylogenetic analysis of *BLH9* and its homologs was performed using the sequences of BLH9 and TALE superfamily proteins from *Arabidopsis thaliana.* The sequences of *A. thaliana* TALE superfamily proteins were obtained from The Arabidopsis Information Resource (TAIR) (Table SM9) based on Hamant & Pautot 2010. The sequences were aligned with Molecular Evolutionary Genetics Analysis (MEGA) v11.0.13 [38] using the ClustalW algorithm. Based on the alignment, a phylogenic tree was constructed with the maximum likelihood method in MEGA using the Jones-Taylor-Thornton model. The bootstrap values were calculated using the nearest neighbor interchange technique with 1,000 replications. The tree was visualized using Interactive Tree Of Life (iTOL) v7.0 [39].

##### 8.2 Plant materials

All *Arabidopsis thaliana* lines are in the Col-0 accession background. Surface-sterilized *A. thaliana* seeds were incubated at 4°C in the dark for two to five days and grown on Murashige and Skoog (MS) medium (1/2 MS, 0.5% sucrose, B5 vitamin solution, 2 mM MES, pH 5.7) under controlled conditions (23°C, 10-h photoperiod). After germination, the plants were transferred to soil and grown under controlled conditions (23°C, 16-h photoperiod, 57 μmol $m^{-2}$ $s^{-1}$ of light).

##### 8.3 Cloning and plasmid construction

Two full-length cDNA clones were used for In-Fusion cloning: *BLH9* from inflorescence stage 2 of the *D. tokoro* Kita1 male individual and *AtBLH9* from the first internode of *A. thaliana* Col-0. Total RNA was extracted from the samples using a Maxwell RSC Plant RNA Kit (Promega) and was reverse-transcribed into cDNA using a ReverTra Ace qPCR RT Kit FSQ-101 (TOYOBO). *BLH9* fragment was amplified from the *D. tokoro* cDNA using KOD FX Neo (TOYOBO). *AtBLH9* fragment was amplified from *A. thaliana* cDNA using PrimeSTAR GXL DNA Polymerase (Takara Bio). After purification using NucleoSpin Gel and PCR Clean-up (Macherey-Nagel), each cDNA fragment was cloned into EcoRI and BamHI sites of the pBICP35 binary vector, which contains the CaMV35S promoter [40], using an In-Fusion HD Cloning Kit (Takara Bio). The insertion was confirmed using Quick Taq HS DyeMix (TOYOBO), and the inserted sequences were confirmed by DNA sequencing (Eurofin Genomics, Tokyo, Japan). All primers used in this step are listed in Table SM10.

##### 8.4 Plant transformation

The pBICP35::BLH9 and pBICP35::AtBLH9 were transformed into *Agrobacterium tumefaciens* strain

GV3101::pMP90 by electroporation. *A. thaliana* transformation was performed by the floral dip method with *A. tumefaciens* strains carrying the binary expression plasmids, and kanamycin-resistant $T_1$ plants were selected. $T_2$ lines that showed a 3:1 segregation ratio for kanamycin resistance were selected, as they were thought to contain a single T-DNA. The selected $T_2$ lines and Col-0 were used to examine inflorescence phenotypes.

##### 8.5 RT-qPCR

To examine inflorescence phenotypes, 16 plants each were grown from eight lines: Col-0, three $T_2$ lines transformed with the plasmid bearing the *BLH9* construct, and four $T_2$ lines transformed with the plasmid bearing the *AtBLH9* construct. To eliminate the effects of kanamycin selection on plant growth, all seeds of $T_2$ plants and Col-0 were grown in Murashige and Skoog (MS) medium without kanamycin. The expression levels of *BLH9* and *AtBLH9* in $T_2$ plants were confirmed by RT-qPCR (Data S4). Total RNA was extracted from each plant using a Maxwell RSC Plant RNA Kit (Promega), and cDNAs were synthesized using a ReverTra Ace qPCR RT Kit FSQ-101 and ReverTra Ace qPCR RT Master Mix FSQ-201S (TOYOBO). RT-qPCR was conducted using PowerUp SYBR Green Master Mix and a StepOne Real-Time PCR System with the following settings: Quantitation Comparative $C_T$ ($\Delta\Delta C_T$) and Standard cycling mode with StepOne Software v2.3 (Applied Biosystems, Waltham, MA, USA). The relative expression levels of the inserted genes were calculated by the $2^{-\Delta\Delta CT}$ method, and $2^{-\Delta\Delta CT}$ values were < 2 without overexpression (Fig S17). $T_2$ plants were separated into four groups: *BLH9* Control ($2^{-\Delta\Delta CT} \leq 2$), *BLH9 OX* ($2^{-\Delta\Delta CT} > 2$), *AtBLH9* Control ($2^{-\Delta\Delta CT} \leq 2$), and *AtBLH9 OX* ($2^{-\Delta\Delta CT} > 2$). *Ubiquitin C* (*UBC*) was used as a reference gene. The primers used in RT-qPCR are listed in Table SM10.

##### 8.6 Measurement of inflorescence phenotypes

At 30 days after seedlings were transplanted to soil, three phenotypes were measured: inflorescence height, mean fruit length, and internode length. Inflorescence height represents the length of the main inflorescence stem from the rosette to apex. The mean fruit length was obtained from ten fruits of the main inflorescence at positions 2 to 11 counting from the lowest fruit on the main stem. When there were fewer than ten fruits, all fruits at positions 2 to 11 were examined to obtain mean fruit length. Aborted buds were skipped. Internode length represents the stem length between two fruits. For each plant, ten internodes were obtained from the main inflorescence at positions 2 to 11 counted from the lowest fruit on the main stem. When there were fewer than ten internodes, all internodes at positions 2 to 11 were included to obtain internode length data. All phenotypic data are listed in Data S5, S6, and S7. The mean inflorescence heights and mean fruit lengths were compared by Wilcoxon rank sum test between the five groups: Col-0, *BLH9* Control, *BLH9 OX*, *AtBLH9* Control, and *AtBLH9 OX*. The variations in internode lengths were compared by performing an *F*-test between the five groups. The *p*-values were adjusted by Bonferroni correction.

**Table SM1. Sampling sites of *Dioscorea tokoro*.**

| Sample | Sex | Site | Latitude | Longitude |
|---|---|---|---|---|
| **1.2 Whole-genome sequencing of female and male individuals using Oxford Nanopore Technology** | | | | |
| Waka1 | Female | Tahara, Wakayama Pref., Japan | 33°32'16.8"N | 135°51'36.0"E |
| Kita1 | Male | Kitakami, Iwate Pref., Japan | 39°17'42.0"N | 140°53'45.6"E |
| **1.3 Illumina library construction and sequencing of male and female individuals** | | | | |
| Waka1 | Female | Tahara, Wakayama Pref., Japan | 33°32'16.8"N | 135°51'36.0"E |
| Kita1 | Male | Kitakami, Iwate Pref., Japan | 39°17'42.0"N | 140°53'45.6"E |
| Female from KTKM (northern area) | Female | Kitakami, Iwate Pref., Japan | 39°18'25.0"N | 140°54'07.0"E |
| Male from KTKM (northern area) | Male | Kitakami, Iwate Pref., Japan | 39°18'25.0"N | 140°54'07.0"E |
| Female from SHG (central area) | Female | Koka, Shiga Pref., Japan | 34°56'24.1"N | 136°13'05.3"E |
| Male from SHG (central area) | Male | Koka, Shiga Pref., Japan | 34°56'24.1"N | 136°13'05.3"E |
| Female from FKOK (southern area) | Female | Kasuya, Fukuoka Pref., Japan | 33°38'08.7"N | 130°30'38.2"E |
| Male from FKOK (southern area) | Male | Kasuya, Fukuoka Pref., Japan | 33°38'08.7"N | 130°30'38.2"E |
| **1.5 RNA libraries and sequencing** | | | | |
| Kita1 | Male | Kitakami, Iwate Pref., Japan | 39°17'42.0"N | 140°53'45.6"E |
| Female from KTKM (northern area) | Female | Takizawa and Kitakami, Iwate Pref., Japan | | |
| Male from KTKM (northern area) | Male | Takizawa and Kitakami, Iwate Pref., Japan | | |
| **1.6 Small RNA libraries and sequencing** | | | | |
| Female from SHG (central area) | Female | Koka, Shiga Pref., Japan | 34°56'00.8"N | 136°13'30.5"E |
| Male from SHG (central area) | Male | Koka, Shiga Pref., Japan | 34°56'00.8"N | 136°13'30.5"E |
| **6.3 PCR amplification of the candidate genes** | | | | |
| Female from HNMK (northern area) | Female | Hanamaki, Iwate Pref., Japan | 39°22'10.0"N | 141°09'16.0"E |

| | | | | |
|---|---|---|---|---|
| Male from HNMK (northern area) | Male | Hanamki, Iwate Pref., Japan | 39°22'10.0"N | 141°09'16.0"E |
| Female from KMMT (southern area) | Female | Kumamoto, Kumamoto Pref., Japan | 32°53'34.8"N | 130°39'22.7"E |
| Male from KMMT (southern area) | Male | Kumamoto, Kumamoto Pref., Japan | 32°53'34.8"N | 130°39'22.7"E |

**Table SM2. Summary of Illumina short reads from the whole genome.**

| Sample | Original fastq | | Filtered fastq | | Genome coverage | Sequencing platform | Accession no. |
|---|---|---|---|---|---|---|---|
| | Number of reads | Total base pairs (Gbp) | Number of reads | Total base pairs (Gbp) | | | |
| Waka1 (female) | 185,338,998 | 27.9 | 180,836,694 | 27.1 | 69.9× | HiSeqX | SRR32328377<br>SRR32328330 |
| Kita1 (male) | 350,412,788 | 62.4 | 347,141,258 | 61.6 | 159.0× | MiSeq, HiseqX | DRX333479<br>DRX335960 |
| Female (northern area) | 41,867,422 | 6.28 | 41,769,731 | 5.95 | 15.3× | HiSeqX | SRR32328338 |
| Male (northern area) | 54,337,944 | 8.15 | 54,187,471 | 7.81 | 20.1× | HiSeqX | SRR32328337 |
| Female (central area) | 64,740,690 | 9.71 | 64,587,572 | 8.95 | 23.1× | HiSeqX | SRR32328334 |
| Male (central area) | 71,080,268 | 10.7 | 70,966,105 | 9.73 | 25.1× | HiSeqX | SRR32328333 |
| Female (southern area) | 72,491,790 | 10.9 | 72,369,383 | 9.76 | 25.2× | HiSeqX | SRR32328336 |
| Male (southern area) | 47,610,492 | 7.14 | 47,496,663 | 6.72 | 17.3× | HiSeqX | SRR32328335 |

Genome coverage was estimated based on the genome size of *D. tokoro* (388 Mb).

**Table SM3. RNA-seq data generated from libraries constructed from different tissues of *Dioscorea tokoro*.**

| Sample | Sex | Original fastq | | Filtered fastq | | Sequencing platform | Accession no. |
|---|---|---|---|---|---|---|---|
| | | Number of reads | Total base pairs (Gbp) | Number of reads | Total base pairs (Gbp) | | |
| Inflorescence stage 0 | Male | 24,276,259 | 1.81 | 24141016 | 1.79 | NextSeq500 | SRR32328332 |
| Inflorescence stage 1 | Male | 24,765,787 | 1.84 | 24637348 | 1.83 | NextSeq500 | SRR32328331 |
| Inflorescence stage 2 | Male | 27,102,871 | 2.02 | 26962941 | 2.00 | NextSeq500 | SRR32328328 |
| Bud | Male | 26,096,017 | 1.94 | 25945931 | 1.93 | NextSeq500 | SRR32328327 |
| Flower | Male | 24,042,221 | 1.79 | 23890712 | 1.77 | NextSeq500 | SRR32328326 |
| Inflorescence stage 0 | Female | 26,613,535 | 1.98 | 26475567 | 1.97 | NextSeq500 | SRR32328325 |
| Inflorescence stage 1 | Female | 24,109,359 | 1.80 | 23993280 | 1.78 | NextSeq500 | SRR32328324 |
| Inflorescence stage 2 | Female | 25,689,180 | 1.91 | 25564897 | 1.90 | NextSeq500 | SRR32328323 |
| Bud | Female | 26,346,910 | 1.96 | 26211849 | 1.95 | NextSeq500 | SRR32328322 |
| Flower | Female | 24,003,719 | 1.79 | 23896870 | 1.78 | NextSeq500 | SRR32328321 |
| Vegetative shoot apex | Kita1 (male) | 23,051,829 | 1.72 | 22935643 | 1.70 | NextSeq500 | SRR32328320 |
| Leaf | Kita1 (male) | 24,299,715 | 1.81 | 24144324 | 1.79 | NextSeq500 | SRR32328319 |
| Stem | Kita1 (male) | 19,216,778 | 1.43 | 19079127 | 1.42 | NextSeq500 | SRR32328317 |
| Root apex | Kita1 (male) | 23,592,409 | 1.76 | 23473872 | 1.74 | NextSeq500 | SRR32328316 |
| Rhizome bud | Kita1 (male) | 26,479,350 | 1.97 | 26346195 | 1.96 | NextSeq500 | SRR32328315 |
| Rhizome root | Kita1 (male) | 22,952,532 | 1.71 | 22819053 | 1.70 | NextSeq500 | SRR32328314 |
| Rhizome stem | Kita1 (male) | 25,506,326 | 1.90 | 25375473 | 1.89 | NextSeq500 | SRR32328313 |
| Rhizome storage | Kita1 (male) | 25,238,872 | 1.88 | 25120091 | 1.87 | NextSeq500 | SRR32328312 |

**Table SM4. Small RNA-seq data generated from libraries constructed from different tissues of *Dioscorea tokoro*.**

| Sample | Sex | Original fastq | | Filtered fastq | | Sequencing platform | Accession no. |
|---|---|---|---|---|---|---|---|
| | | Number of reads | Total base pairs (Gbp) | Number of reads | Total base pairs (Gbp) | | |
| Inflorescence stage 0 | Male | 35,195,175 | 1.35 | 35,092,102 | 1.34 | NovaSeq6000 | SRR32328179 |
| Inflorescence stage 1 | Male | 50,547,350 | 1.79 | 50,377,342 | 1.78 | NovaSeq6000 | SRR32328178 |
| Inflorescence stage 2 | Male | 17,635,303 | 0.64 | 17,575,869 | 0.64 | NovaSeq6000 | SRR32328177 |
| Bud | Male | 48,611,510 | 1.71 | 48,424,940 | 1.7 | NovaSeq6000 | SRR32328176 |
| Flower | Male | 40,153,242 | 1.4 | 39,999,479 | 1.39 | NovaSeq6000 | SRR32328174 |
| Leaf | Male | 52,426,728 | 1.89 | 52,155,143 | 1.88 | NovaSeq6000 | SRR32328173 |
| Stem | Male | 32,635,855 | 1.2 | 32,507,263 | 1.19 | NovaSeq6000 | SRR32328172 |
| Inflorescence stage 0 | Female | 5,404,699 | 0.21 | 5,390,903 | 0.21 | NovaSeq6000 | SRR32328171 |
| Inflorescence stage 1 | Female | 42,777,693 | 1.57 | 42,648,118 | 1.57 | NovaSeq6000 | SRR32328170 |
| Inflorescence stage 2 | Female | 44,021,412 | 1.62 | 43,891,162 | 1.61 | NovaSeq6000 | SRR32328169 |
| Bud | Female | 29,790,989 | 1.05 | 29,708,920 | 1.05 | NovaSeq6000 | SRR32328168 |
| Flower | Female | 20,022,247 | 0.69 | 19,951,825 | 0.69 | NovaSeq6000 | SRR32328167 |
| Leaf | Female | 67,975,326 | 2.4 | 67,650,255 | 2.38 | NovaSeq6000 | SRR32328166 |
| Stem | Female | 40,637,806 | 1.52 | 40,457,258 | 1.51 | NovaSeq6000 | SRR32328165 |
| Vegetative shoot apex | - | 34,742,713 | 1.3 | 34,638,091 | 1.3 | NovaSeq6000 | SRR32328163 |

**Table SM5. Summary of filtered Oxford Nanopore Technology reads of female *Dioscorea tokoro*.**

| Feature | Waka1 (female) | Kita1 (Male) |
|---|---|---|
| Number of reads | 668,621 | 3,890,192 |
| Total base pairs (Gbp) | 15.9 | 42.6 |
| Genome coverage | 41.1× | 109.8× |
| Average fragment size (bp) | 23,825.7 | 10,943 |
| Longest fragment (bp) | 210,758 | 540,057 |
| Shortest fragment (bp) | 1,000 | 1,000 |
| Fragment N50 (bp) | 35,476 | 27,222 |
| Accession number | SRR32328378 | SRR32328156 |

Genome coverage was estimated based on the genome size of *D. tokoro* (388 Mb).

**Table SM6. Assembly summary of Oxford Nanopore Technology (ONT) reads of male and female *Dioscorea tokoro*.**

| | Contigs | Total number of contigs | Total base pairs (bp) | Contig size (bp) | | | | Complete BUSCOs (%) |
|---|---|---|---|---|---|---|---|---|
| | | | | Average | Longest | Shortest | N50 | |
| Step 1 | Waka1 (female) | 1,880 | 438,103,959 | 233,034.0 | 22,942,145 | 592 | 1,049,091 | 97.8 |
| | Kita1 (male) haplotype1 | 128 | 415,009,815 | 3,242,264.2 | 56,343,901 | 559 | 33,851,599 | 98.4 |
| | Kita1 (male) haplotype2 | 415 | 300,822,613 | 724,873.8 | 5,182,372 | 17,347 | 1,379,312 | 76.6 |
| Step 2 | Waka1 (female) | 1,780 | 437,187,904 | 245,611.2 | 22,904,641 | 600 | 1,051,903 | 97.7 |
| | Kita1 (male) haplotype1 | 120 | 414,981,487 | 3,458,179.1 | 56,355,674 | 5,090 | 33,873,162 | 98.4 |
| | Kita1 (male) haplotype2 | 414 | 300,776,275 | 726,512.7 | 5,183,242 | 17,360 | 1,380,047 | 76.6 |
| Step 3 | Waka1 (female) | 1,821 | 438,194,476 | 240,634.0 | 22,941,794 | 600 | 982,230 | 97.6 |
| | Kita1 (male) haplotype1 | 172 | 414,788,701 | 2,411,562.2 | 56,391,428 | 1,051 | 33,885,780 | 98.3 |
| | Kita1 (male) haplotype2 | 414 | 300,776,275 | 726,512.7 | 5,183,242 | 17,360 | 1,380,047 | 76.7 |
| Step 4 | Waka1 (female) | 1,821 | 438,139,280 | 240,603.7 | 22,951,840 | 600 | 982,016 | 98.2 |
| First hypo | Kita1 (male) haplotype1 | 172 | 414,446,737 | 2,409,574.1 | 56,339,659 | 1,051 | 33,852,229 | 98.4 |
| | Kita1 (male) haplotype2 | 440 | 300,437,562 | 682,812.6 | 5,181,085 | 868 | 1,379,306 | 76.7 |
| Step 4 | Waka1 (female) | 1,821 | 438,105,977 | 240,585.4 | 22,951,365 | 600 | 981,934 | 97.6 |
| Second hypo | Kita1 (male) haplotype1 | 172 | 414,436,429 | 2,409,514.10 | 56,338,240 | 1,051 | 33,850,955 | 98.0 |
| | Kita1 (male) haplotype2 | 440 | 300,420,452 | 682,773.80 | 5,180,395 | 868 | 1,379,077 | 76.0 |

**Table SM7. Number of anchored contigs and telomere repeats.**

| Reference | Chromosome | Length (bp) | Number of anchored contigs | Telomere | | |
|---|---|---|---|---|---|---|
| | | | | Side(s) containing telomeres | Number of repeats on left side | Number of repeats on right side |
| Waka1 (female) | Chr1 | 52,115,987 | 104 | Neither | 0 | 0 |
| | Chr2 | 37,046,401 | 79 | Neither | 0 | 0 |
| | Chr3 | 45,688,850 | 86 | Neither | 0 | 0 |
| | Chr4 | 41,014,181 | 37 | Neither | 0 | 0 |
| | Chr5 | 30,704,362 | 77 | Neither | 0 | 0 |
| | Chr6 | 30,230,714 | 80 | Neither | 0 | 0 |
| | Chr7 | 31,220,134 | 76 | Neither | 0 | 0 |
| | Chr8 | 33,058,290 | 39 | Neither | 0 | 0 |
| | Chr9 | 45,171,277 | 77 | Neither | 0 | 0 |
| | Chr10 | 31,241,263 | 75 | Neither | 0 | 0 |
| Kita1 (male) haplotype 1 | Chr1 | 56,338,240 | 1 | Both | 536 | 945 |
| | Chr2 | 37,381,473 | 4 | Right | 0 | 737 |
| | Chr3 | 49,521,298 | 2 | Both | 563 | 573 |
| | Chr4 | 40,747,576 | 1 | Both | 692 | 859 |
| | Chr5 | 33,850,955 | 1 | Both | 921 | 858 |
| | Chr6 | 33,716,633 | 1 | Both | 733 | 1,256 |
| | Chr7 | 30,996,712 | 3 | Both | 591 | 721 |
| | Chr8 | 35,498,219 | 1 | Both | 1,199 | 749 |

| | | | | | | |
|---|---|---|---|---|---|---|
| | Chr9 | 47,071,449 | 3 | Both | 750 | 668 |
| | Chr10 | 33,223,422 | 1 | Both | 863 | 699 |
| Kita1 (male) | Chr1 | 41,080,239 | 42 | Left | 1,059 | 0 |
| haplotype 2 | Chr2 | 31,426,022 | 26 | Both | 1,114 | 461 |
| | Chr3 | 35,402,754 | 36 | Right | 0 | 752 |
| | Chr4 | 25,485,865 | 22 | Left | 1,143 | 0 |
| | Chr5 | 28,881,267 | 34 | Both | 761 | 686 |
| | Chr6 | 28,233,716 | 22 | Neither | 0 | 0 |
| | Chr7 | 23,620,980 | 23 | Neither | 0 | 0 |
| | Chr8 | 20,646,017 | 20 | Both | 748 | 559 |
| | Chr9 | 27,227,112 | 35 | Left | 697 | 0 |
| | Chr10 | 12,256,734 | 23 | Neither | 0 | 0 |

**Table SM8. Primers used for PCR amplification.**

| Primer | Sequence (5' to 3') |
|---|---|
| *BLH9*_F | TCATCAAGCATGGTGACGAGC |
| *BLH9*_R | AGGATATGAAGGCGAGGCAC |
| *HSP90*_F | GGAAACAGAGAGAAAACGG |
| *HSP90*_R | AAACAACAACAACCAGAGCG |
| *Xdh*_F | CCAATTATTGATGCATTCCG |
| *Xdh*_R | CTGTCAATCTAACAGATGCC |

**Table SM9. TALE superfamily members in *A. thaliana* obtained from TAIR.**

| Gene | TAIR Accession | Protein name | Protein length (aa) |
|---|---|---|---|
| *ATH1* | Locus:2005494 | AT4G32980.1 | 473 |
| *BEL1* | Locus:2177856 | AT5G41410.1 | 611 |
| *BLH1* | Locus:2039250 | AT2G35940.1 | 680 |
| *BLH2* | Locus:2115000 | AT4G36870.1 | 739 |
| *BLH3* | Locus:2018398 | AT1G75410.1 | 524 |
| *BLH4* | Locus:2049035 | AT2G23760.4 | 679 |
| *BLH5* | Locus:2039605 | AT2G27220.2 | 449 |
| *BLH6* | Locus:2139614 | AT4G34610.1 | 532 |
| *BLH7* | Locus:2042609 | AT2G16400.1 | 482 |
| *BLH8* | Locus:2057856 | AT2G27990.1 | 584 |
| *BLH9* | Locus:2185183 | AT5G02030.1 | 575 |
| *BLH10* | Locus:2013154 | AT1G19700.1 | 538 |
| *BLH11* | Locus:2018457 | AT1G75430.1 | 290 |
| *KNATM* | Locus:2006782 | AT1G14760.1 | 142 |
| *STM* | Locus:2027089 | AT1G62360.1 | 382 |
| *KNAT1* | Locus:2128828 | AT4G08150.1 | 398 |
| *KNAT2* | Locus:2026810 | AT1G70510.2 | 314 |
| *KNAT3* | Locus:2146945 | AT5G25220.1 | 431 |
| *KNAT4* | Locus:2184911 | AT5G11060.1 | 393 |
| *KNAT5* | Locus:2116632 | AT4G32040.1 | 383 |

| | | | |
|---|---|---|---|
| *KNAT6* | Locus:2028075 | AT1G23380.2 | 329 |
| *KNAT7* | Locus:2015554 | AT1G62990.1 | 291 |

**Table SM10. Primers used for overexpression of *BLH9* and *AtBLH9* in *Arabidopsis thaliana.***

| Primer | Sequence (5' to 3') | Description |
|---|---|---|
| BLH9_BamHI_IF_F | GAGGCCTACGGGGATCCATGTCTTCCGAGGTCGGAGGATA | In-Fusion cloning |
| BLH9_EcoRI_IF_R | CGGGGTACCCGGAATTCTTATTCACCAACACCGAGGCCCAA | In-Fusion cloning |
| AtBLH9_BamHI_IF_F | GAGGCCTACGGGGATCCATGGCTGATGCATACGAGCCTTATC | In-Fusion cloning |
| AtBLH9_EcoRI_IF_R | CGGGGTACCCGGAATTCTCAACCTACAAAATCATGTAGAAACTG | In-Fusion cloning |
| pBICP_35S_F | GATGTGATATCTCCACTGACG | Confirmation of inserted sequence |
| pBICP_35S_R | CTTATCTGGGAACTACTCACA | Confirmation of inserted sequence |
| BLH9_seq_F | GGCGATGCAGGACGTGAATG | Confirmation of inserted sequence |
| AtBLH9_seq_F | GAATGCTATAACGGACCAGC | Confirmation of inserted sequence |
| UBC_qPCR_F | CTGCGACTCAGGGAATCTTCTAA | qPCR |
| UBC_qPCR_R | TTGTGCCATTGAATTGAACCC | qPCR |
| BLH9_qPCR_F | GCTCCTACTTCTCGCTAAACCC | qPCR |
| BLH9_qPCR_R | GGGGAGGGCGTGGATGTGT | qPCR |
| AtBLH9_qPCR_F | CCTAGCTACAGTAATTTCATGGG | qPCR |
| AtBLH9_qPCR_R | CGGCGGCGTTGGCTTCACC | qPCR |
